# Centriole structural integrity defects are a crucial feature of Hydrolethalus Syndrome

**DOI:** 10.1101/2024.03.06.583733

**Authors:** Ana Curinha, Zhaoyu Huang, Taylor Anglen, Margaret A. Strong, Colin R. Gliech, Cayla E. Jewett, Anoek Friskes, Andrew J. Holland

**Affiliations:** Department of Molecular Biology and Genetics, Johns Hopkins University School of Medicine, Baltimore, MD, USA; Department of Biomedical Engineering, Duke University, Durham, NC, USA; Division of Cell Biology, Oncode Institute, The Netherlands Cancer Institute, Amsterdam, The Netherlands

## Abstract

Hydrolethalus Syndrome (HLS) is a lethal, autosomal recessive ciliopathy caused by the mutation of the conserved centriole protein HYLS1. However, how HYLS1 facilitates the centriole-based templating of cilia is poorly understood. Here, we show that mice harboring the HYLS1 disease mutation die shortly after birth and exhibit developmental defects that recapitulate several manifestations of the human disease. These phenotypes arise from tissue-specific defects in cilia assembly and function caused by a loss of centriole integrity. We show that HYLS1 is recruited to the centriole by CEP120 and functions to recruit centriole inner scaffold proteins that stabilize the centriolar microtubule wall. The HLS mutation disrupts the interaction of HYLS1 with CEP120 leading to HYLS1 displacement and degeneration of the centriole distal end. We propose that tissue-specific defects in centriole integrity caused by the HYLS1 mutation prevent ciliogenesis and drive HLS phenotypes.

## INTRODUCTION

Centrioles are barrel-shaped cellular organelles composed of a nine-fold triplet arrangement of microtubules with three important functions. First, centrioles form the core of the centrosome, a microtubule organizing center that nucleates the interphase microtubule cytoskeleton and guides the assembly of the mitotic spindle apparatus during cell division (Gönczy, 2012; Nigg & Holland, 2018). Second, centrioles function as basal bodies that dock at the plasma membrane and initiate the biogenesis of cilia that have motile or sensory functions (Breslow & Holland, 2019; Nigg & Raff, 2009). Finally, recent work revealed centrioles function as signaling centers that restrict the proliferation of polyploid cells in mammals (Sladky *et al*., 2020, 2022). Centrioles are remarkably stable structures that persist over multiple cell cycles with minimal turnover of their molecular components. Understanding how centrioles are assembled and maintained remains an important challenge.

In proliferating cells, centriole biogenesis is tightly coupled to cell cycle progression (Holland *et al*., 2010; Guderian *et al*., 2010; Cunha-Ferreira *et al*., 2013). G1 cells contain a pair of parent centrioles that differ in age and structure, with only the oldest parent centriole possessing distal appendages required to nucleate cilia (Nigg & Stearns, 2011; Tanos *et al*., 2013). Centriole duplication starts at the onset of S-phase, when a new procentriole is assembled orthogonally at the base of each parent centriole. Procentrioles elongate during the S and G2 phase but are maintained in an immature state that is unable to duplicate. In mitosis, procentrioles separate from parent centrioles and undergo maturation to become replication-competent parent centrioles (Azimzadeh & Marshall, 2010; Gönczy, 2012; Nigg & Holland, 2018). At the same time, the mitotic spindle evenly partitions centriole pairs into daughter cells to maintain appropriate centriole numbers. Errors in centriole duplication and assembly have been linked to a variety of diseases, including cancer, growth retardation, and neurodevelopmental defects (Basto *et al*., 2008; Coelho *et al*., 2015; Serçin *et al*., 2016; Levine *et al*., 2017; Levine & Holland, 2018).

Mammalian centrioles can be divided into three main regions: a proximal end, which defines the centriolar 9-fold symmetry and is essential for centriole duplication; a central core, required for the cohesion of the centriolar microtubule triplets; and a distal end, required for basal body membrane docking and ciliogenesis (LeGuennec *et al*., 2021). Centriole duplication begins with the assembly of the cartwheel comprised of the proteins SAS-6, STIL, and CEP135, which link to the centriolar microtubule wall through the protein CPAP (Tang *et al*., 2009; Schmidt *et al*., 2009; Lin *et al*., 2013; Pelletier *et al*., 2006; Vásquez-Limeta *et al*., 2022). The establishment of the centriolar microtubule triplet begins with the formation of a single microtubule (the A-tubule) that attaches to the cartwheel, upon which the B- and C-tubules are assembled (Guichard *et al*., 2010; Li *et al*., 2012b; Guichard *et al*., 2013). Together with CPAP, CEP295 and CEP120 coordinate the recruitment of several proteins that are required for centriole elongation and structural integrity (Chang *et al*., 2016; Tsai *et al*., 2019). Accordingly, the depletion of CPAP leads to short or broken centrioles (Vásquez-Limeta *et al*., 2022; Zheng *et al*., 2016), while the depletion of CEP120, CEP295, or their interacting factors leads to short centrioles that lack distal appendages and the ability to ciliate. By contrast, overexpression of CPAP, CEP120, or CEP295 leads to over-elongation of the centriolar microtubule wall (Lin *et al*., 2013; Chang *et al*., 2016; Comartin *et al*., 2013). Despite a broad understanding of the molecular players, how these proteins specifically cooperate to ensure centriole stability remains unclear.

A large spectrum of disorders, termed ciliopathies, arise from defects in the centriole-cilia apparatus. While complete loss of cilia function leads to developmental arrest and mid-gestation embryonic lethality, some ciliopathies are compatible with life and produce diverse phenotypes such as polydactyly, mental retardation, and infertility (Reiter & Leroux, 2017; Nigg & Raff, 2009). Hydrolethalus Syndrome (HLS) is a severe autosomal recessive ciliopathy characterized by developmental defects (polydactyly, hydrocephaly, and cleft lip or palate) that lead to lethality around birth. HLS is caused by a D211G point mutation in the conserved centriole protein HYLS1 and is enriched in the Finnish population (Mee *et al*., 2005; Honkala *et al*., 2009). HYLS1 is required to anchor basal bodies at the plasma membrane and assemble the transition zone required for cilia biogenesis and signaling, and recent work has implicated cilia dysfunction as the major cause of HLS defects (Dammermann *et al*., 2009; Wei *et al*., 2016; Serwas *et al*., 2017; Hou *et al*., 2020; Chen *et al*., 2021). HYLS1 has been shown to be dispensable for centriole duplication and centrosome function in *C. elegans,* suggesting a basal body specific function (Dammermann *et al*., 2009; Serwas *et al*., 2017). However, we previously identified HYLS1 in a genome-wide screen for genes required to arrest cell proliferation upon centriole amplification (Evans *et al*., 2021), suggesting that in contrast to *C. elegans*, HYLS1 may also function in centriole assembly in mammalian cells. Moreover, HYLS1 has been shown to be required for the elongation of the unusually long spermatocyte centrioles in flies (Hou *et al*., 2020). Ultimately, how the D211G HLS mutation impacts the critical role of HYLS1 in supporting cilia assembly is not understood.

In this study, we uncover a tissue-specific role of HYLS1 in centriole structural stability. Depletion of HYLS1 or knock-in of the HLS disease-causing mutation results in short or broken centrioles that replicate and maintain a normal centrosome content. Centriole structural abnormalities arise from reduced recruitment of mutant HYLS1 to the centriole leading to the impaired localization of inner scaffold centriolar proteins and defective assembly of centriole distal appendages. We show that CEP120 binds and recruits HYLS1 to the centriole, and that the HYLS1 D211G mutation impairs this interaction. Together, our data show that ciliogenesis defects in HLS are the consequence of a loss of centriole stability driven by mutant HYLS1.

## RESULTS

### Hyls1 D226G leads to tissue-specific ciliogenesis defects and perinatal lethality

To probe the underlying cause of HLS, we knocked-in the corresponding human *HYLS1 D211G* mutation into the murine *Hyls1* gene (*Hyls1 D226G*) (**Figure S1A**). Wildtype and heterozygous *Hyls1 D226G* mice (hereafter *Hyls1^+/+^* and *Hyls1^+/DG^*, respectively) were born at Mendelian ratios with no obvious phenotype, while homozygous *Hyls1^DG/DG^* mice died at birth or shortly thereafter (**Figure 1A**). *Hyls1^DG/DG^* embryos were similar in size and weight to *Hyls1^+/+^*and *Hyls1^+/DG^* embryos at embryonic day 14.5 (E14.5) but were smaller and lighter at E18.5 and at birth (**Figure S1B-D**), suggesting late developmental growth defects. *Hyls1^DG/DG^* pups displayed frequent polydactyly (**Figure 1B**), a cleft palate (**Figure 1C**), breathing abnormalities (data not shown), and a modest reduction in cerebral cortex size (**Figure S1E/F**). The kidney at E18.5 and P0 was also smaller in *Hyls1^DG/DG^* animals compared to *Hyls1^+/+^* and *Hyls1^+/DG^* animals (**Figure 1D**), and histological analysis revealed small cysts, dilated tubules, and increased fibrosis in *Hyls1^DG/DG^* kidneys (**Figure S1G**).

**Figure 1:**
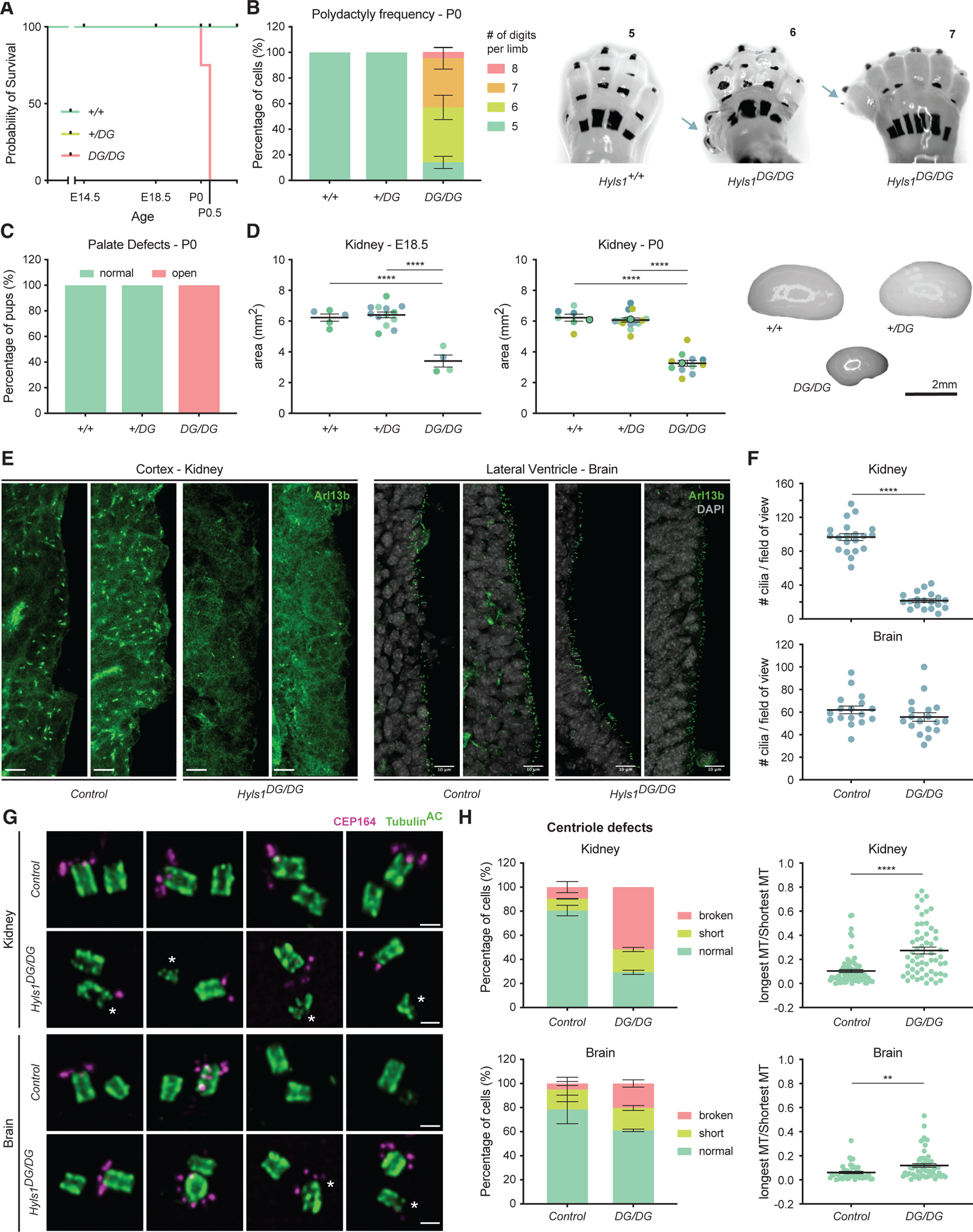
Hyls1 D226G leads to perinatal lethality. (A) Survival of *Hyls1^+/+^*, *Hyls1^+/DG^*, and *Hyls1^DG/DG^* animals. N≥4 mice per genotype. (B) Quantification of the number of digits per limb (left) and representative images (right) in *Hyls1^+/+^*, *Hyls1^+/DG^*, and *Hyls1^DG/DG^*P0 animals. N≥7 mice per genotype. (C) Percentage of pups with normal or open palate in *Hyls1^+/+^*, *Hyls1^+/DG^*, *Hyls1^DG/DG^* P0 animals. N≥3 mice per genotype. (D) Analysis of kidney area in E18.5 (left) and P0 (middle) animals and representative P0 images (right) in *Hyls1^+/+^*, *Hyls1^+/DG^*, and *Hyls1^DG/DG^*animals. Circled points indicate the data used in the representative images. N≥7 mice per genotype. (E) Representative images of immunofluorescence analysis of cilia (Arl13b) in the kidneys (left) and brain (right) in control *(Hyls1^+/+^* or *Hyls1^+/DG^)* and *Hyls1^DG/DG^* P0 animals. (F) Quantification of cilia (Arl13b) abundance in the kidneys (upper) and in the brain (lower) from immunofluorescence images of control *(Hyls1^+/+^* or *Hyls1^+/DG^)* and *Hyls1^DG/DG^* P0 animals. N=2 mice per genotype. (G) Representative images of centrioles (Tubulin^AC^) and distal appendages (CEP164) analyzed by U-ExM in the kidneys (upper) and in the brain (lower) of control *(Hyls1^+/+^* or *Hyls1^+/DG^)* and *Hyls1^DG/DG^* P0 animals. (H) Quantification analysis of centriole defects in the kidneys (upper) and in the brain (lower) in control *(Hyls1^+/+^* or *Hyls1^+/DG^)* and *Hyls1^DG/DG^* P0 animals analyzed by U-ExM. N=2 mice per genotype. Data are represented as mean ± SEM. Statistical significance was assessed using one-way-ANOVA with Tukey’s multiple comparisons test (D) and an unpaired two-tailed Student’s t-test with Welch’s correction (F/H). (**) P < 0.01, (****) P < 0.0001. Only significant results are indicated. Number of digits per limb and centriole defects analysis was assessed using two-way-ANOVA with post-hoc analysis and results are summarized in supplementary material. Scale bar is 10 µm in (E) and 250 nm in (G). Asterisk (*) indicates defective centrioles.

We next analyzed cilia assembly in the kidney and brains of P0 animals. Compared with *Hyls1^+/+^*and *Hyls1^+/DG^* control mice, the number of cilia was substantially reduced in the kidney but not in the brain of *Hyls1^DG/DG^* animals (**Figure 1E/F**). To examine centriole integrity, we performed ultrastructure expansion microscopy (U-ExM) in the kidneys, brains, and epidermis of control (*Hyls1^+/+^; Hyls1^+/DG^*) and *Hyls1^DG/DG^* mice. Consistent with their lack of cilia, we found that ∼70% of centrioles in the kidneys and epidermis from *Hyls1^DG/DG^*mice had structural defects, appearing short or broken (**Figure 1G/H, S1H and S2A/B**). These defects were less evident in the brains of *Hyls1^DG/DG^*animals, where only ∼40% of the centrioles appeared problematic (**Figure 1G/H and S1I**). By contrast, analysis of centrioles in brain multiciliated cells (MCCs) by U-ExM did not reveal obvious structural issues in *Hyls1^DG/DG^*mice (**Figure S2C/D**), and immunofluorescence analysis showed that multiciliogenesis seems to occur normally in these animals (**Figure S2E**). Together, our data show that the *Hyls1^DG/DG^* mouse model recapitulates key features of human Hydrolethalus Syndrome, which is driven by tissue-specific defects in cilia assembly.

### HYLS1 D226G/D211G drives ciliogenesis defects through loss of distal appendages

We next generated mouse embryonic fibroblasts (MEFs) from *Hyls1^+/+^, Hyls1^+/DG^* and *Hyls1^DG/DG^* animals to examine their centriole and cilia defects in more detail. Cells were serum-starved to induce ciliation and stained for centrosomes (γ-tubulin), distal appendages (CEP164), and cilia (acetylated α-tubulin). Interestingly, while centrosome number was maintained for all genotypes, we observed a dramatic reduction in the appearance of centriole distal appendages in *Hyls1^DG/DG^* MEFs, and cilia formation was nearly completely abolished in these cells (**Figure 2A/B**). The rare cilia observed in *Hyls1^DG/DG^*MEFs, however, were of normal length (**Figure S3A/B**) and fully capable of recruiting Smoothened in response to Smoothened agonist (SAG) treatment, suggesting intact Hedgehog signaling (**Figure S3A/C**). This suggests that HYLS1 is required for cilia formation but not cilia maintenance or function in MEFs.

**Figure 2:**
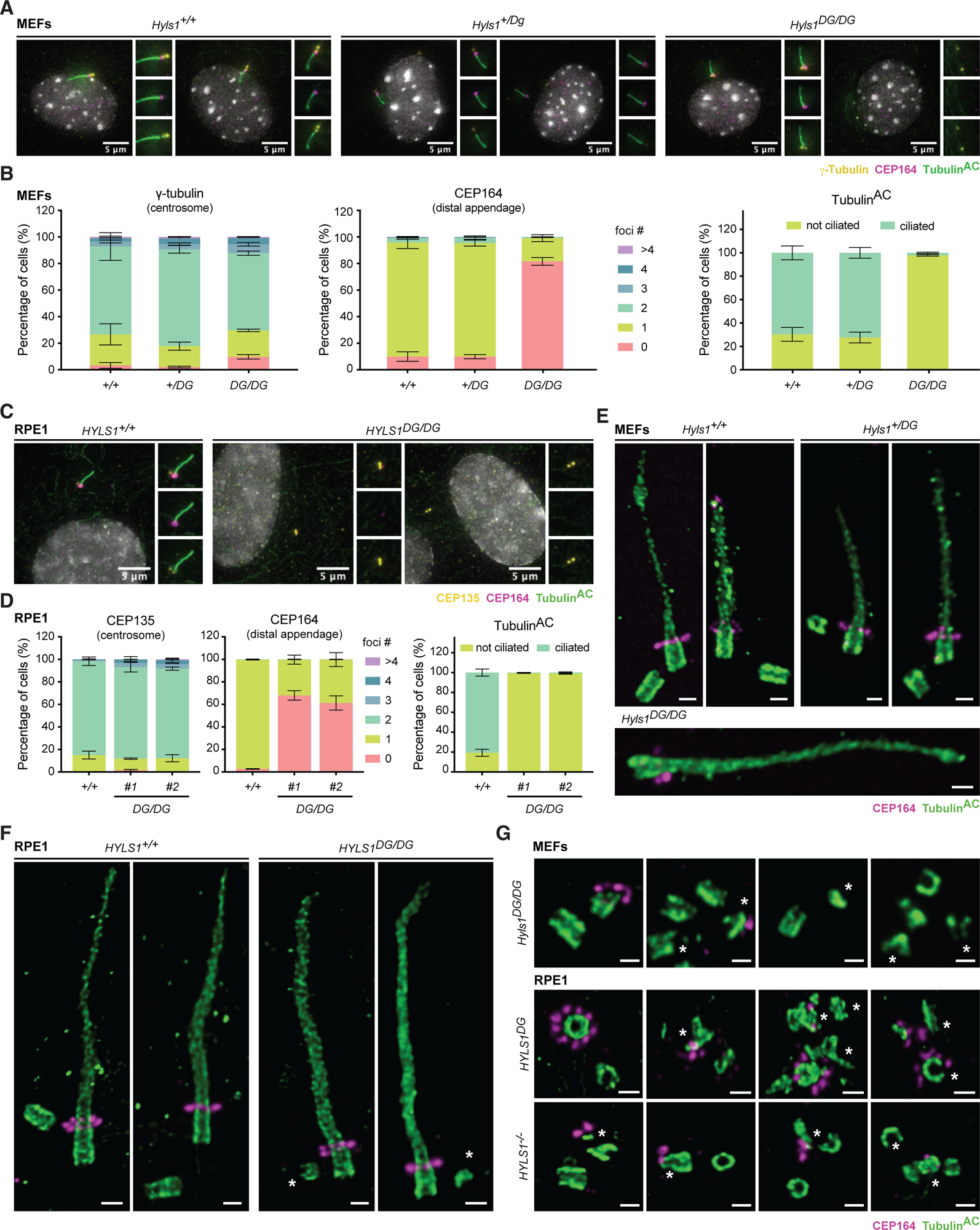
Hyls1 D226G and HYLS1 D211G impair distal appendages assembly. (A) Representative images of centrosomes (γ-Tubulin), distal appendages (CEP164) and cilia (Tubulin^AC^) in MEFs of *Hyls1^+/+^*, *Hyls1^+/DG^*, and *Hyls1^DG/DG^* animals. (B) Quantification of centrosomes (left), distal appendages (middle), and ciliation frequency (right) from immunofluorescence images of *Hyls1^+/+^*, *Hyls1^+/DG^*, and *Hyls1^DG/DG^* MEFs. MEFs generated across N≥3 mice per genotype were analyzed. (C) Representative images of centrosomes (CEP135), distal appendages (CEP164) and cilia (Tubulin^AC^) in *HYLS1^+/+^* and *HYLS1^DG/DG^*RPE1 cells. (D) Quantification of centrosomes (left), distal appendages (middle), and ciliation frequency (right) from immunofluorescence images of *HYLS1^+/+^* and *HYLS1^DG/DG^* RPE1 cells. Data across n=3 biological replicates were analyzed. (E) Representative images of centrioles and cilia (Tubulin^AC^), and distal appendages (CEP164) in MEFs of *Hyls1^+/+^*, *Hyls1^+/DG^*, and *Hyls1^DG/DG^* animals analyzed by U-ExM. (F) Representative images of centrioles and cilia (Tubulin^AC^) and distal appendages (CEP164) in *HYLS1^+/+^*and *HYLS1^DG/DG^* RPE1 cells analyzed by U-ExM. (G) Representative images of unstable centrioles (Tubulin^AC^) and impaired distal appendage assembly (CEP164) in *Hyls1^DG/DG^* MEFs and *HYLS1^DG/DG^* and *HYLS1^-/-^* RPE1 cells analyzed by U-ExM. Data are represented as mean ± SEM. Foci number and ciliogenesis analysis were assessed using two-way-ANOVA with post-hoc analysis and results are summarized in supplementary material. Scale bar is 250 nm in (E/F/G). Asterisk (*) indicates defective centrioles in (F/G).

Since multiciliated cells appeared unaffected in the brains of *Hyls1^DG/DG^* mice, we also examined cilia assembly during the differentiation of ependymal cell progenitors *in vitro*. Consistent with our observations *in vivo*, no clear differences were observed in the number of cilia assembled in *Hyls1^DG/DG^* MCCs (**Figure S3E/F**). Furthermore, analysis of MCC differentiation *in vitro* by U-ExM showed that HYLS1 D226G does not impact MCC differentiation (**Figure S4A**).

To assess whether distal appendage and cilia defects observed in *Hyls1^DG/DG^*MEFs were conserved in humans, we knocked in the D211G mutation into both *HYLS1* alleles in human RPE1 cells (hereafter *HYLS1^DG/DG^*). *HYLS1^DG/DG^* RPE1 cells were similarly defective in cilia assembly (**Figure 2C/D**), with only ∼1% of serum starved *HYLS1^DG/DG^* RPE1 cells forming cilia compared to 80% of *HYLS1^+/+^* cells. In addition, *HYLS1^DG/DG^*RPE1 cells exhibited the same lack of distal appendage formation as *Hyls1^DG/DG^*MEFs, despite maintaining normal centrosome counts (**Figure 2C/D**). Importantly, cell cycle analysis showed that the D211G mutation did not impact cell cycle progression (**Figure S3D**). To compare the impact of the HYLS1 D211G mutation with the complete loss of HYLS1, we generated HYLS1 knockout RPE1 cells (hereafter *HYLS1^-/-^*). Like HYLS1 mutant cells, loss of HYLS1 led to a severe defect in distal appendage formation and cilia assembly (**Figure S4B/C**). We conclude that *HYLS1^DG/DG^*is a loss of function protein mutation that is dispensable for centrosome maintenance but prevents centriole distal appendage formation and ciliation.

Finally, we performed U-ExM on serum-starved MEFs and RPE1 cells. In the rare instances where cilia were present, they displayed normal structure and localized CEP164 at their distal end (**Figure 2E/F and S4D**). In accordance with our data from mouse tissues and consistent with their inability to ciliate, *Hyls1^DG/DG^* MEFs, and *HYLS1^DG/DG^* and *HYLS1^-/-^* RPE1 cells lacking cilia showed evidence of short or disintegrating centrioles with defective recruitment of CEP164 (**Figure 2G**). Together, these data show that the HYLS1 D211G mutation disrupts centriole stability and impairs ciliogenesis in some cell types.

### HYLS1 is required for centriole structural integrity

To characterize the centriole integrity defect more fully, we performed U-ExM analysis in proliferating *Hyls1^DG/DG^*MEFs and *HYLS1^DG/DG^* and *HYLS1^-/-^* RPE1 cells. Centrioles from *Hyls1^+/+^* and *Hyls1^+/DG^* MEFs and *HYLS1^+/+^* RPE1 cells had a normal structure with a length of ∼500 nm and a width of ∼250 nm (**Figure 3A/B and S5A/B**). Consistent with our data in serum-starved cells, ∼57% of centrioles in *Hyls1^DG/DG^* MEFs and ∼55% of *HYLS1^DG/DG^*RPE1 cells displayed a break at the distal end of the centriole, and most of the remaining centrioles were short (< 400nm in length) (**Figure 3A/B and S5A/B**). Centrin2 and CEP164 analysis by U-ExM showed these proteins were detectable at some broken centrioles but lacked their characteristic localization in *HYLS1^DG/DG^* RPE1 cells (**Figure 3B**). These data suggest that HYLS1 is required for proper assembly and/or maintenance of the centriole distal end.

**Figure 3:**
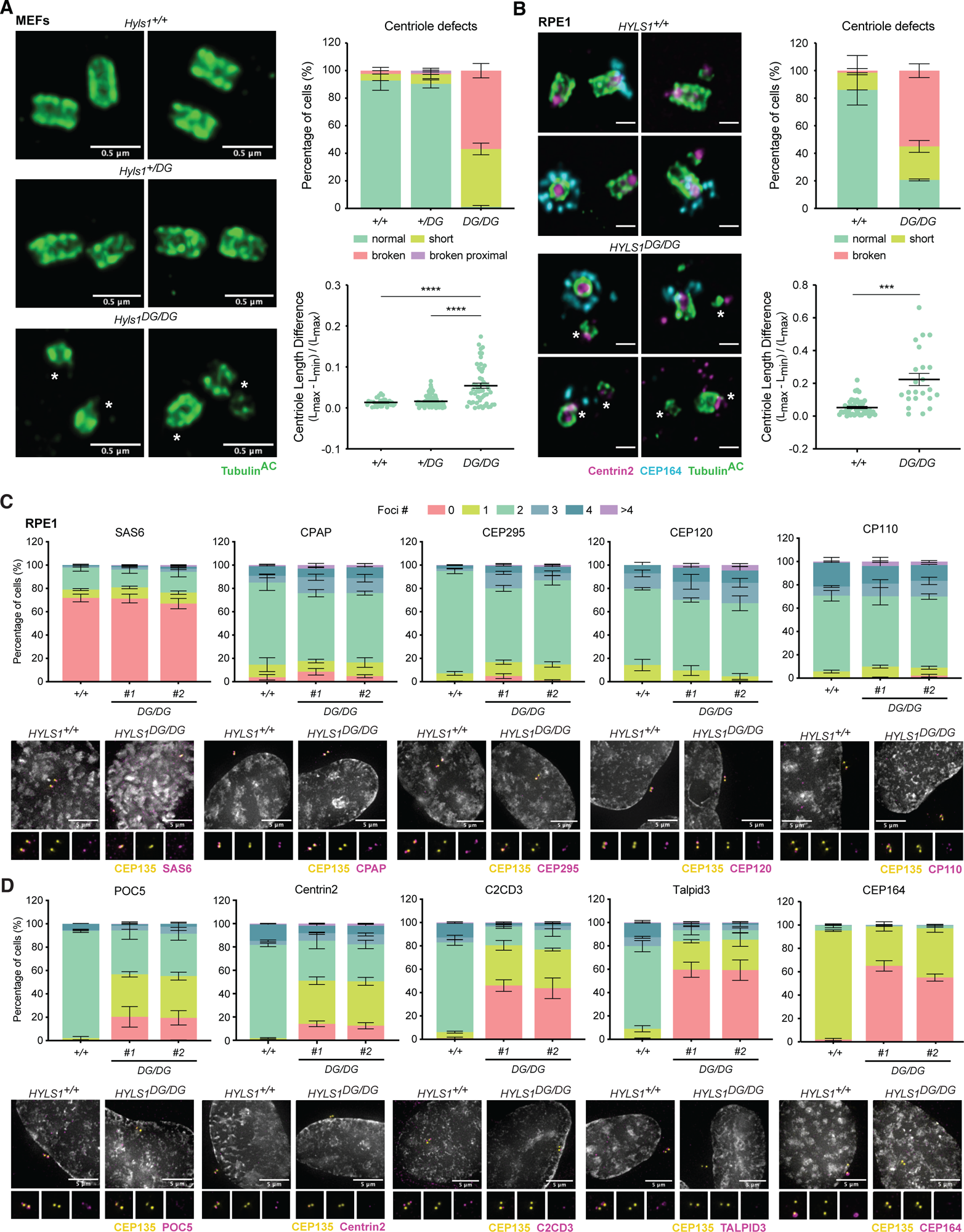
HYLS1 is essential for centriole structural integrity and distal end formation. (A) Representative U-ExM images (left) of centrioles (Tubulin^AC^), quantification of centriole defects (top right), and analysis of the relative length difference between the longest and shortest microtubule in each centriole (bottom right) in MEFs of *Hyls1^+/+^*, *Hyls1^+/DG^*, and *Hyls1^DG/DG^* animals. MEFs generated across N≥2 mice per genotype were analyzed. (B) Representative U-ExM images (left) of centrioles (Tubulin^AC^), Centrin2, and distal appendages (CEP164), quantification of centriole defects (top right), and analysis of the relative length difference between the longest and shortest microtubule in each centriole (bottom right) of *HYLS1^+/+^*and *HYLS1^DG/DG^* RPE1 cells. Data from n=2 biological replicates were analyzed. (C) Quantification (upper) and representative immunofluorescence images (lower) of centriole proximal markers (SAS-6) and centriole elongation and structural integrity factors (CPAP, CEP295, CEP120, and CP110) in *HYLS1^+/+^* and *HYLS1^DG/DG^* RPE1 cells. Data from n≥3 biological replicates were analyzed. (D) Quantification (upper) and representative immunofluorescence images (lower) of inner scaffold proteins (POC5 and Centrin2), distal end proteins (C2CD3 and Talpid3), and distal appendages (CEP164) in *HYLS1^+/+^* and *HYLS1^DG/DG^* RPE1 cells. Data from n≥3 biological replicates were analyzed. Data are represented as mean ± SEM. Statistical significance was assessed using one-way-ANOVA with Tukey’s multiple comparisons test (A – lower) and an unpaired two-tailed Student’s t-test with Welch’s correction (B – lower). (***) P < 0.001, (****) P < 0.0001. Only significant results are indicated. Centriole defects or foci number analysis was assessed using two-way-ANOVA with post-hoc analysis and results are summarized in supplementary material. Scale bar is 250 nm in (B). Asterisk (*) indicates defective centrioles.

Given the short and broken centrioles observed with HYLS1 deficiency, we speculated that HYLS1 may impact the localization of proteins involved in the early steps of centriole assembly and elongation. To better understand at what stage of centriole assembly is HYLS1 required, we analyzed the localization of proximal, central, or distal end centriole proteins in RPE1 cells. Similar to what we observed for CEP135 (**Figure 2D and S5C**), the proximal end cartwheel component SAS-6 was unaffected by HYLS1 D211G mutation (**Figure 3C**) or HYLS1 loss (**Figure S5E**). Analysis of proteins involved in centriole elongation and stability showed that neither *HYLS1* loss nor *HYLS1* mutation affected the localization of CPAP, CEP295, CEP120, or CP110 to centrioles in *HYLS1^DG/DG^* RPE1 cells and *HYLS1^-/-^* RPE1 cells (**Figure 3C and S5E**). By contrast, recruitment of the inner scaffold proteins POC5 and Centrin2 was diminished in *HYLS1^DG/DG^* and *HYLS1^-/-^* RPE1 cells (**Figure 3D and S5E**). Similarly, recruitment of the centriole distal-end proteins Talpid3 and C2CD3, as well as the distal appendage protein CEP164, was also diminished by HYLS1 D211G mutation or HYLS1 loss (**Figure 3D and S5E**). These data suggest that the centriole defects observed upon HYLS1 loss or mutation may result from defective recruitment of inner scaffold proteins, leading to the disintegration of the centriole microtubule wall.

### HYLS1 overexpression drives assembly of centriolar microtubules

The centriole phenotype caused by HYLS1 D211G is reminiscent of phenotypes observed with loss of known elongation factors CPAP, CEP120, and CEP295. We therefore tested if overexpression of HYLS1 would lead to abnormally long centrioles. We generated stable cell lines carrying a doxycycline-inducible WT or D211G HYLS1-mNeonGreen transgene (hereafter HYLS1^WT^-mNG or HYLS1^DG^-mNG) in *HYLS1^-/-^* DLD1 cells. Consistent with what was observed in RPE1 cells, HYLS1 loss led to impaired recruitment of inner scaffold, distal end, and distal appendage proteins in DLD1 cells (**Figure 4A-D and S6A/B**). Expression of HYLS1^WT^-mNG restored the localization of Centrin2 and CEP164 at 2 and 4 days after doxycycline addition (**Figure 4A-D**). Furthermore, analysis by U-ExM confirmed that most centrioles elongated normally and were structurally intact at these time points **(Figure 4E-H and S6C**). By contrast, expression of HYLS1^DG^-mNG was unable to effectively rescue Centrin2 and CEP164 localization (**Figure 4A-D**), and most centrioles remained broken (**Figure 4E-H and S6C**). Overexpression of HYLS1^WT^-mNG drove an increase in centriole length as well as centrioles harboring an extended microtubule at the distal end (**Figure 4E-G and S6C**), a phenotype that was rarely observed in cells overexpressing HYLS1^DG^-mNG. CP110 localized to the distal end of both broken and intact centrioles in all the genotypes analyzed **(Figure 4E)**.

**Figure 4:**
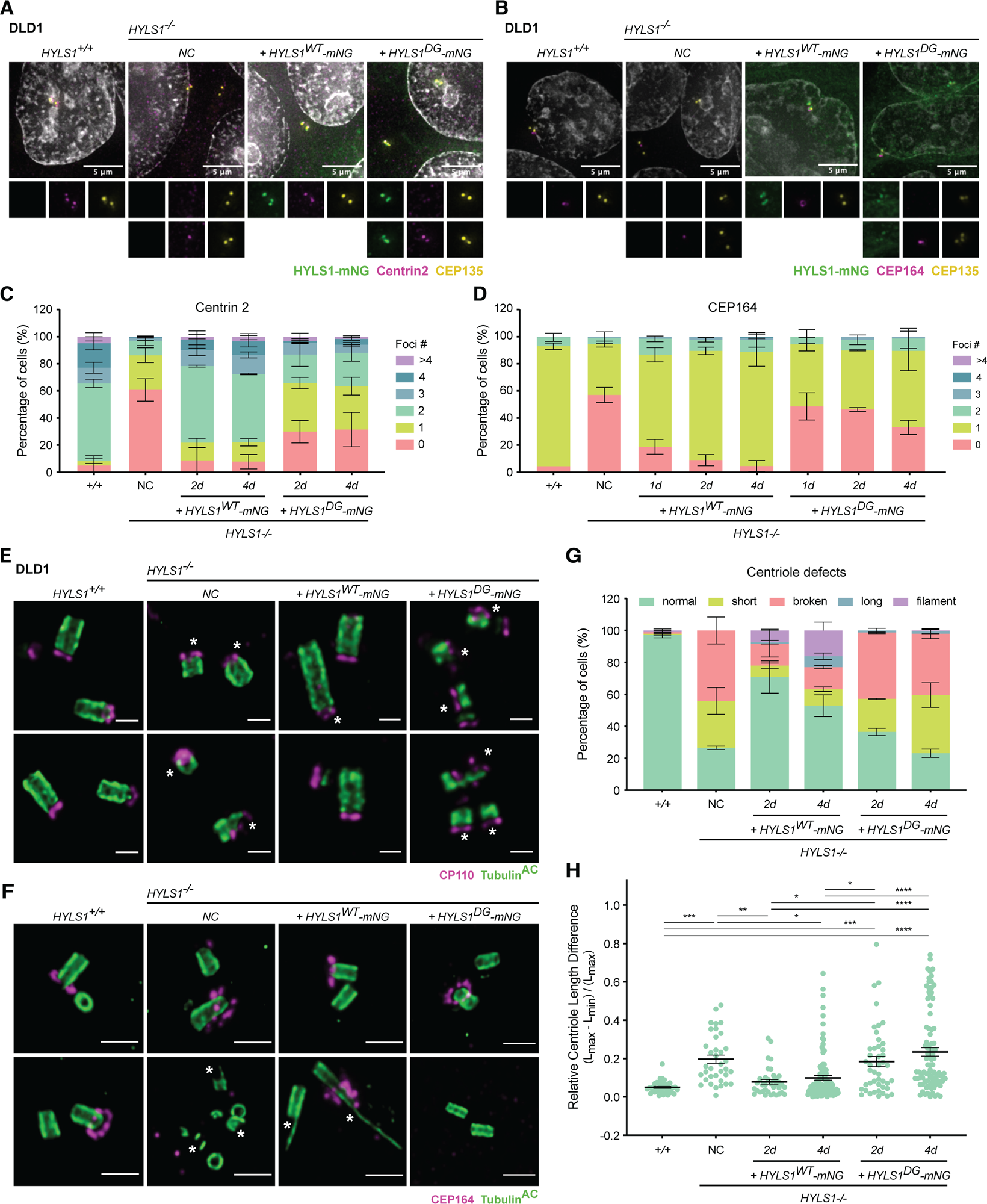
HYLS1 overexpression leads to over-elongated centrioles. (A) Representative immunofluorescence images of HYLS1-mNG, centrioles (CEP135), and an inner scaffold protein (Centrin2) in *HYLS1^+/+^,* and *HYLS1^-/-^* DLD1 cells with HYLS1-mNG (WT or D211G) add-back. (B) Representative immunofluorescence images of HYLS1-mNG, centrioles (CEP135), and distal appendages (CEP164) in *HYLS1^+/+^* and *HYLS1^-/-^* DLD1 cells with HYLS1-mNG (WT or D211G) add-back. (C) Quantification of inner scaffold protein (Centrin2) at centrioles (CEP135) from immunofluorescence images of *HYLS1^+/+^* and *HYLS1^-/-^*DLD1 cells with HYLS1-mNG (WT or D211G) add-back. Data from n=3 biological replicates were analyzed. (D) Quantification of distal appendages (CEP164) at centrioles (CEP135) from immunofluorescence images of *HYLS1^+/+^* and *HYLS1^-/-^*DLD1 cells with HYLS1-mNG (WT or D211G) add-back. Data from n=3 biological replicates were analyzed. (E) Representative images of centrioles (Tubulin^AC^), and centriolar cap protein (CP110) in *HYLS1^+/+^* and *HYLS1^-/-^*DLD1 cells with HYLS1-mNG (WT or D211G) add-back analyzed by U-ExM. (F) Representative images of centrioles (Tubulin^AC^), and distal appendages (CEP164) in *HYLS1^+/+^*and *HYLS1^-/-^* DLD1 cells with HYLS1-mNG (WT or D211G) add-back analyzed by U-ExM. (G) Quantification of centriole defects in *HYLS1^+/+^* and *HYLS1^-/-^* DLD1 cells with HYLS1-mNG (WT or D211G) add-back analyzed by U-ExM. Data from n=3 biological replicates were analyzed. (H) Analysis of the relative length difference between the longest and shortest microtubule in each centriole of *HYLS1^+/+^* and *HYLS1^-/-^* DLD1 cells with HYLS1-mNG (WT or D211G) add-back analyzed by U-ExM. Data from n=3 biological replicates were analyzed. Data are represented as mean ± SEM. Statistical significance was assessed using one-way-ANOVA with Tukey’s multiple comparisons test (H). (*) P < 0.05, (**) P < 0,01, (***) P < 0.001, (****) P < 0.0001. Only significant results are indicated. Foci number and centriole defects analysis were assessed using two-way-ANOVA with post-hoc analysis and results are summarized in supplementary material. Scale bar is 250 nm in (E) and 500 nm in (F). Asterisk (*) indicates defective centrioles.

To define whether the defects observed in *HYLS1^DG/DG^*cells were entirely driven by instability in the centriolar microtubule wall, we attempted to rescue centriole structure with the microtubule-stabilizing agent Taxol. Previous literature suggested that structurally weak centrioles are prone to further destabilize in mitosis (Wang *et al*., 2017; Schweizer *et al*., 2021). Therefore, we Taxol treated G2-phase *HYLS1^+/+^* and *HYLS1^DG/DG^* RPE1 cells, arrested them in mitosis overnight, and analyzed centriole integrity in the following G1-phase after mitotic release with the CDK1 inhibitor RO3306 (**Figure S6D**). Taxol treatment decreased the percentage of broken centrioles and induced the over-elongation of non-acetylated centriolar microtubules observed by U-ExM analysis in *HYLS1^DG/DG^*cells (**Figure S6E/F**). Interestingly, immunofluorescence analysis revealed that Taxol treatment did not rescue Talpid3 recruitment to the centriole and may only partially rescued the recruitment of Centrin2 **(Figure S6G**). As expected, CEP164 recruitment was unaffected by Taxol treatment since distal appendages would not be expected to be recruited until the following mitosis (**Figure S6G**). These data suggest that HYLS1 functions to stabilize the centriole microtubule wall to maintain the structural integrity of centrioles.

### HYLS1 D211G shows defective centriole recruitment

Since HYLS1 D211G behaves similarly to a complete loss of HYLS1 protein, we wondered whether HYLS1 localization to the centriole was affected by the HLS mutation. Since commercial HYLS1 antibodies exhibited poor centriole staining, we knocked-in a HA-tag onto the C-terminus of *HYLS1* in *HYLS1^+/+^* and *HYLS1^DG/DG^* RPE1 cells. HYLS1^WT^-HA was absent on G1 centrioles that lacked the procentriole marker SAS-6 but increased in abundance as cells duplicated centrioles in S phase and progressed through the cell cycle (**Figure 5A/B**). Importantly, HYLS1 was undetectable at basal bodies in serum-starved ciliated cells, arguing against a direct role for HYLS1 in cilia maintenance in human cells (**Figure 5C**). In contrast to HYLS1^WT^-HA, HYLS1^DG^-HA was undetectable at centrioles throughout the cell cycle, suggesting that HYLS1 D211 is critical for recruitment of HYLS1 to the centriole (**Figure 5A/B**).

**Figure 5:**
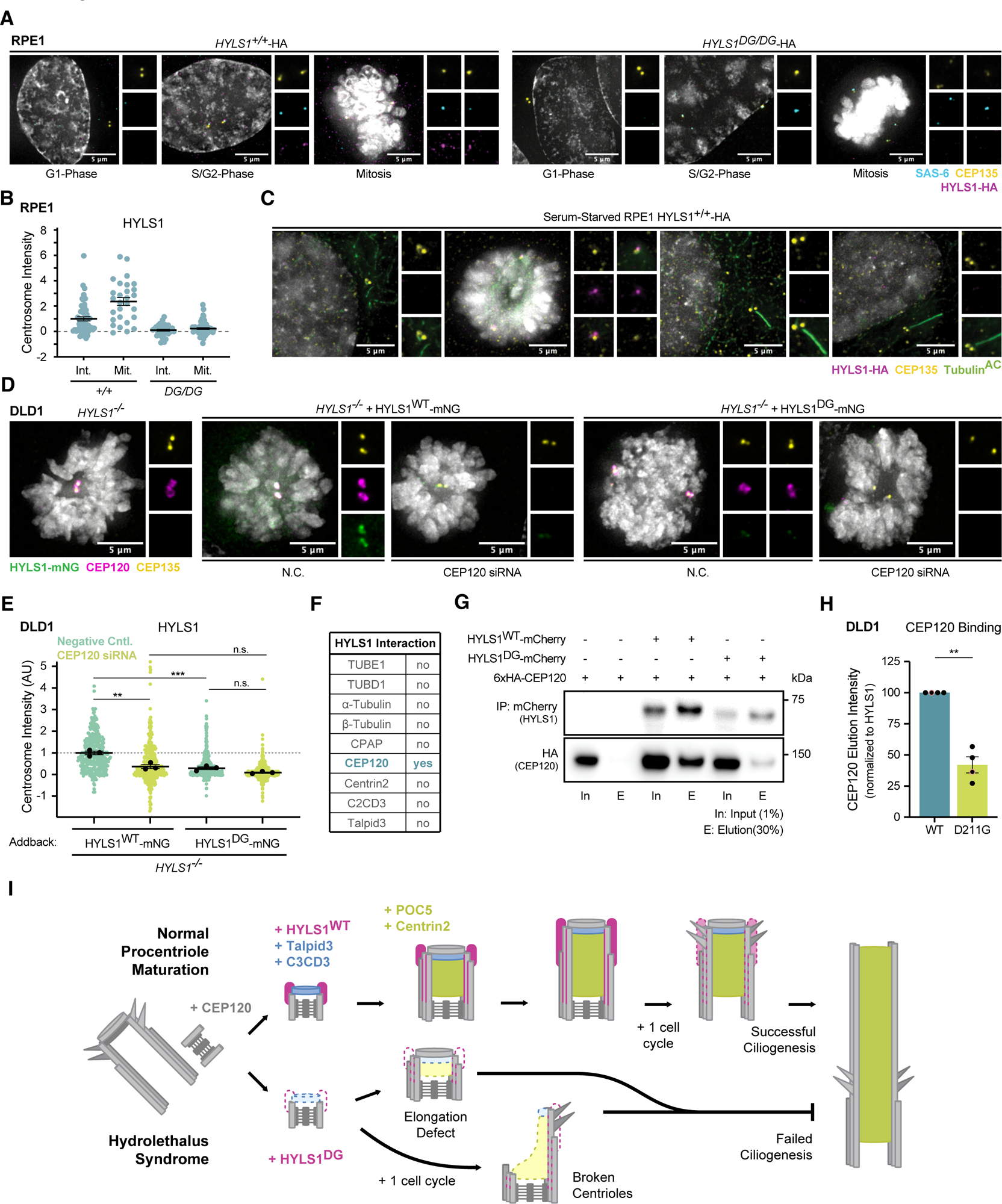
The HYLS1 D211G mutation weakens the interaction with CEP120. (A) Representative images of HYLS1 localization at the centrioles. Immunofluorescence analysis of centrioles (CEP135), centriole duplication marker (SAS-6), and HYLS1-HA in *HYLS1^+/+^*-HA and *HYLS1^DG/DG^*-HA unsynchronized RPE1 cells. (B) Quantification of immunofluorescence analysis of HYLS1-HA abundance at centrioles of interphase or mitotic *HYLS1^+/+^*-HA and *HYLS1^DG/DG^*-HA RPE1 cells. (C) Representative immunofluorescence images of HYLS1-HA, centrioles (CEP135), and cilia (Tubulin^AC^) in serum-starved *HYLS1^+/+^*-HA RPE1 cells. (D) Representative immunofluorescence images of HYLS1-mNG at the centriole (CEP135) upon CEP120 depletion by siRNA in *HYLS1^-/-^* DLD1 cells with HYLS1-mNG (WT or D211G) add-back. (E) Quantification of HYLS1-mNG intensity at the centriole (CEP135) upon CEP120 depletion by siRNA in *HYLS1^-/-^* DLD1 cells with HYLS1-mNG (WT or D211G) add-back. Data from n=3 biological replicates were analyzed. (F) Summary of HYLS1 WT co-immunoprecipitation experiments for all proteins analyzed. (G) Representative co-immunoprecipitation of 6xHA-CEP120 with HYLS1^WT^-mCherry or HYLS1^DG^-mCherry in *HYLS1^-/-^*DLD1 cells. (H) Quantification of co-immunoprecipitation experiments of 6xHA-CEP120 with HYLS1^WT^-mCherry or HYLS1^DG^-mCherry in *HYLS1^-/-^* DLD1 cells. Data from n=4 biological replicates were analyzed. (I) A model for how impaired recruitment of HYLS1 D211G to the centriole and subsequent decrease in the recruitment of inner scaffold and centriole distal end proteins cause the cilia defects underlying Hydrolethalus Syndrome. Data are represented as mean ± SEM. Statistical significance was assessed using one-way-ANOVA with Tukey’s multiple comparisons test of mean values from each replicate (D) and an unpaired two-tailed Student’s t-test (G). (**) P < 0.01, (***) P < 0.001.

High-resolution analysis of RPE1 *HYLS1^WT^-HA* cells with U-ExM showed that HYLS1 was more abundant at younger compared to older parent centrioles and was barely detectable at the basal body of ciliated cells (**Figure S7A/B**). In addition, HYLS1^DG^-HA was undetectable at centrioles, confirming our immunofluorescence analysis (**Figure S7A**). To examine the localization of HYLS1 D211G in cells with normal centrioles, we analyzed *HYLS1^+/+^*DLD1 cells carrying a doxycycline-inducible WT or D211G HYLS1-HA transgene (hereafter HYLS1^WT^-HA or HYLS1^DG^-HA). Consistent with what was observed in RPE1 cells, HYLS1^WT^-HA localized asymmetrically to the two parent centrioles in G1 cells and was recruited to the microtubule wall during early procentriole assembly before acetylation of the centriole microtubules (**Figure S7C**). In the growing procentrioles, HYLS1 was localized as a cap-like structure around the microtubule wall as the microtubules become acetylated (**Figure S7C**). Interestingly, the presence of endogenous wildtype HYLS1 partially enabled HYLS1^DG^-HA to localize to the centrioles of DLD1 cells, but the cap-like structure around the microtubule wall was absent (**Figure S7C**). Together, these data show that the D211G mutation prevents the effective recruitment of HYLS1 to the centriole microtubule wall during the early stages of procentriole assembly.

### The D211G HYLS1 mutation disrupts binding of HYLS1 to CEP120

We next set out to establish how the HYLS1 D211G mutation impacts its recruitment to the centriole. Our data suggests that, like CEP120 (Tsai *et al*., 2019), HYLS1 is enriched at the younger parent centriole and required for C2CD3 and Talpid3 recruitment to the centriole. Our immunofluorescence analysis, however, showed that HYLS1 mutation (**Figure 3C**) or HYLS1 depletion (**Figure S5E and S6A**) did not impair CEP120 recruitment to the centriole. Therefore, we tested whether CEP120 may also be responsible for HYLS1 recruitment to the centriole. CEP120 depletion by siRNA led to impaired centriole recruitment of HYLS1^WT^-mNG to similar levels as HYLS1^DG^-mNG (**Figure 5E and S7D/E**). These data suggest that CEP120 recruits HYLS1 to the centriole, which might promote the recruitment of Talpid3 and C2CD3. Finally, we tested whether HYLS1 interacts with proteins involved in centriole elongation and/or structural integrity by co-immunoprecipitation analysis in DLD1 HYLS1 add-back cell lines. Of the nine protein interactors we analyzed (**Figure 5F**), we observed a clear interaction between HYLS1 and CEP120, which was diminished by the HYLS1 D211G mutation (**Figure 5G/H**). These data show the HYLS1 D211G mutation disrupts its interaction with CEP120, leading to defects in HYLS1 recruitment to the centriole and the downstream recruitment of proteins involved in centriole structural integrity and distal end assembly.

## DISCUSSION

Despite recent studies aimed at understanding the function of HYLS1, how HYLS1 mutation impairs ciliogenesis and leads to HLS remains poorly understood. Using genetically engineered mice and cell lines, we identify centriole structural instability as a key feature responsible for the ciliogenesis defects caused by HYLS1 mutation. We show that the HYLS1 D211G disease mutation reduces recruitment of HYLS1 to the centriole, which in turn drives degeneration of the centriole distal end that harbors distal appendages required to initiate ciliogenesis. Strikingly, the defective centrioles lacking HYLS1 duplicate and maintain a normal centrosome number. Thus, we identify a new role of HYLS1 in maintaining centriole stability, offering mechanistic insight into how HYLS1 mutation leads to defects in ciliogenesis.

Analysis of primary murine and cultured human cells confirmed that, as was seen in other organisms, HYLS1 deficiency led to a ciliation defect (Dammermann *et al*., 2009; Wei *et al*., 2016; Hou *et al*., 2020; Chen *et al*., 2021; Serwas *et al*., 2017). Importantly, U-ExM analysis revealed that HYLS1 mutation or HYLS1 loss leads to centrioles that are short or broken at their distal end, consistent with a weakened microtubule wall. Similar structural integrity phenotypes have been described upon loss of the centriole microtubule wall protein WDR90 and the centriole inner scaffold proteins HAUS6 and CCDC15 (Steib *et al*., 2020; Schweizer *et al*., 2021; Arslanhan *et al*., 2023). Like HYLS1, WDR90 depletion has also been shown to impair the recruitment of the inner scaffold proteins POC5 and Centrin2 (Steib *et al*., 2020). Thus, our data indicate that the ciliogenesis defects underlying the clinical features of HLS result from a structural weakness of the centriole microtube wall driven by dysfunctional HYLS1 protein. However, defects in centriole integrity and cilia assembly were tissue-specific in the HLS mouse model and were not evident in the brains of *Hyls1^DG/DG^* animals. We speculate that a variable dependency on HYLS1 for centriole integrity or the presence of centriolar proteins with overlapping functions may explain this tissue specificity. Complete absence of cilia is lethal embryonically, suggesting that the tissue-specific defects in ciliogenesis caused by HYLS1 mutation produces more subtle phenotypes that support embryonic development in HLS patients.

HYLS-1 was shown to be required to initiate ciliogenesis but dispensable for centriole function and embryonic viability in *C. elegans* (Dammermann *et al*., 2009; Serwas *et al*., 2017). Unlike in mammals however, *C. elegans* centrioles degenerate during ciliogenesis, indicating they are dispensable for cilia maturation and maintenance (Dammermann *et al*., 2009; Serwas *et al*., 2017). *C. elegans* centrioles also have key structural differences such as a shorter length (∼175 nm), a centriole wall composed of microtubule singlets rather than triplets, and a lack of clear distal appendages that are required for ciliation in mammalian systems (Nigg & Holland, 2018; Blanco-Ameijeiras *et al*., 2022; Tanos *et al*., 2013). These key differences may underlie our observations of an expanded role for HYLS1 in the stability and elongation of the mammalian centriole.

The centriole integrity defects observed in HYLS1 mutant cells were partly rescued by microtubule stabilization with Taxol. In addition, HYLS1 overexpression led to over-elongated centriolar microtubules, possibly explaining the requirement of HYLS1 for the elongation of the giant centriole/basal body in fly spermatocytes (Hou *et al*., 2020). Similar phenotypes have been described for CPAP, CEP120, CEP295, or POC5 over-expression (Lin *et al*., 2013; Comartin *et al*., 2013; Kohlmaier *et al*., 2009; Schmidt *et al*., 2009; Chang *et al*., 2016; Schweizer *et al*., 2021), suggesting that HYLS1 might cooperate with these proteins to regulate the early steps of centriole assembly. The impaired localization of the distal end proteins and distal appendages observed in HYLS1 deficient cells likely arises because of the structural instability of the centriole and/or defective elongation of centriolar microtubules. This is consistent with the fact that Centrin2 and CEP164 were still observed at broken centrioles despite an overall impairment in their recruitment. Interestingly, direct microtubule stabilization with Taxol did not rescue the recruitment of Talpid3 and only partially rescued Centrin2 abundance at the centriole, raising the possibility that HYLS1 may play a direct role in recruiting these proteins. In the future, it will be important to establish if the phenotypes observed upon HYLS1 loss or mutation are a result of the impaired recruitment of inner scaffold proteins or due to a direct role of HYLS1 in fortifying the centriole microtubule wall.

CPAP, CEP120, and CEP295 bind microtubules and have also been shown to regulate centriole length and/or structural integrity (Sharma *et al*., 2018; Vásquez-Limeta *et al*., 2022; Tang *et al*., 2009; Schmidt *et al*., 2009; Chang *et al*., 2016; Sharma *et al*., 2016, 2021). Although HYLS1 loss of mutation did not affect the recruitment of these proteins, it drove a strong decrease in the recruitment of the downstream CEP295 and CEP120 interacting proteins: POC5, C2CD3, and Talpid3. In particular, C2CD3 and Talpid3 are recruited by CEP120 to the procentrioles during centriole duplication and stabilize the centriole distal end (Wang *et al*., 2018). We conclude that HYSL1 governs early steps of procentriole assembly by facilitating the recruitment of proteins required for centriole structural integrity and distal end assembly.

In contrast to what has been shown in *C. elegans* (Dammermann *et al*., 2009), we did not observe an interaction between HYLS1 and CPAP (**Figure S7F**). In addition, HYLS1 did not bind tubulin isoforms required for the assembly of the centriolar microtubules, such as the ε-tubulin (data not shown). Rather, we found that HYLS1 robustly interacted with CEP120, and the HYLS1 D211G mutation weakened this interaction. Since CEP120 is required for HYLS1 recruitment to the centriole, our data suggest that defective binding of HYLS1 D211G to CEP120 leads to reduced HYLS1 D211G recruitment to the centriole. This in turn impairs the recruitment of proteins required for the cohesion of centriolar microtubules (POC5 and Centrin2) and centriole distal end assembly (Talpid3 and C2CD3). The disruption of these centriole elongation and centriole stability factors creates a fragile microtubule wall that breaks to produce short centrioles that lack distal appendages and are unable to ciliate (**Figure 5I**). We conclude that tissue-specific defects in centriole structural integrity are a key driver of HLS syndrome.

## MATERIAL AND METHODS

### Animals

Mice were housed and cared for in an AAALAC-accredited facility. All animal experiments were approved by the Johns Hopkins University Institute Animal Care and Use Committee (MO21M300).

### Mouse model

*Hyls1^+/DG^* mice were generated using CRISPR/Cas9 technology as previously described (Phan *et al*., 2022; Sladky *et al*., 2022). Briefly, one sgRNA (5’-CTACTCGGTCTATCTTGCCC-3’) was co-injected with a ssDNA repair template (5’-CAGTCTCTTTTGTACTCAAAATATCTGGCTACTCGACCCATTTTTCCCCGGTTTCGGCTTAACTGATCCAGTCTGGGGAG-3’) containing the GAC>GGT modification coding for the amino acid 211 as well as a restriction cut site for Taq1 at the site of the edit. Johns Hopkins University Transgenic Core performed pronuclear injection of one-cell B6SJL/F2 embryos (The Jackson Laboratory). Injected embryos were transferred into the oviducts of pseudopregnant females. Offspring resulting from embryo injections were genotyped and sequenced to check for the presence of the edit. The following primers were used for PCR amplification: forward (5′-GAACGAATGTTAGCTGCTGC-3′) and reverse (5′-GCGGGAAGGGAAGCTTCTTG-3′).

### Mouse tissue collection and sectioning

Kidneys of P0 mice were harvested and embedded fresh in O.C.T. compound (VWR Scientific). Brains of P0 mice were harvested, fixed in 4% paraformaldehyde (PFA) for 2 hr, and embedded in 30% sucrose for 3 days at 4C. This was followed by mounting the brains in O.C.T. compound. Tissues in OCT were frozen for cryosectioning (Leica CM1950 cryostat), and 20µm sections were collected on Superfrost Plus microscope slides (Thermo Fisher Scientific). To obtain the epidermal sheet, the skin from P0 mice (dermal side up) was placed on a precooled glass plate placed on crushed ice. The dermis was scraped off until the epidermis was reached. The epidermal sheet was fixed with 4% PFA in PBS for 10 min at room temperature (RT).

### MCCs

Mouse ependymal cell cultures were prepared as previously described (Delgehyr *et al*., 2015; LoMastro *et al*., 2022). Briefly, brains were dissected from P0 mice in Hank’s solution (10% HBSS, 5% HEPES, 5% sodium bicarbonate, and 1% penicillin/streptomycin (P/S) in pure water) and the telencephalon was cut into pieces. After enzymatic digestion (DMEM GlutaMAX, 1% P/S, 3% papain [Worthington 3126], 1.5% 10 mg/ml DNAse, and 2.4% 12 mg/ml cysteine) for 45 min at 37°C, the digestion was stopped by a solution of trypsin inhibitors (Leibovitz’s L15 medium, 10% 1 mg/ml trypsin inhibitor [Worthington], and 2% 10 mg/ml DNAse) and cells were washed with L15 medium. Cells were resuspended in DMEM GlutaMAX supplemented with 10% fetal bovine serum (FBS) and 1% P/S and grown for 4-5 days on poly-L-lysine-coated flasks. Once cells were confluent, they were incubated overnight with vigorous shaking (250 rpm), and then re-plated at a density of 2×10^5^ cells/cm^2^ on poly-L-lysine-coated coverslips in DMEM GlutaMAX, 10% FBS and 1% P/S. The following day, the medium was replaced with serum-free DMEM GlutaMAX and 1% P/S to trigger ependymal differentiation *in vitro* (differentiation day 0). For immunostaining, cells were fixed on differentiation day 7 with 4% PFA for 20 min at RT or with 100% ice cold methanol for 10 min at -20°C.

### Limb staining

Limbs were stained as previously described (Rigueur & Lyons, 2014) with the modification that only bone was stained. Briefly, limbs were fixed in 95% ethanol. The samples were transferred to acetone overnight at RT. Samples were then placed in 80% ethanol and 20% (glacial) acetic acid overnight. After washing two times with 70% ethanol, limbs were incubated in 95 % ethanol overnight. To pre-clear the tissue, the 95% ethanol was removed, and 1% potassium hydroxide (KOH) solution was added for 1 hr at RT. The KOH solution was replaced by Alizarin red solution (0.005 % (w/v) in 1% (w/v) KOH) for 3-4 hr at RT and then replaced with a 50/50 glycerol/KOH (1%) solution for 30 min. The glycerol/KOH solution was changed, and samples were incubated at RT for 3-4 days. If excess dye persisted the glycerol/KOH was changed again and maintained until imaging. After imaging, samples were transferred to 100% glycerol for long-term storage.

### Cell culture

MEFs were cultured in Dulbecco’s modified Eagle’s medium (DMEM-1X, Corning) containing 4.5g/L glucose & L-glutamine without sodium pyruvate, supplemented with 10% FBS, 1% P/S, and 0.1mM β-mercaptoethanol. MEFs were maintained at 37°C in 5% CO_2_ and 3% O_2_ incubator. MEFs were frozen down in 60% complete DMEM, 30% FBS and 10% DMSO. RPE1 cells were cultured in Dulbecco’s modified Eagle’s medium/Hams F-12 50/50 Mix (DMEM/F-12 50/50 1X, Corning) without L-glutamine, supplemented with 10% fetal bovine essence (FBE), 1% P/S, 1% L-glutamine, and 0.5% sodium bicarbonate. RPE1 cells were frozen down in 60% complete DMEM/F-12 media, 30% FBE and 10% DMSO. DLD1 cells were cultured in Dulbecco’s modified Eagle’s medium (DMEM-1X, Corning) containing 4.5g/L glucose & L-glutamine without sodium pyruvate, supplemented with 10% FBE, 1% P/S, and 1% L-glutamine. DLD1 cells were frozen down in complete DMEM (95%) and DMSO (5%). Both RPE1 and DLD1 cells were maintained at 37°C in 5% CO_2_ incubators.

### Cell line generation

Wildtype (*Hyls1^+/+^*), heterozygous (*Hyls1^+/DG^*), and homozygous (*Hyls1^DG/DG^*) embryos from a CRISPR-Cas9 mediated HYLS1 D226G knock-in mouse model were harvested at E14.5 and transferred to a dish containing sterile 1X PBS. After the heart, liver and brain were removed, each embryo was transferred to a new dish containing 0.25% trypsin-EDTA (Thermo Scientific) and cut into fine pieces. The processed embryo was transferred to a 15mL tube, the volume was filled to 3mL with 0.25% trypsin-EDTA and incubated overnight at 4°C. The tubes containing embryos were then incubated for 30 min at 37°C. MEF culture media was added to a final volume of 8 mL and the solution pipetted vigorously to break down the digested tissue into a cell suspension. The supernatant was transferred to a new tube and the previous step was repeated to collect more supernatant. The mixed solution was then cultured into a 10 cm tissue culture dish. CRISPR-Cas9-mediated technology was used to generate *HYLS1^-/-^* DLD1 and RPE1 cells as previously described (Gliech *et al*., 2024). Briefly, HYLS1 crRNA was cloned into the Lenti-CRISPR-V2 backbone (a gift from Feng Zhang; Addgene plasmid #52961) and transfected with the pCMV-VSV-G (a gift from Bob Weinberg, Addgene plasmid #8454) and the psPAX2 (a gift from Didier Trono, Addgene plasmid #12260), in HEK 293T cells using calcium phosphate. 48h later, viral supernatant was harvested, filtered, mixed with polybrene (10µg/mL – Sigma-Aldrich) and administered to cells for 24 hr. Transduced cells were then selected with puromycin (2.5µg/mL) for 72 hr. We made use of the Flp-In System to express *HYLS1^WT^* or *HYLS1^DG^* tagged with either 1x-mNeonGreen, 1x-mCherry, or 1x-HA (cloned into the pcDNA5 FRT/TO plasmid) in DLD1 *HYLS1^+/+^* or *HYLS1^-/-^* cells. Cell lines were generated by transfecting the pcDNA5 FRT/TO plasmids with the POG44 Flp-recombinase plasmid into DLD1 Flp-In T-Rex cells. Cells were then selected with 400 µg/mL hygromycin B for at least 2 weeks. Expression of reporters was induced with 1 µg/mL of doxycycline. Knock-in of an endogenous 1xHA tag on the C-terminus of the *HYLS1* gene (*HYLS1^WT^*^-^HA) or an endogenous knock-in of the HYLS1 D211G mutation in *HYLS1^WT^*^-^HA RPE1 cells was generated as previously described (Ghetti *et al*., 2021).

### Treatments in cultured cells

To rescue centriole integrity with Taxol treatment, RPE1 cells were arrested in late G1 with Palbociclib (PB – 200nM), a CDK4/6 inhibitor for 24 hr. Cells were then released into fresh media containing Dimethylenastron (DMN - 2µM) to arrest the cells in mitosis. 12 hr after the initial release from PB (cells in G2-phase) DMSO or Taxol (10µM) were added, and the cells were kept overnight in mitosis. The following morning, RO3306 (10µM), a Cdk1 inhibitor, was added to the cells to force them to exit mitosis. Cells (in G1-phase) were fixed 5 hr later followed by immunofluorescence or U-ExM.

To induce ciliogenesis in MEFs and RPE1 cells, cells were seeded on coverslips and serum-starved by adding the corresponding culture media without FBS/FBE for 48hr. Cells were fixed in 4% PFA and treated accordingly for immunofluorescence or U-ExM. Hedgehog signaling response was analyzed by treating serum starved MEFs with 500nM of SAG (Sigma Aldrich #566660), 24 hr before the cells were fixed.

### CEP120 siRNA-mediated gene depletion

DLD1 cells were transfected with 100 nM CEP120 siRNA (ThermoFisher #s45768) using Lipofectamine RNAiMAX Transfection Reagent (Life Technologies), according to the manufacturer’s guidelines. Cells were fixed for immunofluorescence analysis 48 hr after transfection.

### Immunofluorescence staining in cultured cells

Cells were grown on glass coverslips and fixed for 10 min in ice-cold MeOH at −20°C or 20 min in 4% PFA at RT. MEFs, RPE1, and DLD1 cells were blocked in blocking solution containing 2.5% FBS, 200 mM glycine, 0.1% Triton X-100 in 1X PBS for 1 hr, and MCCs blocked for 1-2 hr in blocking buffer containing 1X PBS, 0.1% Triton X-100, 10% normal donkey serum at RT. Cells were then incubated in primary antibody diluted in the blocking solution for 1 hr (MEFs, RPE1, and DLD1 cells) or overnight (MCCs) at RT and subsequently washed with PBST (0.1% Triton X-100 in 1X PBS) three times for 10 min each. Secondary antibodies were added for 1-2 hr at RT. DNA was stained with DAPI for 5 min and cells washed with PBST three times for 10 min each. Coverslips were then mounted on microscope slides using ProLong Gold antifade mountant (Invitrogen).

### Immunofluorescence staining in tissue sections

Slides containing tissue sections were washed in 1X PBS before antigen retrieval was performed using L.A.B. solution (Polysciences #24310-500) for 10 min at RT. After washing with 1X PBS the sections were blocked for 1 hr at RT in blocking solution (10% goat serum, 0.2% Triton-X, 1X PBS). Primary antibodies were incubated for 3 hr at RT or overnight at 4°C in blocking buffer. After washing the sections three times with 1X PBS, secondary antibodies were incubated for 3 hr at RT. After washing the slides three times with 1X PBS, the tissue sections were mounted using ProLong Gold Antifade Mountant (Invitrogen).

### Ultrastructure expansion microscopy (U-ExM)

U-ExM was performed based on a previously published protocol (Gambarotto *et al*., 2021b, 2021a). Briefly, cells were seeded on a 25 mm (MEFs, RPE1 and DLD1) or a 12 mm (MCCs at differentiation day 6) glass coverslip in a 6-well plate. Cells were rinsed with 1X PBS, and 2 mL of the anchor solution (1X PBS, 1.4% formaldehyde (FAA), 2% acrylamide mixture prepared fresh) was added. The plate was incubated at 37°C for 5 hr (MEFs, RPE1 and DLD1) or overnight (mouse tissues and MCCs). A coverslip was then placed on top of an 850 µL (MEFS, RPE1 and DLD1) or 50 µL (MCCs) drop of precooled monomer solution (1X PBS, 23% sodium acrylate (w/v), 10% acrylamide, 0.1% N,N′-methylenbisacrylamide, 0.5% TEMED and 0.5% ammonium persulfate) sitting on top of parafilm in a humid chamber. For tissue sections, the monomer solution was added on top of the slide containing the tissue section and a coverslip placed on top. The humidified chamber was incubated on ice for 30 min (MEFs, RPE1, DLD1, and MCCs) or for 60 min (mouse tissues) and then moved to 37°C for 1 hr (MEFs, RPE1, DLD1, and MCCs) or for 2 hr (mouse tissues). Following gel polymerization, a 4mm biopsy punch was used to create several punches from each coverslip. Punches were transferred to a 50 mL Falcon tube (MEFs, RPE1, DLD1, and tissue sections) or into a 1.5 ml Eppendorf tube containing denaturation buffer (200 mM SDS, 200 mM NaCl, 50 mM Tris) and incubated for 15 min at RT with gentle agitation. The 50 mL Falcon tubes were transferred to a water bath at 95°C for 1.5 hr with gentle agitation every 20 min. The Eppendorf tubes containing the MCC punches were incubated at 95°C for 1 hr. Punches from mouse tissues were kept in denaturation buffer with gentle agitation for an additional 1 hr at RT. Punches were washed in water three times for 20 min each and kept overnight at RT with gentle agitation for the first round of gel expansion. Expanded punches were transferred to 1X PBS for 1 hr at RT and blocked with 2% BSA in PBS at 37°C for 30 min - 1 hr. This step reduced the gel size to ∼2x the original 4 mm width. Primary antibodies were diluted in 2% BSA in PBS and stained for 3 hr (MEFs, RPE1, DLD1, and tissue sections) or overnight (MCCs) at 37°C with gentle agitation. After washing the punches three times for 10 min at RT in PBST (1X PBS, 0.1% Tween-20), secondary antibodies and DAPI were added for 3 hr at 37°C. This was followed by another round of washes with PBST (three times for 10 min at RT) and 3 washes with water for 20 min each. Gels were incubated in water overnight at room temperature with gentle agitation for the second round of gel expansion. Expansion factors were calculated by measuring the expanded punches with calipers and dividing the value measured by 4 (initial size of the gel after using the 4 mm-biopsy punch). The gel expansion was consistently 4.0-4.1x.

### Cell cycle analysis

Cell cycle profile analysis was performed by staining fixed cells (70% ethanol at -20°C for at least 30 min) with a propidium iodide solution. After fixation, the cells were washed three times with 1X PBS and stained with propidium iodide (50µg/mL) containing RNaseA (100µg/mL) in the dark at 37°C for 30 min. Subsequently, the cells were analyzed by flow cytometry using the BD FACSCalibur Flow Cytometer (BD Biosciences). Quantification analysis of cell cycle profiles was generated using the FlowJo software package (FlowJo, LLC; USA).

### Antibodies used for image analysis

The following primary antibodies were used for immunofluorescence or U-ExM in MEFs, RPE1, or DLD1 cells: anti-HA (Rat, Roche #ROAHAHA, 1:500), anti-SAS-6 (Mouse, Santa Cruz Biotechnology #sc-81431, 1:1000), anti-CEP135 (Rabbit, a gift from A. Hyman, Max Planck Institute for Molecular Cell Biology and Genetics, Dresden, Germany, 1:1000), anti-CPAP (Rabbit, Proteintech #11517-1-AP, 1:1000), anti-CEP120 (Rabbit, a gift from M. Mahjoub, Washington University in St Louis, USA, 1:1000), anti-Centrin2 (Rabbit, Homemade (Moyer & Holland, 2019), 1:1000), anti-POC5 (Rabbit, Bethyl Laboratories #a303-341, 1:1000), anti-CP110 (Rabbit, Proteintech #12780, 1:1000), anti-Talpid3 (Rabbit, Proteintech #24432-1-AP, 1:1000), anti-C2CD3 (Rabbit, Sigma-Aldrich #HPA038552, 1:1000), anti-CEP164 (Rabbit, EMD Millipore #ABE2621, 1:1000), anti-ANKRD26 (Rabbit, GeneTex #GTX128255, 1:1000), anti-Acetylated α-tubulin (Mouse, Santa Cruz Biotechnology #sc-23950, 1:1000), anti-Smoothened (Mouse, Santa Cruz Biotechnology #sc-166685, 1:500), anti-α-Tubulin (Guinea Pig, Geneva Antibody Facility #AA344, 1:1000), anti-β-Tubulin (Guinea Pig, Geneva Antibody Facility #AA345, 1:1000), anti-Arl13b (Mouse, NeuroMab/Antibodies Incorporated #75-287, 1:1000), γ-Tubulin (Goat, Homemade, 1:1000). The following primary antibodies were used for MCC staining: anti-Acetylated-α-Tubulin (Lys40) (Mouse, Cell Signaling #12152, 1:1000), anti-SAS-6 (Mouse, Santa Cruz Biotechnology # sc-81431, 1:250), anti-Deup1 (Rabbit, homemade, 1:1000), anti-β-Tubulin (Guinea Pig, Geneva Antibody Facility #AA345, 1:1000), anti-Centrin2 (Rabbit, homemade, 1:1000). The following secondary antibodies were used for immunofluorescence or U-ExM: Alexa Fluor conjugated donkey anti-mouse A488 (#A21202; Thermo Fisher Scientific), donkey anti-rat A488 (#A21208; Thermo Fisher Scientific), goat anti-guinea pig A488 (#A11073; Thermo Fisher Scientific) donkey anti-mouse A555 (#A31570; Thermo Fisher Scientific), donkey anti-rabbit A555 (#A31572; Thermo Fisher Scientific), goat anti-guinea pig A555 (#A21435; Thermo Fisher Scientific), donkey anti-rabbit A647 (#A31573; Thermo Fisher Scientific), donkey anti-rat A647 (#A48272; Thermo Fisher Scientific).

### Protein co-immunoprecipitation

*HYLS1^-/-^*, and *HYLS1^-/-^* with HYLS1-mCherry (WT or DG) add-back DLD1 cells were induced with doxycycline (1µg/mL) and 48 hr later transiently transfected with the construct of interest cloned into the pcDNA5 FRT/TO plasmid (Life Technologies). 48 hr after the transient transfection, the cells were washed with 1X PBS and then 2mL of PBS was added to the dish. Using a cell scraper, cells were scraped from the plate into the small amount of remaining PBS and transferred to a 15 mL conical tube on ice. After centrifugation for 5 min at 1500 rpm at 4°C, supernatant was removed, and cells were snap frozen. The pellet was resuspended in supplemented lysis buffer (10mM Tris pH 7.5, 250mM NaCl, 1mM EDTA, 50mM NaF, 50mM β-glycerophosphate, 0.1%Triton X-100, supplemented with fresh 1% PMSF, 1mM DTT, and cOmplete mini EDTA-free protease inhibitors (Sigma-Aldrich)) and transferred to a 2 mL Eppendorf tube. After vortexing briefly, cells were kept on ice for 30 min. Each tube was sonicated three times for 6 s at 30% duty cycle and put on ice between the sonication cycles. Cells were centrifuged for 15 min at 15000 rpm at 4°C. For each sample, 5% of the lysate was saved as ‘input’, and the remaining clarified lysate was added to the pre-washed beads coupled to mCherry-binding protein (Rothbauer *et al*., 2008) and gently tumbled at 4°C for 4 hr. Beads were then spun down at 4°C at 1000 rpm for 1 min and the supernatant was discarded. Beads were washed three times with 1 mL of lysis buffer and, immunopurified protein was analyzed by immunoblot. Analysis of HYLS1 binding to CEP120 was assessed by normalizing the intensity of the elution fraction of the target protein to the intensity of the elution fraction of the bait protein using the following equation: Relative Elution Intensity = (Intensity_CEP120_/Intensity_HYLS1_)_DG-Elution_/(Intensity_CEP120_/Intensity_HYLS1_)_WT-Elution_.

### Immunoblotting analysis

Cell lysates were loaded on denaturing SDS-polyacrylamide gels and separated by gel electrophoresis under constant volt (100V) on a Mini Gel Tank (Thermo Fisher). The separated proteins were transferred to a nitrocellulose membrane using a Mini Trans-blot Module (Bio-Rad) at constant 100V for 60-90 min, depending on the size of the protein of interest. Membranes were then incubated in blocking buffer (5% dried milk powder with 0.1% Tween-20 in 1X PBS) for 1 hr at RT, and primary antibodies (diluted in 3% BSA in 1X PBS) were subsequently incubated overnight at 4°C. Membranes were then washed three times (10 min each) with washing buffer (0.1% Tween-20 in 1X PBS) and incubated with the secondary antibodies (diluted in 3% BSA in 1X PBS) for 1 hr at RT. This step was followed by three, 10 min washing steps with washing buffer, before the membrane was incubated with WesternBright ECL (Advansta) for 3 min. Chemoluminescent signal was visualized using a Genesys G:Box Chemi-XX6 system (Syngene). The following primary antibodies were used: anti-HA (Rat, Roche #ROAHAHA, 1:500), anti-RFP (Rabbit, Abcam #ab167453, 1:500). The following secondary antibodies were used: HRP-conjugated anti-rat (Goat, Cell Signaling #7077, 1:5000), or HRP-conjugated anti-rabbit (Goat, Cell Signaling #7074, 1:5000).

### Image and data analysis

Immunofluorescence images were acquired using a DeltaVision Elite system (GE Healthcare) controlling a scientific CMOS camera (pco.edge 5.5). Images were acquired with Applied Precision immersion oil (n = 1.516) at 100X or 60X with 0.2 μm z-sections. Acquisition parameters were controlled by SoftWoRx suite (GE Healthcare). U-ExM images of MEFs, RPE1, and DLD1 cells were acquired using a Leica SP8 confocal microscope with a 63X/1.4 NA oil immersion objective with 0.2 µm z-step size and a pixel size of 22.55nm. Immunofluorescence and U-ExM MCC images were acquired using a Leica SP8 confocal microscope with a 40X plan-apochromat oil immersion objective with 1.30 NA, or a Zeiss Axio Observer 7 inverted microscope with Slidebook 2023 software (3i—Intelligent, Imaging Innovations, Inc.), CSU-W1 (Yokogawa) T1 Super-Resolution SoRa Spinning Disk, and Prime 95B CMOS camera (Teledyne Photometrics) with a 40X plan-apochromat oil immersion objective with 1.30 NA. Images were deconvolved with Leica’s lightning process software or Microvolution software built into Slidebook. Images were processed in FIJI (Schindelin *et al*., 2012). All images presented in figures are Z-stack maximum intensity projections. All imaging analysis was performed blinded using ImageJ (v2.1.0/1.53c, National Institutes of Health, http://imagej.net). Statistical analysis was performed using GraphPad Prism software.

## Supporting information

Extended Statistical Analysis

## ACKNOWLEDGMENTS

We thank the Holland Lab for fruitful discussions and helpful comments. AJH acknowledges funding by the National Institutes of Health (R01 GM133897, R01 GM114119, R01 CA266199).

## AUTHOR CONTRIBUTIONS

Author contributions: A.C. and A.J.H conceptualized the study. A.C. performed and analyzed most of the experiments and prepared the figures. Z.H. performed and analyzed most of the immunofluorescence analysis. M.A.S. assisted with dissection, and analysis of *Hyls1^DG/DG^* mice. C.R.G. assisted with cloning and analysis of experiments. C.E.J. performed MCCs experiments and analysis *in vitro*. T.A. and A.F. performed the initial experiments that identified the HYLS1 loss phenotype. A.C., A.J.H., C.R.G. wrote, reviewed, and edited the manuscript.

## COMPETING INTERESTS

The Authors declare that they have no competing interests.

---

**Figure S1:**
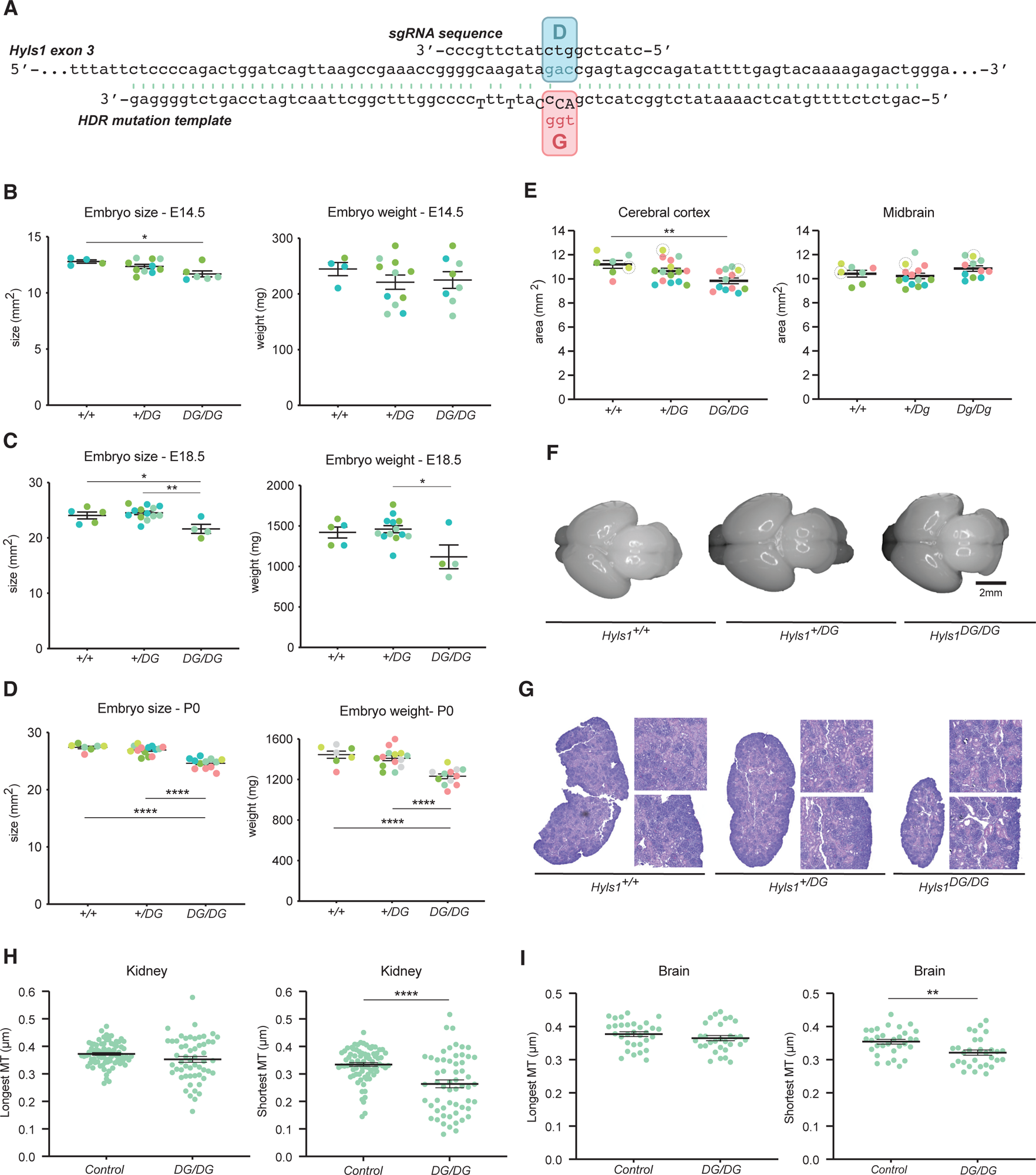
Hyls1 D226G leads to smaller animals with kidney developmental defects. (A) Schematic representation of the strategy used to generate the *Hyls1 D226G* mouse model. (B-D) Embryo size and weight at E14.5 (B), E18.5 (C), and P0 (D) developmental stages in *Hyls1^+/+^*, *Hyls1^+/DG^*, and *Hyls1^DG/DG^* animals. N≥4 mice per genotype. (E) Cerebral cortex (left) and Midbrain (right) area analysis in *Hyls1^+/+^*, *Hyls1^+/DG^*, and *Hyls1^DG/DG^* P0 animals. Dotted circles indicate the data point used for the representative images in (F). N≥7 mice per genotype. (F) Representative images from the brain size analysis in (E). (G) Histological analysis of kidneys in *Hyls1^+/+^*, *Hyls1^+/DG^*, and *Hyls1^DG/DG^* P0 animals. (H-I) Length analysis of the longer or shorter centriole microtubules by U-ExM in kidneys (H) or brain (I) of *Hyls1^+/+^*, *Hyls1^+/DG^*, and *Hyls1^DG/DG^* P0 animals. N≥2 mice per genotype. Data are represented as mean ± SEM. Statistical significance was assessed using one-way-ANOVA with Tukey’s multiple comparisons test (B-E) and an unpaired two-tailed Student’s t-test with Welch’s correction (H/I). (*) P < 0.05, (**) P < 0.01, (****) P < 0.0001. Only significant results are indicated.

**Figure S2:**
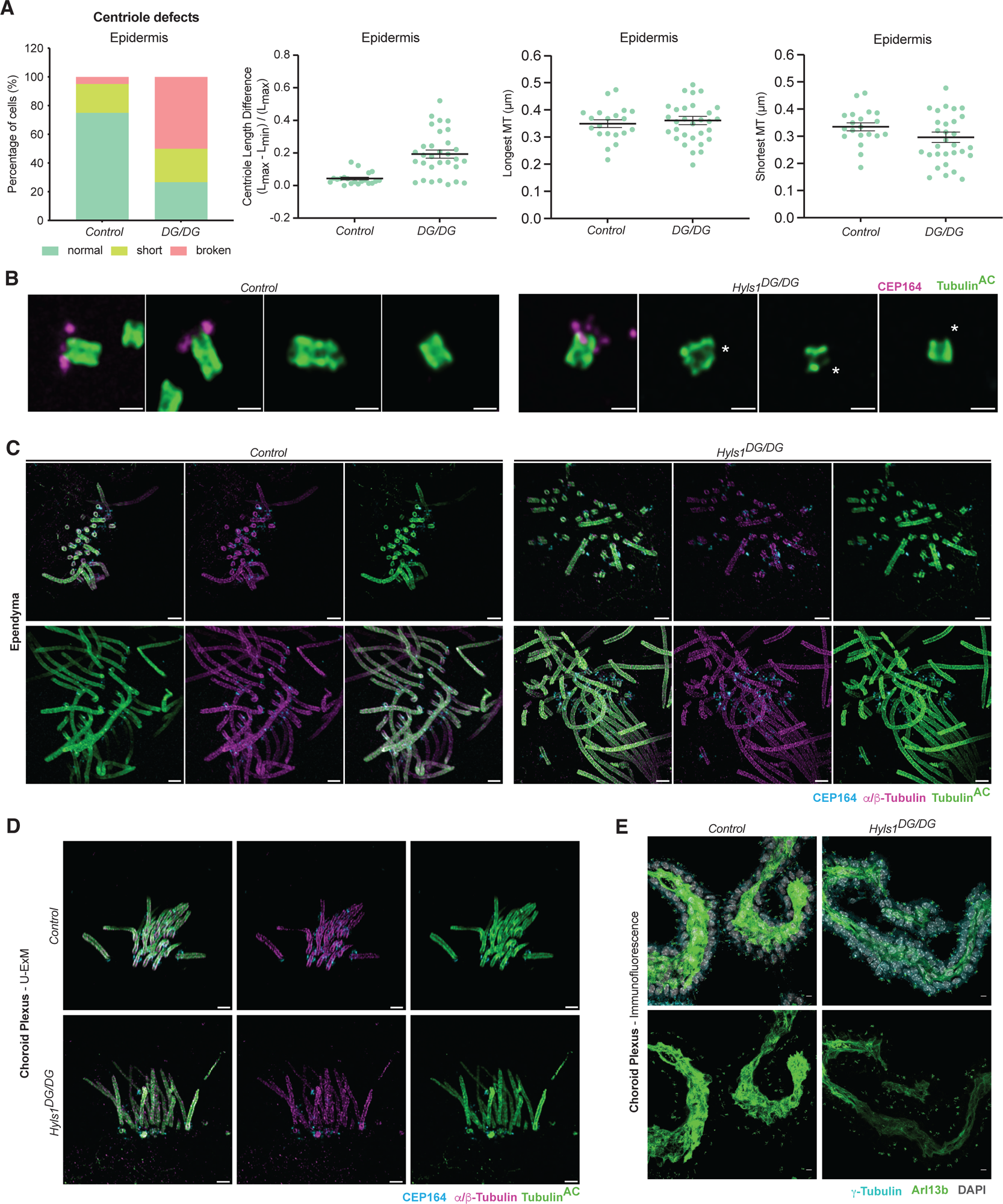
Hyls1 D226G does not impair MCC differentiation. (A) Quantification of centriole defects (left), the relative length difference between the longest and shortest microtubule in each centriole (middle), and maximum and minimum centriole length measurements (right) from U-ExM analysis of the epidermis of control *(Hyls1^+/+^* or *Hyls1^+/DG^)* and *Hyls1^DG/DG^* P0 animals. N=1 mouse per genotype. (B) Representative images of the analysis performed in (A). (C) U-ExM of MCCs in the ependyma of the lateral ventricles in the brain of control *(Hyls1^+/+^* or *Hyls1^+/DG^)* and *Hyls1^DG/DG^* P0 animals. (D) U-ExM analysis of MCCs in the choroid plexus in the brain of *controls (Hyls1^+/+^* or *Hyls1^+/DG^) and Hyls1^DG/DG^* P0 animals. (E) Immunofluorescence analysis of MCCs in the choroid plexus in the brain of *controls (Hyls1^+/+^* or *Hyls1^+/DG^)* and *Hyls1^DG/DG^*P0 animals. Scale bar is 250 nm in (B), 1µm in (C/D) and 10 µm in (E). Asterisk (*) indicates defective centrioles.

**Figure S3:**
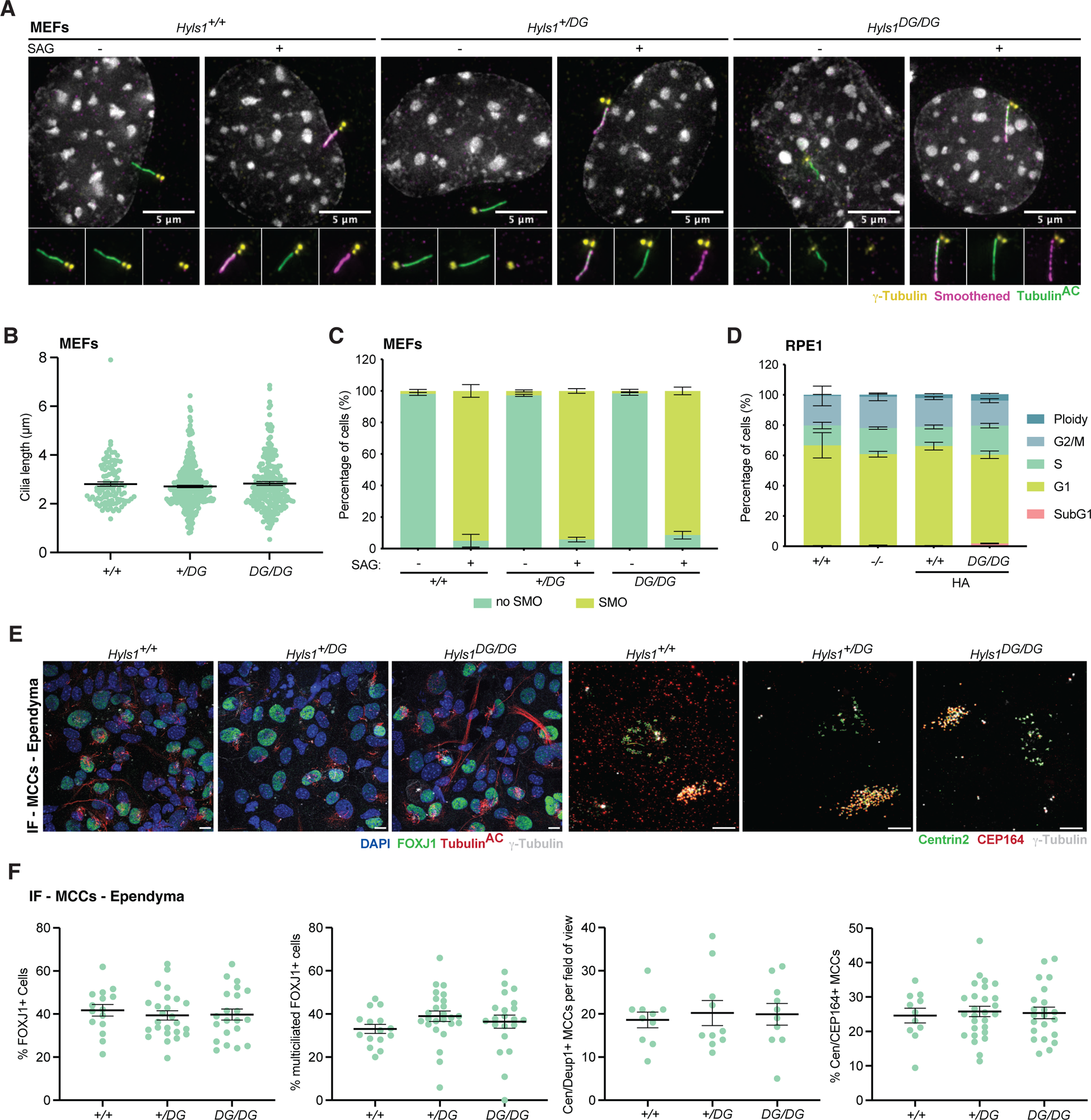
Cilia length and Hedgehog signaling are not impaired in the rare cilia formed in Hyls1 D226G MEFs. (A) Representative immunofluorescence images of the centrosome (γ-Tubulin), Smoothened, and cilia (Tubulin^AC^) in *Hyls1^+/+^*, *Hyls1^+/DG^*, and *Hyls1^DG/DG^* MEFs. (B) Cilia length analysis of *Hyls1^+/+^*, *Hyls1^+/DG^*, and *Hyls1^DG/DG^* MEFs. MEFs generated across N≥3 mice per genotype were analyzed. (C) Quantification of Smoothened recruitment to cilia after SAG treatment from immunofluorescence images of *Hyls1^+/+^*, *Hyls1^+/DG^*, and *Hyls1^DG/DG^* MEFs. MEFs generated across N≥3 mice per genotype were analyzed. (D) Cell cycle profile analysis of *HYLS1^+/+^*, *HYLS1^-/-^*, *HYLS1^+/+^*-HA and *HYLS1^DG/DG^*-HA RPE1 cells. Data from n=3 biological replicates were analyzed. (E) Representative immunofluorescence images of the centrosome (γ-Tubulin), FOXJ1, and cilia (Tubulin^AC^) (left), and of the centrosome (γ-Tubulin), centriole (Centrin2), and distal appendages (CEP164) (right) in *Hyls1^+/+^*, *Hyls1^+/DG^*, and *Hyls1^DG/DG^* in ependymal MCC. (F) Quantification of differentiated MCC abundance from immunofluorescence images of *Hyls1^+/+^*, *Hyls1^+/DG^*, and *Hyls1^DG/DG^*ependymal cells. Cells generated across N≥2 mice per genotype were analyzed. Data are represented as mean ± SEM. Statistical significance was assessed using one-way-ANOVA with Tukey’s multiple comparisons test (B/F). Only significant results are indicated. Smoothened (C) and cell cycle profile (D) analyses were assessed using two-way-ANOVA with post-hoc analysis, and results are summarized in supplementary material. Scale bar is 5 µm in (E).

**Figure S4:**
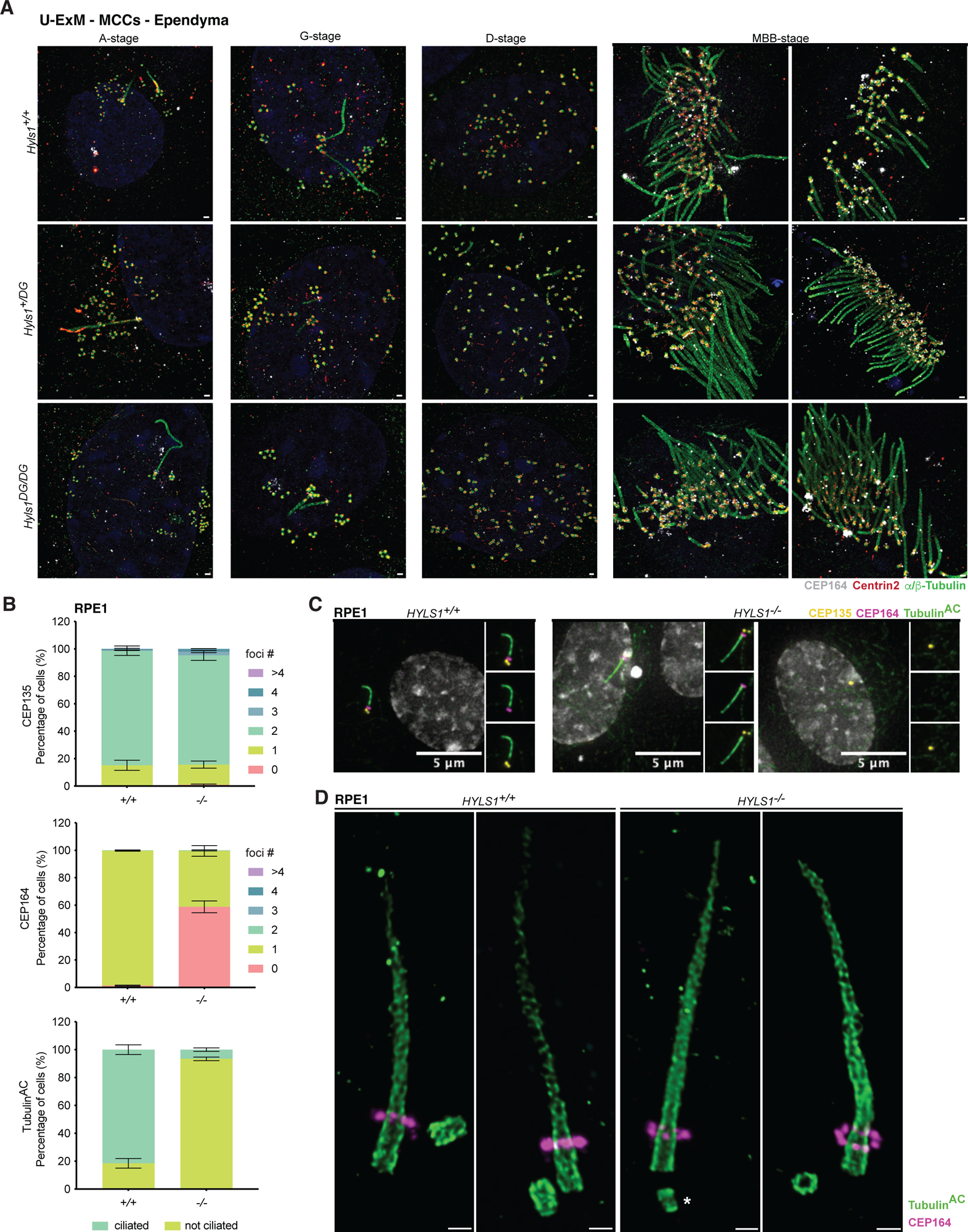
Hyls1 D226G does not affect MCC differentiation in vitro. (A) U-ExM analysis of different stages of MCC differentiation in ependymal cells *in vitro* stained for centrioles and cilia (α/μ-Tubulin), Centrin2 and distal appendages (CEP164) in *Hyls1^+/+^*, *Hyls1^+/DG^*, *Hyls1^DG/DG^* cells. (B) Quantification of centrosomes (CEP135), distal appendages (CEP164) and cilia (Tubulin^AC^) in *HYLS1^+/+^* and *HYLS1^-/-^* RPE1 cells. Data from n=3 biological replicates were analyzed. (C) Representative immunofluorescence images of centrosomes (CEP135), distal appendages (CEP164) and cilia (Tubulin^AC^) in *HYLS1^+/+^* and *HYLS1^-/-^* RPE1 cells. (D) U-ExM analysis of distal appendages (CEP164) and cilia (Tubulin^AC^) in *HYLS1^+/+^* and *HYLS1^-/-^* RPE1 cells. Data are represented as mean ± SEM. Statistical significance was determined using two-way-ANOVA with post-hoc analysis (B) and results are summarized in supplementary material. Scale bar is 500 nm in (A) and 250 nm in (D). Asterisk (*) indicates defective centrioles.

**Figure S5:**
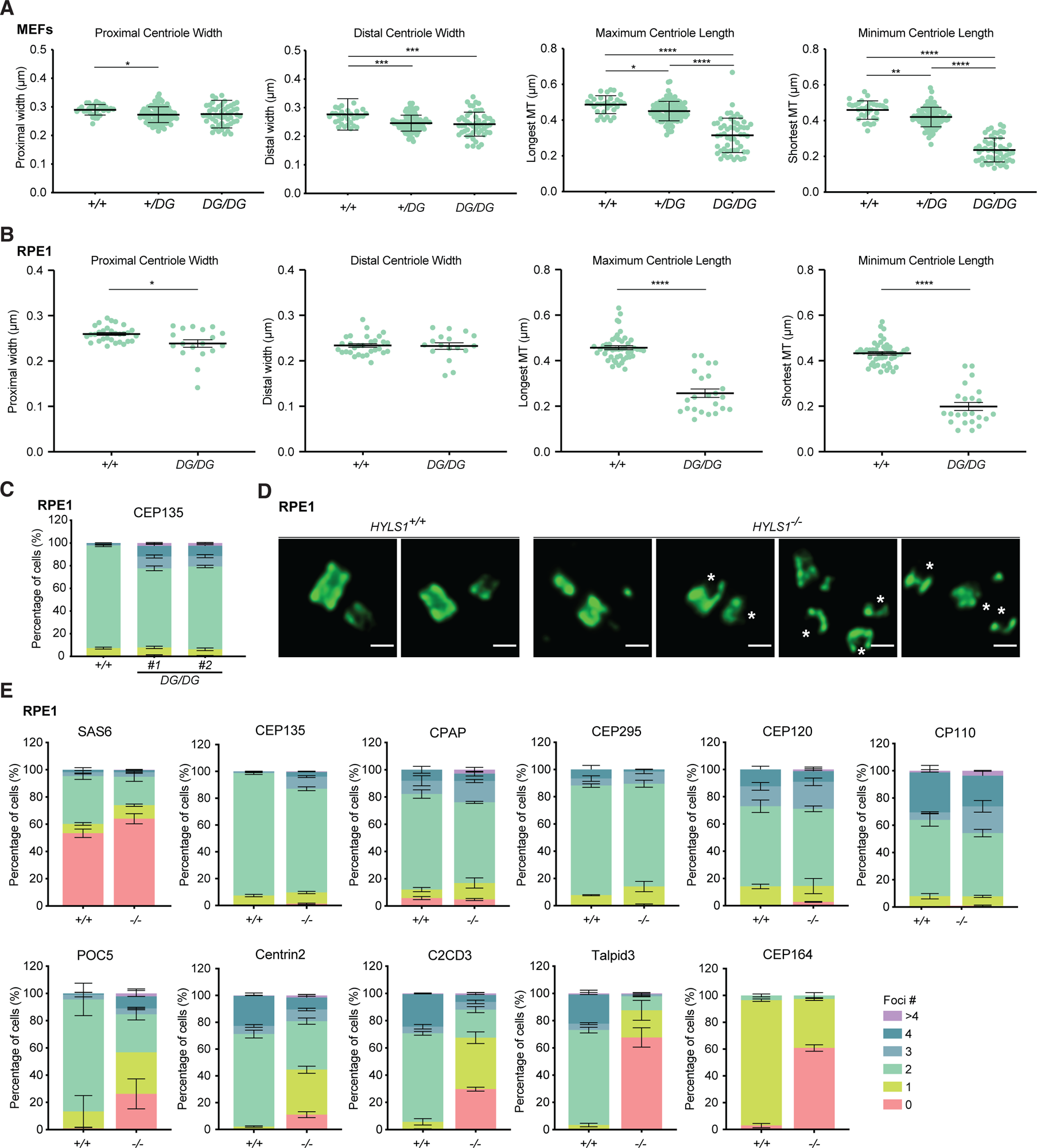
HYLS1^-/-^ leads to impaired recruitment of inner scaffold and centriole distal end proteins. (A) Quantification of centriole width and length from U-ExM images of *Hyls1^+/+^*, *Hyls1^+/DG^*, and *Hyls1^DG/DG^* MEFs. MEFs generated across N≥2 mice per genotype were analyzed. (B) Quantification of centriole width and length from U-ExM images of *HYLS1^+/+^* and *HYLS1^DG/DG^* RPE1 cells. Data from n=2 biological replicates were analyzed. (C) Quantification of centrioles (CEP135) from immunofluorescence images of *HYLS1^+/+^* and *HYLS1^DG/DG^* RPE1 cells. Data from n≥3 biological replicates were analyzed. (D) U-ExM analysis of centriole integrity in *HYLS1^+/+^* and *HYLS1^-/-^* RPE1 cells. (E) Quantification of proximal, inner scaffold, distal end, and distal appendage proteins from immunofluorescence images of *HYLS1^+/+^* and *HYLS1^-/-^* RPE1 cells. Data from n≥2 biological replicates were analyzed. Data are represented as mean ± SEM. Statistical significance was determined using one-way-ANOVA with Tukey’s multiple comparisons test (A), and an unpaired two-tailed Student’s t-test with Welch’s correction (B). (*) P < 0.05, (**) P < 0.01, (***) P<0.001, (****) P < 0.0001. Only significant results are indicated. Foci number analysis was assessed using two-way-ANOVA with post-hoc analysis, and results are summarized in supplementary material. Scale bar is 250 nm in (D). Asterisk (*) indicates defective centrioles.

**Figure S6:**
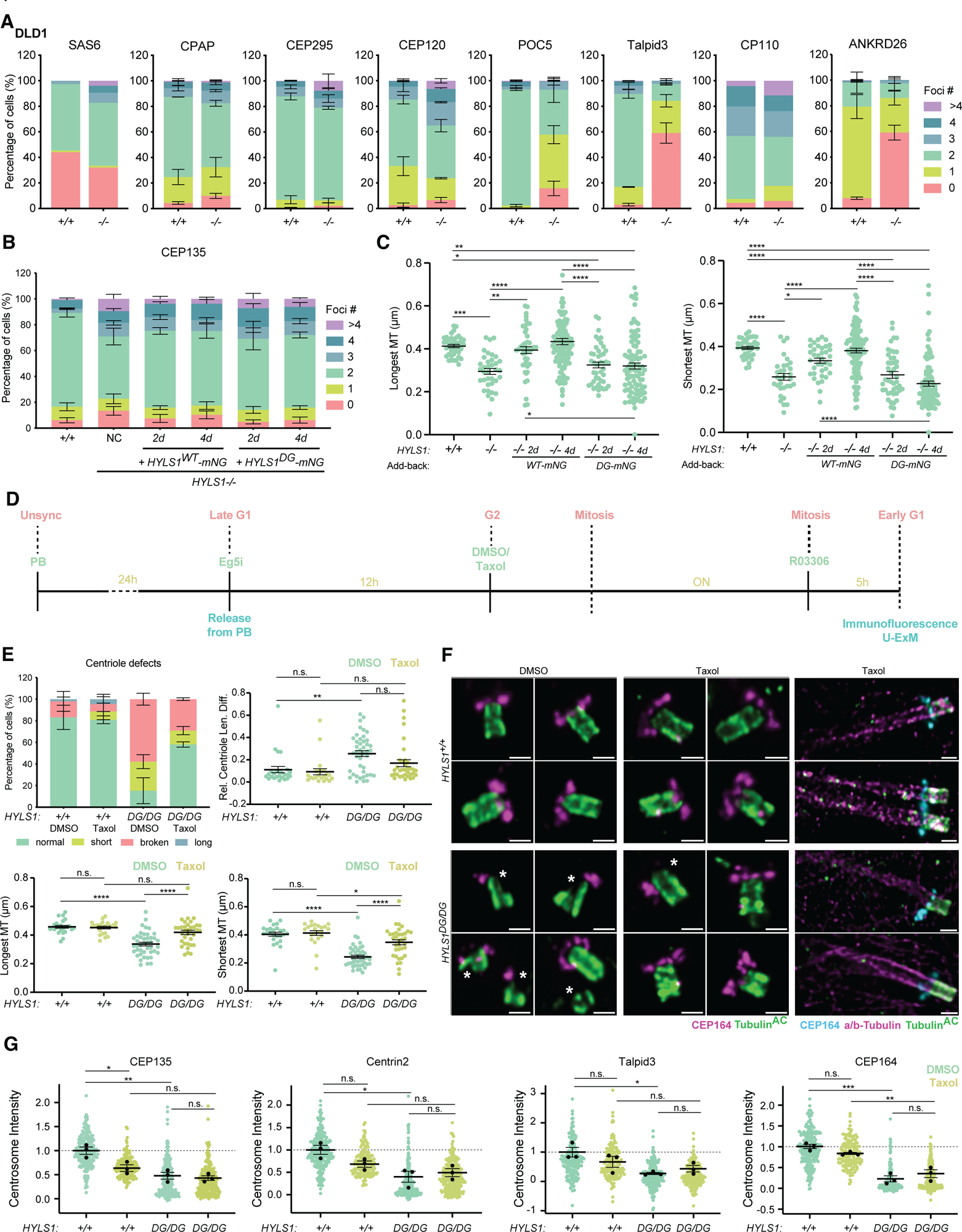
HYLS1 is required for centriole microtubule wall stability. (A) Quantification of proximal, inner scaffold, distal end, and distal appendage proteins from immunofluorescence images of *HYLS1^+/+^* and *HYLS1^-/-^* DLD1 cells. Data from n≥1 biological replicates were analyzed. (B) Quantification of centrosome number (CEP135) in *HYLS1^+/+^*, *HYLS1^-/-^*, and *HYLS1^-/-^*DLD1 cells with HYLS1-mNG (WT or D2111G) add-back. Data from n≥3 biological replicates were analyzed. (C) Quantification of maximum (left) and minimum (right) centriole length from *HYLS1^+/+^*, *HYLS1^-/-^*, and *HYLS1^-/-^* DLD1 cells with HYLS1-mNG (WT or D211G) add-back. Data from n=3 biological replicates were analyzed. (D) Schematic representation of the experimental strategy for microtubule stabilization by Taxol in mitosis in *HYLS1^+/+^* and *HYLS1^DG/DG^* RPE1 cells. (E) Quantification of centriole defects (top left), the relative length difference between the longest and shortest microtubule in each centriole (top right), maximum (bottom left) and minimum (bottom right) centriole length upon Taxol treatment from U-ExM images of *HYLS1^+/+^* and *HYLS1^DG/DG^* RPE1 cells. Data from n=2 biological replicates were analyzed. (F) Representative U-ExM images of centrioles (Tubulin^AC^ and α/μ-Tubulin) and distal appendages (CEP164) in *HYLS1^+/+^* and *HYLS1^DG/DG^* RPE1 cells treated with DMSO or Taxol. (G) Quantification of immunofluorescence signal intensity for centrioles (CEP135), Centrin2, Talpid3 and distal appendages (CEP164) in *HYLS1^+/+^* and *HYLS1^DG/DG^* RPE1 cells treated with DMSO or Taxol. Data from n=3 biological replicates were analyzed. Data are represented as mean ± SEM. Statistical significance was determined using one-way-ANOVA with Tukey’s multiple comparisons test from pooled data (C/E) or with mean values from each replicate (G). (*) P < 0.05, (**) P < 0.01, (***) P<0.001, (****) P < 0.0001. Only significant results are indicated (C). Foci number and centriole defects analysis were assessed using two-way-ANOVA with pot-hoc analysis and results are summarized in supplementary data. Scale bar is 250 nm in (F). Asterisk (*) indicates defective centrioles.

**Figure S7:**
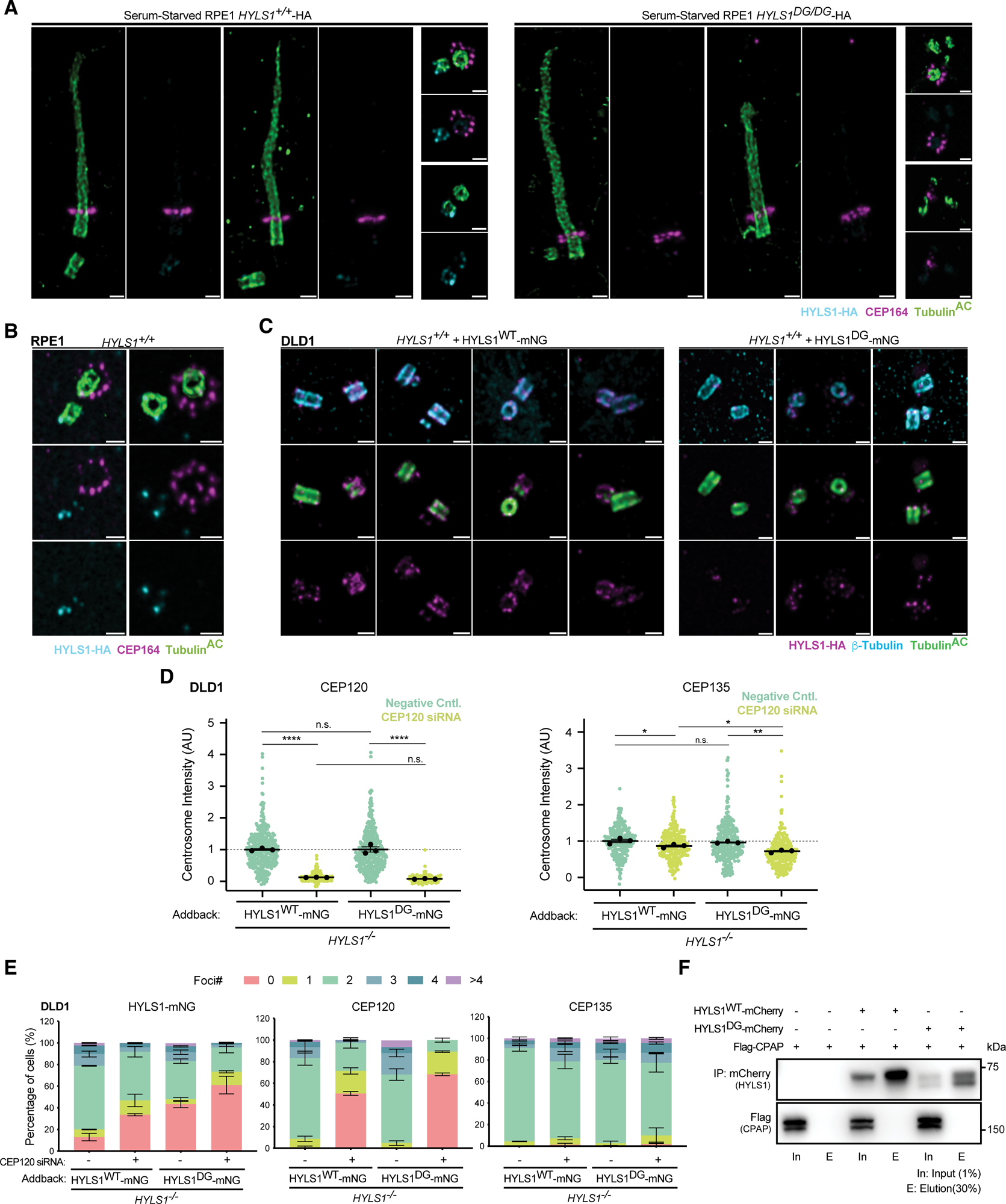
HYLS1 is asymmetrically localized to the younger parent centriole. (A) Representative U-ExM images of cilia (Tubulin^AC^), distal appendages (CEP164), and HYLS1-HA in *HYLS1^+/+^*-HA and *HYLS1^DG/DG^*-HA serum-starved RPE1 cells. (B) Representative U-ExM images of cilia (Tubulin^AC^), distal appendages (CEP164), and HYLS1-HA in *HYLS1^+/+^*-HA unsynchronized RPE1 cells. (C) Representative U-ExM images of centrioles (Tubulin^AC^ and β-Tubulin) and HYLS1-HA in *HYLS1^+/+^* DLD1 cells with HYLS1-mNG (WT or D211G) add-back. (D) Quantification of immunofluorescence signal intensity for CEP120 and centrioles (CEP135) in *HYLS1^-/-^* DLD1 cells with HYLS1-mNG (WT or D211G) add-back. Data from n=3 biological replicates were analyzed. (E) Quantification of centriole foci number for HYLS1-mNG, CEP120 and centrioles (CEP135) in *HYLS1^-/-^* DLD1 cells with HYLS1-mNG (WT or D211G) add-back. Data from n=3 biological replicates were analyzed. (F) Representative co-immunoprecipitation of Flag-CPAP with HYLS1^WT^-mCherry or HYLS1^DG^-mCherry in *HYLS1^-/-^* DLD1 cells with HYLS1-mCherry (WT or D211G) add-back. Data are represented as mean ± SEM. Statistical significance was determined using one-way-ANOVA with Tukey’s multiple comparisons test from mean values from each replicate (D). (*) P < 0.05, (**) P < 0.01, (****) P < 0.0001. Foci number analysis was assessed using two-way-ANOVA with post-hoc analysis and results are summarized in supplementary material. Scale bar is 250 nm in (A/B/C).

## Notes

### Competing Interest Statement

The authors have declared no competing interest.

