## Extended Statistical Analysis for "Centriole structural integrity defects are a crucial feature of Hydrolethalus Syndrome"

### Supplementary Material

#### Extended statistical analysis

#### FIGURE 1B

Statistics calculated for polydactyly analysis
Two-way anova with Tukey's multiple comparisons test

| **5 digits** | Summary | Adjusted P Value |
|---|---|---|
| +/+ vs. +/DG | ns | >0.9999 |
| +/+ vs. DG/DG | **** | <0.0001 |
| +/DG vs. DG/DG | **** | <0.0001 |

| **6 digits** | | |
|---|---|---|
| +/+ vs. +/DG | ns | >0.9999 |
| +/+ vs. DG/DG | **** | <0.0001 |
| +/DG vs. DG/DG | **** | <0.0001 |

| **7 digits** | | |
|---|---|---|
| +/+ vs. +/DG | ns | >0.9999 |
| +/+ vs. DG/DG | **** | <0.0001 |
| +/DG vs. DG/DG | **** | <0.0001 |

| **8 digits** | | |
|---|---|---|
| +/+ vs. +/DG | ns | >0.9999 |
| +/+ vs. DG/DG | ns | 0.7302 |
| +/DG vs. DG/DG | ns | 0.6651 |

#### FIGURE 1F

Statistics calculated for Cilia analysis
Unpaired t test with Welch's correction

| **Kidney** | Summary | Adjusted P Value |
|---|---|---|
| Control vs. DG/D | **** | <0.0001 |

| **Brain** | Summary | Adjusted P Value |
|---|---|---|
| Control vs. DG/D | ns | 0.2305 |

#### FIGURE 1D

Statistics calculated for Kidney size analysis
One-way anova with Tukey's multiple comparisons test

| **E18.5** | Summary | Adjusted P Value |
|---|---|---|
| +/+ vs. +/DG | ns | 0.865 |
| +/+ vs. DG/DG | **** | <0.0001 |
| +/DG vs. DG/DG | **** | <0.0001 |

| **P0** | Summary | Adjusted P Value |
|---|---|---|
| +/+ vs. +/DG | ns | 0.8343 |
| +/+ vs. DG/DG | **** | <0.0001 |
| +/DG vs. DG/DG | **** | <0.0001 |

#### FIGURE 1H

Statistics calculated for Centriole defects analysis
Two-way anova with Sidak's multiple comparisons test

**Kidney**

| Control vs. DG/DG | Summary | Adjusted P Value |
|---|---|---|
| normal | **** | <0.0001 |
| short | ns | 0.1061 |
| broken | **** | <0.0001 |
| long | ns | >0.9999 |

**Brain**

| Control vs. DG/DG | Summary | Adjusted P Value |
|---|---|---|
| normal | ns | 0.2259 |
| short | ns | 0.9984 |
| broken | ns | 0.324 |
| long | ns | >0.9999 |

Statistics calculated for longest MT/ Shortest MT ratio analysis
Unpaired t test with Welch's correction

| **Kidney** | Summary | Adjusted P Value |
|---|---|---|
| Control vs. DG/DG | **** | <0.0001 |

| **Brain** | Summary | Adjusted P Value |
|---|---|---|
| Control vs. DG/DG | ** | 0.0011 |

### FIGURE 2B

Statistics calculated for foci number analysis in MEFs
Two-way anova with Tukey's multiple comparisons test

**y-tubulin**

| **0** | Summary | Adjusted P Value |
|---|---|---|
| +/+ vs. +/DG | ns | 0.9466 |
| +/+ vs. DG/DG | ns | 0.2311 |
| +/DG vs. DG/DG | * | 0.0147 |
| **1** | | |
| +/+ vs. +/DG | ns | 0.1131 |
| +/+ vs. DG/DG | ns | 0.6491 |
| +/DG vs. DG/DG | ns | 0.28 |
| **2** | | |
| +/+ vs. +/DG | ns | 0.1851 |
| +/+ vs. DG/DG | ns | 0.1096 |
| +/DG vs. DG/DG | **** | <0.0001 |
| **3** | | |
| +/+ vs. +/DG | ns | 0.998 |
| +/+ vs. DG/DG | ns | 0.742 |
| +/DG vs. DG/DG | ns | 0.5845 |
| **4** | | |
| +/+ vs. +/DG | ns | 0.877 |
| +/+ vs. DG/DG | ns | 0.8329 |
| +/DG vs. DG/DG | ns | 0.9867 |
| **>4** | | |
| +/+ vs. +/DG | ns | 0.9998 |
| +/+ vs. DG/DG | ns | 0.9992 |
| +/DG vs. DG/DG | ns | 0.9964 |

**CEP164**

| **0** | Summary | Adjusted P Value |
|---|---|---|
| +/+ vs. +/DG | ns | 0.9998 |
| +/+ vs. DG/DG | **** | <0.0001 |
| +/DG vs. DG/DG | **** | <0.0001 |
| **1** | | |
| +/+ vs. +/DG | ns | 0.9886 |
| +/+ vs. DG/DG | **** | <0.0001 |
| +/DG vs. DG/DG | **** | <0.0001 |
| **2** | | |
| +/+ vs. +/DG | ns | 0.9494 |
| +/+ vs. DG/DG | ns | 0.669 |
| +/DG vs. DG/DG | ns | 0.2182 |
| **3** | | |
| +/+ vs. +/DG | ns | 0.9997 |
| +/+ vs. DG/DG | ns | 0.9957 |
| +/DG vs. DG/DG | ns | 0.986 |
| **4** | | |
| +/+ vs. +/DG | ns | 0.998 |
| +/+ vs. DG/DG | ns | 0.9957 |
| +/DG vs. DG/DG | ns | 0.9989 |
| **>4** | | |
| +/+ vs. +/DG | ns | 0.9942 |
| +/+ vs. DG/DG | ns | 0.9946 |
| +/DG vs. DG/DG | ns | >0.9999 |

**Acetylated Tubulin**

| **not ciliated** | Summary | Adjusted P Value |
|---|---|---|
| +/+ vs. +/DG | ns | 0.9293 |
| +/+ vs. DG/DG | **** | <0.0001 |
| +/DG vs. DG/DG | **** | <0.0001 |
| **ciliated** | | |
| +/+ vs. +/DG | ns | 0.9293 |
| +/+ vs. DG/DG | **** | <0.0001 |
| +/DG vs. DG/DG | **** | <0.0001 |

### FIGURE 2D

Statistics calculated for foci number analysis in RPE1 cells
Two-way anova with Tukey's multiple comparisons test

**CEP135**

| **0** | Summary | Adjusted P Value |
|---|---|---|
| +/+ vs. DG #1 | ns | 0.8511 |
| +/+ vs. DG #2 | ns | 0.9992 |
| DG #1 vs. DG #2 | ns | 0.8693 |
| **1** | | |
| +/+ vs. DG #1 | ns | 0.222 |
| +/+ vs. DG #2 | ns | 0.5509 |
| DG #1 vs. DG #2 | ns | 0.7978 |
| **2** | | |
| +/+ vs. DG #1 | ns | 0.727 |
| +/+ vs. DG #2 | ns | 0.3784 |
| DG #1 vs. DG #2 | ns | 0.8295 |
| **3** | | |
| +/+ vs. DG #1 | ns | 0.5612 |
| +/+ vs. DG #2 | ns | 0.5045 |
| DG #1 vs. DG #2 | ns | 0.9952 |
| **4** | | |
| +/+ vs. DG #1 | ns | 0.6812 |
| +/+ vs. DG #2 | ns | 0.6012 |
| DG #1 vs. DG #2 | ns | 0.9909 |
| **>4** | | |
| +/+ vs. DG #1 | ns | 0.999 |
| +/+ vs. DG #2 | ns | 0.9617 |
| DG #1 vs. DG #2 | ns | 0.9729 |

**CEP164**

| **0** | Summary | Adjusted P Value |
|---|---|---|
| +/+ vs. DG #1 | **** | <0.0001 |
| +/+ vs. DG #2 | **** | <0.0001 |
| DG #1 vs. DG #2 | ns | 0.1508 |
| **1** | | |
| +/+ vs. DG #1 | **** | <0.0001 |
| +/+ vs. DG #2 | **** | <0.0001 |
| DG #1 vs. DG #2 | ns | 0.1503 |
| **2** | | |
| +/+ vs. DG #1 | ns | 0.9982 |
| +/+ vs. DG #2 | ns | 0.9961 |
| DG #1 vs. DG #2 | ns | 0.9996 |
| **3** | | |
| +/+ vs. DG #1 | ns | >0.9999 |
| +/+ vs. DG #2 | ns | >0.9999 |
| DG #1 vs. DG #2 | ns | >0.9999 |
| **4** | | |
| +/+ vs. DG #1 | ns | >0.9999 |
| +/+ vs. DG #2 | ns | >0.9999 |
| DG #1 vs. DG #2 | ns | >0.9999 |
| **>4** | | |
| +/+ vs. DG #1 | ns | >0.9999 |
| +/+ vs. DG #2 | ns | 0.9997 |
| DG #1 vs. DG #2 | ns | 0.9997 |

**Acetylated Tubulin**

| **not ciliated** | Summary | Adjusted P Value |
|---|---|---|
| +/+ vs. DG #1 | **** | <0.0001 |
| +/+ vs. DG #2 | **** | <0.0001 |
| DG #1 vs. DG #2 | ns | 0.9957 |
| **ciliated** | | |
| +/+ vs. DG #1 | **** | <0.0001 |
| +/+ vs. DG #2 | **** | <0.0001 |
| DG #1 vs. DG #2 | ns | 0.9957 |

#### FIGURE 3A

Statistics calculated for Centriole defects analysis in MEFs
Two-way anova with Tukey's multiple comparisons test

| **normal** | Summary | Adjusted P Value |
| --- | --- | --- |
| +/+ vs. +/DG | ns | 0.8803 |
| +/+ vs. DG/DG | **** | <0.0001 |
| +/DG vs. DG/DG | **** | <0.0001 |
| **short** | | |
| +/+ vs. +/DG | ns | 0.906 |
| +/+ vs. DG/DG | **** | <0.0001 |
| +/DG vs. DG/DG | **** | <0.0001 |
| **broken** | | |
| +/+ vs. +/DG | ns | 0.9545 |
| +/+ vs. DG/DG | **** | <0.0001 |
| +/DG vs. DG/DG | **** | <0.0001 |
| **broken proximal** | | |
| +/+ vs. +/DG | ns | 0.9352 |
| +/+ vs. DG/DG | ns | >0.9999 |
| +/DG vs. DG/DG | ns | 0.9046 |

Statistics calculated for longest MT/ Shortest MT ratio analysis
One-way anova with Tukey's multiple comparisons test

| | | |
| --- | --- | --- |
| +/+ vs. +/DG | ns | 0.8859 |
| +/+ vs. DG/DG | **** | <0.0001 |
| +/DG vs. DG/DG | **** | <0.0001 |

#### FIGURE 3B

Statistics calculated for Centriole defects analysis in RPE1
Two-way anova with Sidak's multiple comparisons test

| +/+ vs. DG/Dg | Summary | Adjusted P Value |
| --- | --- | --- |
| normal | ** | 0.0023 |
| short | ns | 0.6561 |
| broken | ** | 0.0063 |

Statistics calculated for longest MT/ Shortest MT ratio analysis
Unpaired t test with Welch's correction

| | Summary | Adjusted P Value |
| --- | --- | --- |
| +/+ vs. DG/Dg | *** | 0.0001 |

#### FIGURE 3C

Statistics calculated for foci number analysis in RPE1
Two-way anova with Tukey's multiple comparisons test

**SAS6**

| | Summary | Adjusted P Value |
|---|---|---|
| **0** | | |
| +/+ vs. DG #1 | ns | 0.9855 |
| +/+ vs. DG #2 | ns | 0.2664 |
| DG #1 vs. DG #2 | ns | 0.3448 |
| **1** | | |
| +/+ vs. DG #1 | ns | 0.729 |
| +/+ vs. DG #2 | ns | 0.7516 |
| DG #1 vs. DG #2 | ns | 0.9992 |
| **2** | | |
| +/+ vs. DG #1 | ns | 0.4881 |
| +/+ vs. DG #2 | ns | 0.9384 |
| DG #1 vs. DG #2 | ns | 0.7001 |
| **3** | | |
| +/+ vs. DG #1 | ns | 0.9505 |
| +/+ vs. DG #2 | ns | 0.8742 |
| DG #1 vs. DG #2 | ns | 0.9802 |
| **4** | | |
| +/+ vs. DG #1 | ns | 0.9951 |
| +/+ vs. DG #2 | ns | 0.9583 |
| DG #1 vs. DG #2 | ns | 0.9816 |
| **>4** | | |
| +/+ vs. DG #1 | ns | 0.9872 |
| +/+ vs. DG #2 | ns | 0.9119 |
| DG #1 vs. DG #2 | ns | 0.9645 |

**CPAP**

| | Summary | Adjusted P Value |
|---|---|---|
| **0** | | |
| +/+ vs. DG #1 | ns | 0.5212 |
| +/+ vs. DG #2 | ns | 0.9751 |
| DG #1 vs. DG #2 | ns | 0.6534 |
| **1** | | |
| +/+ vs. DG #1 | ns | 0.9221 |
| +/+ vs. DG #2 | ns | 0.9765 |
| DG #1 vs. DG #2 | ns | 0.8254 |
| **2** | | |
| +/+ vs. DG #1 | * | 0.0204 |
| +/+ vs. DG #2 | * | 0.0428 |
| DG #1 vs. DG #2 | ns | 0.9473 |
| **3** | | |
| +/+ vs. DG #1 | ns | 0.171 |
| +/+ vs. DG #2 | ns | 0.2454 |
| DG #1 vs. DG #2 | ns | 0.9763 |
| **4** | | |
| +/+ vs. DG #1 | ns | 0.9848 |
| +/+ vs. DG #2 | ns | 0.9657 |
| DG #1 vs. DG #2 | ns | 0.9081 |
| **>4** | | |
| +/+ vs. DG #1 | ns | 0.8996 |
| +/+ vs. DG #2 | ns | 0.9767 |
| DG #1 vs. DG #2 | ns | 0.9709 |

**CEP295**

| | Summary | Adjusted P Value |
|---|---|---|
| **0** | | |
| +/+ vs. DG #1 | ns | 0.1929 |
| +/+ vs. DG #2 | ns | 0.9387 |
| DG #1 vs. DG #2 | ns | 0.3344 |
| **1** | | |
| +/+ vs. DG #1 | ns | 0.1316 |
| +/+ vs. DG #2 | * | 0.0329 |
| DG #1 vs. DG #2 | ns | 0.7961 |
| **2** | | |
| +/+ vs. DG #1 | **** | <0.0001 |
| +/+ vs. DG #2 | **** | <0.0001 |
| DG #1 vs. DG #2 | ** | 0.0038 |
| **3** | | |
| +/+ vs. DG #1 | *** | 0.0003 |
| +/+ vs. DG #2 | ns | 0.2543 |
| DG #1 vs. DG #2 | * | 0.0217 |
| **4** | | |
| +/+ vs. DG #1 | ns | 0.4065 |
| +/+ vs. DG #2 | ns | 0.5286 |
| DG #1 vs. DG #2 | ns | 0.976 |
| **>4** | | |
| +/+ vs. DG #1 | ns | 0.9926 |
| +/+ vs. DG #2 | ns | 0.8965 |
| DG #1 vs. DG #2 | ns | 0.9419 |

**CEP120**

| | Summary | Adjusted P Value |
|---|---|---|
| **0** | | |
| +/+ vs. DG #1 | ns | >0.9999 |
| +/+ vs. DG #2 | ns | 0.9956 |
| DG #1 vs. DG #2 | ns | 0.9956 |
| **1** | | |
| +/+ vs. DG #1 | ns | 0.5718 |
| +/+ vs. DG #2 | ns | 0.0853 |
| DG #1 vs. DG #2 | ns | 0.4719 |
| **2** | | |
| +/+ vs. DG #1 | ns | 0.5501 |
| +/+ vs. DG #2 | ns | 0.8377 |
| DG #1 vs. DG #2 | ns | 0.8799 |
| **3** | | |
| +/+ vs. DG #1 | ns | 0.8613 |
| +/+ vs. DG #2 | ns | 0.6204 |
| DG #1 vs. DG #2 | ns | 0.9093 |
| **4** | | |
| +/+ vs. DG #1 | ns | 0.566 |
| +/+ vs. DG #2 | ns | 0.7462 |
| DG #1 vs. DG #2 | ns | 0.9538 |
| **>4** | | |
| +/+ vs. DG #1 | ns | 0.8627 |
| +/+ vs. DG #2 | ns | 0.578 |
| DG #1 vs. DG #2 | ns | 0.8779 |

**CP110**

| | Summary | Adjusted P Value |
|---|---|---|
| **0** | | |
| +/+ vs. DG #1 | ns | 0.9948 |
| +/+ vs. DG #2 | ns | 0.8885 |
| DG #1 vs. DG #2 | ns | 0.9286 |
| **1** | | |
| +/+ vs. DG #1 | ns | 0.691 |
| +/+ vs. DG #2 | ns | 0.9655 |
| DG #1 vs. DG #2 | ns | 0.8362 |
| **2** | | |
| +/+ vs. DG #1 | ns | 0.5753 |
| +/+ vs. DG #2 | ns | 0.6695 |
| DG #1 vs. DG #2 | ns | 0.9874 |
| **3** | | |
| +/+ vs. DG #1 | ns | 0.8021 |
| +/+ vs. DG #2 | ns | 0.4626 |
| DG #1 vs. DG #2 | ns | 0.8408 |
| **4** | | |
| +/+ vs. DG #1 | ns | 0.4883 |
| +/+ vs. DG #2 | ns | 0.3 |
| DG #1 vs. DG #2 | ns | 0.9337 |
| **>4** | | |
| +/+ vs. DG #1 | ns | 0.8169 |
| +/+ vs. DG #2 | ns | 0.8944 |
| DG #1 vs. DG #2 | ns | 0.9865 |

#### FIGURE 3D

Statistics calculated for foci number analysis in RPE1
Two-way anova with Tukey's multiple comparisons test

**POC5**

| **0** | Summary | Adjusted P Value |
|---|---|---|
| +/+ vs. DG #1 | *** | 0.0008 |
| +/+ vs. DG #2 | ** | 0.0013 |
| DG #1 vs. DG #2 | ns | 0.9863 |
| **1** | | |
| +/+ vs. DG #1 | **** | <0.0001 |
| +/+ vs. DG #2 | **** | <0.0001 |
| DG #1 vs. DG #2 | ns | 0.991 |
| **2** | | |
| +/+ vs. DG #1 | **** | <0.0001 |
| +/+ vs. DG #2 | **** | <0.0001 |
| DG #1 vs. DG #2 | ns | 0.9686 |
| **3** | | |
| +/+ vs. DG #1 | ns | 0.7548 |
| +/+ vs. DG #2 | ns | 0.5325 |
| DG #1 vs. DG #2 | ns | 0.9311 |
| **4** | | |
| +/+ vs. DG #1 | ns | 0.6865 |
| +/+ vs. DG #2 | ns | 0.824 |
| DG #1 vs. DG #2 | ns | 0.9699 |
| **>4** | | |
| +/+ vs. DG #1 | ns | 0.9974 |
| +/+ vs. DG #2 | ns | >0.9999 |
| DG #1 vs. DG #2 | ns | 0.9974 |

**Centrin2**

| **0** | Summary | Adjusted P Value |
|---|---|---|
| +/+ vs. DG #1 | *** | 0.0003 |
| +/+ vs. DG #2 | ** | 0.0015 |
| DG #1 vs. DG #2 | ns | 0.8792 |
| **1** | | |
| +/+ vs. DG #1 | **** | <0.0001 |
| +/+ vs. DG #2 | **** | <0.0001 |
| DG #1 vs. DG #2 | ns | 0.9466 |
| **2** | | |
| +/+ vs. DG #1 | **** | <0.0001 |
| +/+ vs. DG #2 | **** | <0.0001 |
| DG #1 vs. DG #2 | ns | 0.7376 |
| **3** | | |
| +/+ vs. DG #1 | ns | 0.653 |
| +/+ vs. DG #2 | ns | 0.2972 |
| DG #1 vs. DG #2 | ns | 0.8123 |
| **4** | | |
| +/+ vs. DG #1 | ns | 0.0732 |
| +/+ vs. DG #2 | ns | 0.1374 |
| DG #1 vs. DG #2 | ns | 0.9532 |
| **>4** | | |
| +/+ vs. DG #1 | ns | 0.9475 |
| +/+ vs. DG #2 | ns | 0.9466 |
| DG #1 vs. DG #2 | ns | >0.9999 |

**C2CD3**

| **0** | Summary | Adjusted P Value |
|---|---|---|
| +/+ vs. DG #1 | **** | <0.0001 |
| +/+ vs. DG #2 | **** | <0.0001 |
| DG #1 vs. DG #2 | ns | 0.8574 |
| **1** | | |
| +/+ vs. DG #1 | **** | <0.0001 |
| +/+ vs. DG #2 | **** | <0.0001 |
| DG #1 vs. DG #2 | ns | 0.955 |
| **2** | | |
| +/+ vs. DG #1 | **** | <0.0001 |
| +/+ vs. DG #2 | **** | <0.0001 |
| DG #1 vs. DG #2 | ns | 0.9838 |
| **3** | | |
| +/+ vs. DG #1 | ns | 0.7662 |
| +/+ vs. DG #2 | ns | 0.9856 |
| DG #1 vs. DG #2 | ns | 0.8544 |
| **4** | | |
| +/+ vs. DG #1 | ns | 0.0657 |
| +/+ vs. DG #2 | ns | 0.0805 |
| DG #1 vs. DG #2 | ns | 0.995 |
| **>4** | | |
| +/+ vs. DG #1 | ns | 0.9991 |
| +/+ vs. DG #2 | ns | 0.9997 |
| DG #1 vs. DG #2 | ns | 0.9999 |

**Talpid3**

| **0** | Summary | Adjusted P Value |
|---|---|---|
| +/+ vs. DG #1 | **** | <0.0001 |
| +/+ vs. DG #2 | **** | <0.0001 |
| DG #1 vs. DG #2 | ns | 0.9969 |
| **1** | | |
| +/+ vs. DG #1 | ** | 0.0071 |
| +/+ vs. DG #2 | ** | 0.0028 |
| DG #1 vs. DG #2 | ns | 0.9356 |
| **2** | | |
| +/+ vs. DG #1 | **** | <0.0001 |
| +/+ vs. DG #2 | **** | <0.0001 |
| DG #1 vs. DG #2 | ns | 0.962 |
| **3** | | |
| +/+ vs. DG #1 | ns | 0.8072 |
| +/+ vs. DG #2 | ns | 0.8256 |
| DG #1 vs. DG #2 | ns | 0.9994 |
| **4** | | |
| +/+ vs. DG #1 | ns | 0.0824 |
| +/+ vs. DG #2 | ns | 0.0853 |
| DG #1 vs. DG #2 | ns | 0.9999 |
| **>4** | | |
| +/+ vs. DG #1 | ns | 0.9971 |
| +/+ vs. DG #2 | ns | 0.9999 |
| DG #1 vs. DG #2 | ns | 0.9982 |

**CEP164**

| **0** | Summary | Adjusted P Value |
|---|---|---|
| +/+ vs. DG #1 | **** | <0.0001 |
| +/+ vs. DG #2 | **** | <0.0001 |
| DG #1 vs. DG #2 | ** | 0.0016 |
| **1** | | |
| +/+ vs. DG #1 | **** | <0.0001 |
| +/+ vs. DG #2 | **** | <0.0001 |
| DG #1 vs. DG #2 | ** | 0.0097 |
| **2** | | |
| +/+ vs. DG #1 | ns | 0.3114 |
| +/+ vs. DG #2 | ns | 0.754 |
| DG #1 vs. DG #2 | ns | 0.7277 |
| **3** | | |
| +/+ vs. DG #1 | ns | 0.9975 |
| +/+ vs. DG #2 | ns | 0.9999 |
| DG #1 vs. DG #2 | ns | 0.9984 |
| **4** | | |
| +/+ vs. DG #1 | ns | 0.9988 |
| +/+ vs. DG #2 | ns | >0.9999 |
| DG #1 vs. DG #2 | ns | 0.9988 |
| **>4** | | |
| +/+ vs. DG #1 | ns | >0.9999 |
| +/+ vs. DG #2 | ns | >0.9999 |
| DG #1 vs. DG #2 | ns | >0.9999 |

#### FIGURE 4C

Statistics calculated for foci number analysis in DLD1
Two-way anova with Tukey's multiple comparisons test

**Centrin2**

| **0** | Summary | Adjusted P Value |
|---|---|---|
| +/+ vs. -/- | **** | <0.0001 |
| +/+ vs. -/- WT dox2d | ns | 0.9256 |
| +/+ vs. -/- WT dox4d | ns | 0.9771 |
| +/+ vs. -/- DG dox2d | **** | <0.0001 |
| +/+ vs. -/- DG dox4d | **** | <0.0001 |
| -/- vs. -/- WT dox2d | **** | <0.0001 |
| -/- vs. -/- WT dox4d | **** | <0.0001 |
| -/- vs. -/- DG dox2d | **** | <0.0001 |
| -/- vs. -/- DG dox4d | **** | <0.0001 |
| -/- WT dox2d vs. -/- WT dox4d | ns | >0.9999 |
| -/- WT dox2d vs. -/- DG dox2d | *** | 0.0002 |
| -/- WT dox2d vs. -/- DG dox4d | **** | <0.0001 |
| -/- WT dox4d vs. -/- DG dox2d | *** | 0.0001 |
| -/- WT dox4d vs. -/- DG dox4d | **** | <0.0001 |
| -/- DG dox2d vs. -/- DG dox4d | ns | 0.9994 |

| **1** | | |
|---|---|---|
| +/+ vs. -/- | **** | <0.0001 |
| +/+ vs. -/- WT dox2d | ns | 0.1192 |
| +/+ vs. -/- WT dox4d | ns | 0.0596 |
| +/+ vs. -/- DG dox2d | **** | <0.0001 |
| +/+ vs. -/- DG dox4d | **** | <0.0001 |
| -/- vs. -/- WT dox2d | ** | 0.0095 |
| -/- vs. -/- WT dox4d | * | 0.0223 |
| -/- vs. -/- DG dox2d | ns | 0.1627 |
| -/- vs. -/- DG dox4d | ns | 0.7117 |
| -/- WT dox2d vs. -/- WT dox4d | ns | 0.9999 |
| -/- WT dox2d vs. -/- DG dox2d | **** | <0.0001 |
| -/- WT dox2d vs. -/- DG dox4d | ** | 0.0013 |
| -/- WT dox4d vs. -/- DG dox2d | *** | 0.0002 |
| -/- WT dox4d vs. -/- DG dox4d | ** | 0.0029 |
| -/- DG dox2d vs. -/- DG dox4d | ns | 0.9603 |

| **2** | | |
|---|---|---|
| +/+ vs. -/- | **** | <0.0001 |
| +/+ vs. -/- WT dox2d | ns | >0.9999 |
| +/+ vs. -/- WT dox4d | ns | 0.4855 |
| +/+ vs. -/- DG dox2d | **** | <0.0001 |
| +/+ vs. -/- DG dox4d | **** | <0.0001 |
| -/- vs. -/- WT dox2d | **** | <0.0001 |
| -/- vs. -/- WT dox4d | **** | <0.0001 |
| -/- vs. -/- DG dox2d | ns | 0.116 |
| -/- vs. -/- DG dox4d | * | 0.0103 |
| -/- WT dox2d vs. -/- WT dox4d | ns | 0.7696 |
| -/- WT dox2d vs. -/- DG dox2d | **** | <0.0001 |
| -/- WT dox2d vs. -/- DG dox4d | **** | <0.0001 |
| -/- WT dox4d vs. -/- DG dox2d | **** | <0.0001 |
| -/- WT dox4d vs. -/- DG dox4d | **** | <0.0001 |
| -/- DG dox2d vs. -/- DG dox4d | ns | 0.9725 |

| **3** | Summary | Adjusted P Value |
|---|---|---|
| +/+ vs. -/- | * | 0.0207 |
| +/+ vs. -/- WT dox2d | ns | >0.9999 |
| +/+ vs. -/- WT dox4d | ns | 0.9877 |
| +/+ vs. -/- DG dox2d | ns | 0.9693 |
| +/+ vs. -/- DG dox4d | ns | 0.6962 |
| -/- vs. -/- WT dox2d | ns | 0.145 |
| -/- vs. -/- WT dox4d | * | 0.0319 |
| -/- vs. -/- DG dox2d | ns | 0.5527 |
| -/- vs. -/- DG dox4d | ns | 0.9136 |
| -/- WT dox2d vs. -/- WT dox4d | ns | 0.9952 |
| -/- WT dox2d vs. -/- DG dox2d | ns | 0.9846 |
| -/- WT dox2d vs. -/- DG dox4d | ns | 0.8218 |
| -/- WT dox4d vs. -/- DG dox2d | ns | 0.8399 |
| -/- WT dox4d vs. -/- DG dox4d | ns | 0.5055 |
| -/- DG dox2d vs. -/- DG dox4d | ns | 0.9934 |

| **4** | | |
|---|---|---|
| +/+ vs. -/- | **** | <0.0001 |
| +/+ vs. -/- WT dox2d | ns | 0.0966 |
| +/+ vs. -/- WT dox4d | ns | 0.3195 |
| +/+ vs. -/- DG dox2d | *** | 0.0006 |
| +/+ vs. -/- DG dox4d | ** | 0.0075 |
| -/- vs. -/- WT dox2d | ns | 0.4608 |
| -/- vs. -/- WT dox4d | ns | 0.1644 |
| -/- vs. -/- DG dox2d | ns | >0.9999 |
| -/- vs. -/- DG dox4d | ns | 0.9496 |
| -/- WT dox2d vs. -/- WT dox4d | ns | 0.996 |
| -/- WT dox2d vs. -/- DG dox2d | ns | 0.7075 |
| -/- WT dox2d vs. -/- DG dox4d | ns | 0.9695 |
| -/- WT dox4d vs. -/- DG dox2d | ns | 0.3913 |
| -/- WT dox4d vs. -/- DG dox4d | ns | 0.793 |
| -/- DG dox2d vs. -/- DG dox4d | ns | 0.9867 |

| **>4** | | |
|---|---|---|
| +/+ vs. -/- | ns | 0.6732 |
| +/+ vs. -/- WT dox2d | ns | 0.9847 |
| +/+ vs. -/- WT dox4d | ns | 0.9992 |
| +/+ vs. -/- DG dox2d | ns | 0.999 |
| +/+ vs. -/- DG dox4d | ns | 0.9658 |
| -/- vs. -/- WT dox2d | ns | 0.9975 |
| -/- vs. -/- WT dox4d | ns | 0.9724 |
| -/- vs. -/- DG dox2d | ns | 0.9742 |
| -/- vs. -/- DG dox4d | ns | 0.9995 |
| -/- WT dox2d vs. -/- WT dox4d | ns | 0.9998 |
| -/- WT dox2d vs. -/- DG dox2d | ns | 0.9999 |
| -/- WT dox2d vs. -/- DG dox4d | ns | >0.9999 |
| -/- WT dox4d vs. -/- DG dox2d | ns | >0.9999 |
| -/- WT dox4d vs. -/- DG dox4d | ns | 0.999 |
| -/- DG dox2d vs. -/- DG dox4d | ns | 0.9991 |

#### FIGURE 4D

Statistics calculated for foci number analysis in DLD1
Two-way anova with Tukey's multiple comparisons test

**CEP164**

| 0 | Summary | Adjusted P Value |
|---|---|---|
| +/+ vs. -/- | **** | <0.0001 |
| +/+ vs. -/- WT dox1d | **** | <0.0001 |
| +/+ vs. -/- WT dox2d | *** | 0.0003 |
| +/+ vs. -/- WT dox4d | ns | 0.2477 |
| +/+ vs. -/- DG dox1d | **** | <0.0001 |
| +/+ vs. -/- DG dox2d | **** | <0.0001 |
| +/+ vs. -/- DG dox4d | **** | <0.0001 |
| -/- vs. -/- WT dox1d | **** | <0.0001 |
| -/- vs. -/- WT dox2d | **** | <0.0001 |
| -/- vs. -/- WT dox4d | **** | <0.0001 |
| -/- vs. -/- DG dox1d | ns | >0.9999 |
| -/- vs. -/- DG dox2d | ns | >0.9999 |
| -/- vs. -/- DG dox4d | ** | 0.0013 |
| -/- WT dox1d vs. -/- WT dox2d | * | 0.0347 |
| -/- WT dox1d vs. -/- WT dox4d | **** | <0.0001 |
| -/- WT dox1d vs. -/- DG dox1d | *** | 0.0003 |
| -/- WT dox1d vs. -/- DG dox2d | *** | 0.0002 |
| -/- WT dox1d vs. -/- DG dox4d | ns | 0.9112 |
| -/- WT dox2d vs. -/- WT dox4d | ns | 0.39 |
| -/- WT dox2d vs. -/- DG dox1d | **** | <0.0001 |
| -/- WT dox2d vs. -/- DG dox2d | **** | <0.0001 |
| -/- WT dox2d vs. -/- DG dox4d | *** | 0.0004 |
| -/- WT dox4d vs. -/- DG dox1d | **** | <0.0001 |
| -/- WT dox4d vs. -/- DG dox2d | **** | <0.0001 |
| -/- WT dox4d vs. -/- DG dox4d | **** | <0.0001 |
| -/- DG dox1d vs. -/- DG dox2d | ns | >0.9999 |
| -/- DG dox1d vs. -/- DG dox4d | * | 0.0189 |
| -/- DG dox2d vs. -/- DG dox4d | * | 0.0166 |

| 1 | | |
|---|---|---|
| +/+ vs. -/- | **** | <0.0001 |
| +/+ vs. -/- WT dox1d | **** | <0.0001 |
| +/+ vs. -/- WT dox2d | **** | <0.0001 |
| +/+ vs. -/- WT dox4d | ** | 0.0039 |
| +/+ vs. -/- DG dox1d | **** | <0.0001 |
| +/+ vs. -/- DG dox2d | **** | <0.0001 |
| +/+ vs. -/- DG dox4d | **** | <0.0001 |
| -/- vs. -/- WT dox1d | ** | 0.003 |
| -/- vs. -/- WT dox2d | **** | <0.0001 |
| -/- vs. -/- WT dox4d | **** | <0.0001 |
| -/- vs. -/- DG dox1d | ns | >0.9999 |
| -/- vs. -/- DG dox2d | ns | >0.9999 |
| -/- vs. -/- DG dox4d | ** | 0.0079 |
| -/- WT dox1d vs. -/- WT dox2d | ns | 0.1551 |
| -/- WT dox1d vs. -/- WT dox4d | **** | <0.0001 |
| -/- WT dox1d vs. -/- DG dox1d | * | 0.0482 |
| -/- WT dox1d vs. -/- DG dox2d | * | 0.0208 |
| -/- WT dox1d vs. -/- DG dox4d | ns | >0.9999 |
| -/- WT dox2d vs. -/- WT dox4d | * | 0.0497 |
| -/- WT dox2d vs. -/- DG dox1d | **** | <0.0001 |
| -/- WT dox2d vs. -/- DG dox2d | **** | <0.0001 |
| -/- WT dox2d vs. -/- DG dox4d | ns | 0.0861 |
| -/- WT dox4d vs. -/- DG dox1d | **** | <0.0001 |
| -/- WT dox4d vs. -/- DG dox2d | **** | <0.0001 |
| -/- WT dox4d vs. -/- DG dox4d | **** | <0.0001 |
| -/- DG dox1d vs. -/- DG dox2d | ns | >0.9999 |
| -/- DG dox1d vs. -/- DG dox4d | ns | 0.0884 |
| -/- DG dox2d vs. -/- DG dox4d | * | 0.0411 |

| 2 | | |
|---|---|---|
| +/+ vs. -/- | ns | >0.9999 |
| +/+ vs. -/- WT dox1d | ns | 0.8956 |
| +/+ vs. -/- WT dox2d | ns | 0.5278 |
| +/+ vs. -/- WT dox4d | ns | 0.9999 |
| +/+ vs. -/- DG dox1d | ns | >0.9999 |
| +/+ vs. -/- DG dox2d | ns | >0.9999 |
| +/+ vs. -/- DG dox4d | ns | 0.9999 |
| -/- vs. -/- WT dox1d | ns | 0.8038 |
| -/- vs. -/- WT dox2d | ns | 0.3962 |
| -/- vs. -/- WT dox4d | ns | 0.9981 |
| -/- vs. -/- DG dox1d | ns | >0.9999 |
| -/- vs. -/- DG dox2d | ns | >0.9999 |
| -/- vs. -/- DG dox4d | ns | 0.9985 |
| -/- WT dox1d vs. -/- WT dox2d | ns | 0.9993 |
| -/- WT dox1d vs. -/- WT dox4d | ns | 0.9905 |
| -/- WT dox1d vs. -/- DG dox1d | ns | 0.9594 |
| -/- WT dox1d vs. -/- DG dox2d | ns | 0.8838 |
| -/- WT dox1d vs. -/- DG dox4d | ns | 0.9938 |
| -/- WT dox2d vs. -/- WT dox4d | ns | 0.8422 |
| -/- WT dox2d vs. -/- DG dox1d | ns | 0.7487 |
| -/- WT dox2d vs. -/- DG dox2d | ns | 0.5844 |
| -/- WT dox2d vs. -/- DG dox4d | ns | 0.8816 |
| -/- WT dox4d vs. -/- DG dox1d | ns | >0.9999 |
| -/- WT dox4d vs. -/- DG dox2d | ns | 0.9985 |
| -/- WT dox4d vs. -/- DG dox4d | ns | >0.9999 |
| -/- DG dox1d vs. -/- DG dox2d | ns | >0.9999 |
| -/- DG dox1d vs. -/- DG dox4d | ns | >0.9999 |
| -/- DG dox2d vs. -/- DG dox4d | ns | 0.9987 |

| 3 | Summary | Adjusted P Value |
|---|---|---|
| +/+ vs. -/- | ns | >0.9999 |
| +/+ vs. -/- WT dox1d | ns | >0.9999 |
| +/+ vs. -/- WT dox2d | ns | >0.9999 |
| +/+ vs. -/- WT dox4d | ns | 0.9965 |
| +/+ vs. -/- DG dox1d | ns | >0.9999 |
| +/+ vs. -/- DG dox2d | ns | >0.9999 |
| +/+ vs. -/- DG dox4d | ns | >0.9999 |
| -/- vs. -/- WT dox1d | ns | >0.9999 |
| -/- vs. -/- WT dox2d | ns | >0.9999 |
| -/- vs. -/- WT dox4d | ns | 0.9999 |
| -/- vs. -/- DG dox1d | ns | >0.9999 |
| -/- vs. -/- DG dox2d | ns | >0.9999 |
| -/- vs. -/- DG dox4d | ns | >0.9999 |
| -/- WT dox1d vs. -/- WT dox2d | ns | >0.9999 |
| -/- WT dox1d vs. -/- WT dox4d | ns | >0.9999 |
| -/- WT dox1d vs. -/- DG dox1d | ns | >0.9999 |
| -/- WT dox1d vs. -/- DG dox2d | ns | >0.9999 |
| -/- WT dox1d vs. -/- DG dox4d | ns | >0.9999 |
| -/- WT dox2d vs. -/- WT dox4d | ns | >0.9999 |
| -/- WT dox2d vs. -/- DG dox1d | ns | >0.9999 |
| -/- WT dox2d vs. -/- DG dox2d | ns | >0.9999 |
| -/- WT dox2d vs. -/- DG dox4d | ns | >0.9999 |
| -/- WT dox4d vs. -/- DG dox1d | ns | 0.9981 |
| -/- WT dox4d vs. -/- DG dox2d | ns | 0.9998 |
| -/- WT dox4d vs. -/- DG dox4d | ns | 0.9996 |
| -/- DG dox1d vs. -/- DG dox2d | ns | >0.9999 |
| -/- DG dox1d vs. -/- DG dox4d | ns | >0.9999 |
| -/- DG dox2d vs. -/- DG dox4d | ns | >0.9999 |

| 4 | | |
|---|---|---|
| +/+ vs. -/- | ns | >0.9999 |
| +/+ vs. -/- WT dox1d | ns | >0.9999 |
| +/+ vs. -/- WT dox2d | ns | >0.9999 |
| +/+ vs. -/- WT dox4d | ns | >0.9999 |
| +/+ vs. -/- DG dox1d | ns | >0.9999 |
| +/+ vs. -/- DG dox2d | ns | >0.9999 |
| +/+ vs. -/- DG dox4d | ns | >0.9999 |
| -/- vs. -/- WT dox1d | ns | >0.9999 |
| -/- vs. -/- WT dox2d | ns | >0.9999 |
| -/- vs. -/- WT dox4d | ns | >0.9999 |
| -/- vs. -/- DG dox1d | ns | >0.9999 |
| -/- vs. -/- DG dox2d | ns | >0.9999 |
| -/- vs. -/- DG dox4d | ns | >0.9999 |
| -/- WT dox1d vs. -/- WT dox2d | ns | >0.9999 |
| -/- WT dox1d vs. -/- WT dox4d | ns | >0.9999 |
| -/- WT dox1d vs. -/- DG dox1d | ns | >0.9999 |
| -/- WT dox1d vs. -/- DG dox2d | ns | >0.9999 |
| -/- WT dox1d vs. -/- DG dox4d | ns | >0.9999 |
| -/- WT dox2d vs. -/- WT dox4d | ns | >0.9999 |
| -/- WT dox2d vs. -/- DG dox1d | ns | >0.9999 |
| -/- WT dox2d vs. -/- DG dox2d | ns | >0.9999 |
| -/- WT dox2d vs. -/- DG dox4d | ns | >0.9999 |
| -/- WT dox4d vs. -/- DG dox1d | ns | >0.9999 |
| -/- WT dox4d vs. -/- DG dox2d | ns | >0.9999 |
| -/- WT dox4d vs. -/- DG dox4d | ns | >0.9999 |
| -/- DG dox1d vs. -/- DG dox2d | ns | >0.9999 |
| -/- DG dox1d vs. -/- DG dox4d | ns | >0.9999 |
| -/- DG dox2d vs. -/- DG dox4d | ns | >0.9999 |

| >4 | | |
|---|---|---|
| +/+ vs. -/- | ns | >0.9999 |
| +/+ vs. -/- WT dox1d | ns | >0.9999 |
| +/+ vs. -/- WT dox2d | ns | >0.9999 |
| +/+ vs. -/- WT dox4d | ns | >0.9999 |
| +/+ vs. -/- DG dox1d | ns | >0.9999 |
| +/+ vs. -/- DG dox2d | ns | >0.9999 |
| +/+ vs. -/- DG dox4d | ns | >0.9999 |
| -/- vs. -/- WT dox1d | ns | >0.9999 |
| -/- vs. -/- WT dox2d | ns | >0.9999 |
| -/- vs. -/- WT dox4d | ns | >0.9999 |
| -/- vs. -/- DG dox1d | ns | >0.9999 |
| -/- vs. -/- DG dox2d | ns | >0.9999 |
| -/- vs. -/- DG dox4d | ns | >0.9999 |
| -/- WT dox1d vs. -/- WT dox2d | ns | >0.9999 |
| -/- WT dox1d vs. -/- WT dox4d | ns | >0.9999 |
| -/- WT dox1d vs. -/- DG dox1d | ns | >0.9999 |
| -/- WT dox1d vs. -/- DG dox2d | ns | >0.9999 |
| -/- WT dox1d vs. -/- DG dox4d | ns | >0.9999 |
| -/- WT dox2d vs. -/- WT dox4d | ns | >0.9999 |
| -/- WT dox2d vs. -/- DG dox1d | ns | >0.9999 |
| -/- WT dox2d vs. -/- DG dox2d | ns | >0.9999 |
| -/- WT dox2d vs. -/- DG dox4d | ns | >0.9999 |
| -/- WT dox4d vs. -/- DG dox1d | ns | >0.9999 |
| -/- WT dox4d vs. -/- DG dox2d | ns | >0.9999 |
| -/- WT dox4d vs. -/- DG dox4d | ns | >0.9999 |
| -/- DG dox1d vs. -/- DG dox2d | ns | >0.9999 |
| -/- DG dox1d vs. -/- DG dox4d | ns | >0.9999 |
| -/- DG dox2d vs. -/- DG dox4d | ns | >0.9999 |

#### FIGURE 4G

Statistics calculated for Centriole defects analysis in DLD1
Two-way anova with Tukey's multiple comparisons test

| **normal** | Summary | Adjusted P Value |
|---|---|---|
| +/+ vs. -/- | **** | <0.0001 |
| +/+ vs. -/- WT dox2d | *** | 0.0008 |
| +/+ vs. -/- WT dox4d | **** | <0.0001 |
| +/+ vs. -/- DG dox2d | **** | <0.0001 |
| +/+ vs. -/- DG dox4d | **** | <0.0001 |
| -/- vs. -/- WT dox2d | **** | <0.0001 |
| -/- vs. -/- WT dox4d | *** | 0.0001 |
| -/- vs. -/- DG dox2d | ns | 0.5504 |
| -/- vs. -/- DG dox4d | ns | 0.9923 |
| -/- WT dox2d vs. -/- WT dox4d | * | 0.0417 |
| -/- WT dox2d vs. -/- DG dox2d | **** | <0.0001 |
| -/- WT dox2d vs. -/- DG dox4d | **** | <0.0001 |
| -/- WT dox4d vs. -/- DG dox2d | ns | 0.0785 |
| -/- WT dox4d vs. -/- DG dox4d | *** | 0.0001 |
| -/- DG dox2d vs. -/- DG dox4d | ns | 0.3269 |

| **short** | | |
|---|---|---|
| +/+ vs. -/- | **** | <0.0001 |
| +/+ vs. -/- WT dox2d | ns | 0.8974 |
| +/+ vs. -/- WT dox4d | ns | 0.4828 |
| +/+ vs. -/- DG dox2d | * | 0.0183 |
| +/+ vs. -/- DG dox4d | **** | <0.0001 |
| -/- vs. -/- WT dox2d | ** | 0.006 |
| -/- vs. -/- WT dox4d | ** | 0.0096 |
| -/- vs. -/- DG dox2d | ns | 0.6949 |
| -/- vs. -/- DG dox4d | ns | 0.8341 |
| -/- WT dox2d vs. -/- WT dox4d | ns | 0.9937 |
| -/- WT dox2d vs. -/- DG dox2d | ns | 0.2992 |
| -/- WT dox2d vs. -/- DG dox4d | *** | 0.0006 |
| -/- WT dox4d vs. -/- DG dox2d | ns | 0.4968 |
| -/- WT dox4d vs. -/- DG dox4d | *** | 0.0008 |
| -/- DG dox2d vs. -/- DG dox4d | ns | 0.1703 |

| **broken** | | |
|---|---|---|
| +/+ vs. -/- | **** | <0.0001 |
| +/+ vs. -/- WT dox2d | ns | 0.293 |
| +/+ vs. -/- WT dox4d | ns | 0.1703 |
| +/+ vs. -/- DG dox2d | **** | <0.0001 |
| +/+ vs. -/- DG dox4d | **** | <0.0001 |
| -/- vs. -/- WT dox2d | **** | <0.0001 |
| -/- vs. -/- WT dox4d | **** | <0.0001 |
| -/- vs. -/- DG dox2d | ns | 0.9975 |
| -/- vs. -/- DG dox4d | ns | 0.9272 |
| -/- WT dox2d vs. -/- WT dox4d | ns | >0.9999 |
| -/- WT dox2d vs. -/- DG dox2d | ** | 0.0012 |
| -/- WT dox2d vs. -/- DG dox4d | ** | 0.0048 |
| -/- WT dox4d vs. -/- DG dox2d | *** | 0.0004 |
| -/- WT dox4d vs. -/- DG dox4d | ** | 0.0018 |
| -/- DG dox2d vs. -/- DG dox4d | ns | 0.9971 |

| **long** | Summary | Adjusted P Value |
|---|---|---|
| +/+ vs. -/- | ns | >0.9999 |
| +/+ vs. -/- WT dox2d | ns | >0.9999 |
| +/+ vs. -/- WT dox4d | ns | 0.7879 |
| +/+ vs. -/- DG dox2d | ns | >0.9999 |
| +/+ vs. -/- DG dox4d | ns | 0.9999 |
| -/- vs. -/- WT dox2d | ns | >0.9999 |
| -/- vs. -/- WT dox4d | ns | 0.7879 |
| -/- vs. -/- DG dox2d | ns | >0.9999 |
| -/- vs. -/- DG dox4d | ns | 0.9999 |
| -/- WT dox2d vs. -/- WT dox4d | ns | 0.9246 |
| -/- WT dox2d vs. -/- DG dox2d | ns | >0.9999 |
| -/- WT dox2d vs. -/- DG dox4d | ns | >0.9999 |
| -/- WT dox4d vs. -/- DG dox2d | ns | 0.9359 |
| -/- WT dox4d vs. -/- DG dox4d | ns | 0.9435 |
| -/- DG dox2d vs. -/- DG dox4d | ns | >0.9999 |

| **filament** | | |
|---|---|---|
| +/+ vs. -/- | ns | >0.9999 |
| +/+ vs. -/- WT dox2d | ns | 0.8918 |
| +/+ vs. -/- WT dox4d | ns | 0.0664 |
| +/+ vs. -/- DG dox2d | ns | >0.9999 |
| +/+ vs. -/- DG dox4d | ns | >0.9999 |
| -/- vs. -/- WT dox2d | ns | 0.816 |
| -/- vs. -/- WT dox4d | * | 0.0418 |
| -/- vs. -/- DG dox2d | ns | >0.9999 |
| -/- vs. -/- DG dox4d | ns | >0.9999 |
| -/- WT dox2d vs. -/- WT dox4d | ns | 0.673 |
| -/- WT dox2d vs. -/- DG dox2d | ns | 0.8663 |
| -/- WT dox2d vs. -/- DG dox4d | ns | 0.8968 |
| -/- WT dox4d vs. -/- DG dox2d | ns | 0.0897 |
| -/- WT dox4d vs. -/- DG dox4d | ns | 0.1079 |
| -/- DG dox2d vs. -/- DG dox4d | ns | >0.9999 |

### FIGURE 4H

Statistics calculated for longest MT/ Shortest MT ratio analysis in DLD1
One-way anova with Tukey's multiple comparisons test

| | Summary | Adjusted P Value |
|---|---|---|
| +/+ vs. -/- | *** | 0.0004 |
| +/+ vs. -/- WT dox2d | ns | 0.9613 |
| +/+ vs. -/- WT dox4d | ns | 0.4802 |
| +/+ vs. -/- DG dox2d | *** | 0.0007 |
| +/+ vs. -/- DG dox4d | **** | <0.0001 |
| -/- vs. -/- WT dox2d | ** | 0.01 |
| -/- vs. -/- WT dox4d | * | 0.0106 |
| -/- vs. -/- DG dox2d | ns | 0.9991 |
| -/- vs. -/- DG dox4d | ns | 0.7979 |
| -/- WT dox2d vs. -/- WT dox4d | ns | 0.9759 |
| -/- WT dox2d vs. -/- DG dox2d | * | 0.0182 |
| -/- WT dox2d vs. -/- DG dox4d | **** | <0.0001 |
| -/- WT dox4d vs. -/- DG dox2d | * | 0.019 |
| -/- WT dox4d vs. -/- DG dox4d | **** | <0.0001 |
| -/- DG dox2d vs. -/- DG dox4d | ns | 0.4458 |

**FIGURE 5E**

Statistics calculated for HYLS1 intensity upon CEP120 siRNA

One-way anova with Tukey's multiple comparisons test

| | Summary | Adjusted P Value |
|---|---|---|
| -/- WT no siRNA vs. -/- WT CEP120 siRNA | ** | 0.001 |
| -/- WT no siRNA vs. -/- DG no siRNA | *** | 0.0005 |
| -/- WT no siRNA vs. -/- DG CEP120 siRNA | **** | <0.0001 |
| -/- WT CEP120 siRNA vs. -/- DG no siRNA | ns | 0.8644 |
| -/- WT CEP120 siRNA vs. -/- DG CEP120 siRNA | ns | 0.0872 |
| -/- DG no siRNA vs. -/- DG CEP120 siRNA | ns | 0.2522 |

**FIGURE 5G**

Statistics calculated for HYLS1 interaction with CEP120 analysis

Unpaired t test with Welch's correction

| | Summary | Adjusted P Value |
|---|---|---|
| -/- WT-mCherry vs. -/- DG-mCherry | ** | 0.0039 |

**FIGURE S1B**
Statistics calculated for E14.5 pup size
one-way anova with Tukey's multiple comparisons test

| **E14.5 Size** | Summary | Adjusted P Value |
|---|---|---|
| +/+ vs. +/DG | ns | 0.4171 |
| +/+ vs. DG/DG | * | 0.0214 |
| +/DG vs. DG/DG | ns | 0.0908 |

| **E14.5 weight** | Summary | Adjusted P Value |
|---|---|---|
| +/+ vs. +/DG | ns | 0.5838 |
| +/+ vs. DG/DG | ns | 0.7074 |
| +/DG vs. DG/DG | ns | 0.977 |

**FIGURE S1C**
Statistics calculated for E18.5 pup size
one-way anova with Tukey's multiple comparisons test

| **E18.5 Size** | Summary | Adjusted P Value |
|---|---|---|
| +/+ vs. +/DG | ns | 0.7706 |
| +/+ vs. DG/DG | * | 0.0334 |
| +/DG vs. DG/DG | ** | 0.003 |

| **E18.5 weight** | Summary | Adjusted P Value |
|---|---|---|
| +/+ vs. +/DG | ns | 0.9115 |
| +/+ vs. DG/DG | ns | 0.063 |
| +/DG vs. DG/DG | * | 0.0119 |

**FIGURE S1D**
Statistics calculated for P0 pup size
one-way anova with Tukey's multiple comparisons test

| **P0 Size** | Summary | Adjusted P Value |
|---|---|---|
| +/+ vs. +/DG | ns | 0.4238 |
| +/+ vs. DG/DG | **** | <0.0001 |
| +/DG vs. DG/DG | **** | <0.0001 |

| **P0 Weight** | Summary | Adjusted P Value |
|---|---|---|
| +/+ vs. +/DG | ns | 0.6782 |
| +/+ vs. DG/DG | **** | <0.0001 |
| +/DG vs. DG/DG | **** | <0.0001 |

**FIGURE S1E**
Statistics calculated for brain size
one-way anova with Tukey's multiple comparisons test

| **Cerebral Cortex Size** | Summary | Adjusted P Value |
|---|---|---|
| +/+ vs. +/DG | ns | 0.3657 |
| +/+ vs. DG/DG | ** | 0.0074 |
| +/DG vs. DG/DG | ns | 0.0596 |

| **Midbrain Size** | Summary | Adjusted P Value |
|---|---|---|
| +/+ vs. +/DG | ns | 0.8556 |
| +/+ vs. DG/DG | ns | 0.5134 |
| +/DG vs. DG/DG | ns | 0.1431 |

**FIGURE S1H**
Statistics calculated for centriole microtubule length analysis in the kidney
Unpaired t test with Welch's correction

| **Longest MT** | Summary | Adjusted P Value |
|---|---|---|
| Control vs. DG/DG | ns | 0.0934 |
| **Shortest MT** | | |
| Control vs. DG/DG | **** | <0.0001 |

**FIGURE S1I**
Statistics calculated for centriole microtubule length analysis in the brain
Unpaired t test with Welch's correction

| **Longest MT** | Summary |
|---|---|
| Control vs. DG/DG | ns |
| **Shortest MT** | |
| Control vs. DG/DG | ** |

#### FIGURE S3B

Statistics calculated for cilia length analysis in MEFs
one-way anova with Tukey's multiple comparisons test

| | Summary | Adjusted P Value |
|---|---|---|
| +/+ vs. +/DG | ns | 0.6647 |
| +/+ vs. DG/DG | ns | 0.9785 |
| +/DG vs. DG/DG | ns | 0.3227 |

#### FIGURE S3C

Statistics calculated for Smoothened analysis in MEFS
two-way anova with Tukey's multiple comparisons test

| **Smo negative** | Summary | Adjusted P Value |
|---|---|---|
| -SAG +/+ vs. +SAG +/+ | **** | <0.0001 |
| -SAG +/+ vs. -SAG +/DG | ns | 0.9996 |
| -SAG +/+ vs.+SAG +/DG | **** | <0.0001 |
| -SAG +/+ vs. -SAG DG/DG | ns | >0.9999 |
| -SAG +/+ vs. +SAG DG/DG | **** | <0.0001 |
| +SAG +/+ vs. -SAG +/DG | **** | <0.0001 |
| +SAG +/+ vs.+SAG +/DG | ns | 0.9998 |
| +SAG +/+ vs. -SAG DG/DG | **** | <0.0001 |
| +SAG +/+ vs. +SAG DG/DG | ns | 0.8527 |
| -SAG +/DG vs. +SAG +/DG | **** | <0.0001 |
| -SAG +/DG vs. -SAG DG/DG | ns | 0.9936 |
| -SAG +/DG vs. +SAG DG/DG | **** | <0.0001 |
| +SAG +/DG vs. -SAG DG/DG | **** | <0.0001 |
| +SAG +/DG vs. +SAG DG/DG | ns | 0.7742 |
| -SAG DG/DG vs. +SAG DG/DG | **** | <0.0001 |

| **Smo positive** | | |
|---|---|---|
| -SAG +/+ vs. +SAG +/+ | **** | <0.0001 |
| -SAG +/+ vs. -SAG +/DG | ns | 0.9996 |
| -SAG +/+ vs.+SAG +/DG | **** | <0.0001 |
| -SAG +/+ vs. -SAG DG/DG | ns | >0.9999 |
| -SAG +/+ vs. +SAG DG/DG | **** | <0.0001 |
| +SAG +/+ vs. -SAG +/DG | **** | <0.0001 |
| +SAG +/+ vs.+SAG +/DG | ns | 0.9998 |
| +SAG +/+ vs. -SAG DG/DG | **** | <0.0001 |
| +SAG +/+ vs. +SAG DG/DG | ns | 0.8527 |
| -SAG +/DG vs. +SAG +/DG | **** | <0.0001 |
| -SAG +/DG vs. -SAG DG/DG | ns | 0.9936 |
| -SAG +/DG vs. +SAG DG/DG | **** | <0.0001 |
| +SAG +/DG vs. -SAG DG/DG | **** | <0.0001 |
| +SAG +/DG vs. +SAG DG/DG | ns | 0.7739 |
| -SAG DG/DG vs. +SAG DG/DG | **** | <0.0001 |

#### FIGURE S3D

Statistics calculated for Cell Cylcle profile analysis
Tukey's multiple comparisons test

| SubG1 | Summary | Adjusted P Value |
|---|---|---|
| wt vs. ko | ns | >0.9999 |
| wt vs. wt 1xHA | ns | >0.9999 |
| wt vs. DG 1xHA | ns | 0.9662 |
| ko vs. wt 1xHA | ns | >0.9999 |
| ko vs. DG 1xHA | ns | 0.9805 |
| wt 1xHA vs. DG 1xHA | ns | 0.9458 |
| G1 | | |
| wt vs. ko | ns | 0.4359 |
| wt vs. wt 1xHA | ns | 0.9978 |
| wt vs. DG 1xHA | ns | 0.0792 |
| ko vs. wt 1xHA | ns | 0.406 |
| ko vs. DG 1xHA | ns | 0.9554 |
| wt 1xHA vs. DG 1xHA | * | 0.0253 |
| S | | |
| wt vs. ko | ns | 0.733 |
| wt vs. wt 1xHA | ns | 0.9995 |
| wt vs. DG 1xHA | ns | 0.219 |
| ko vs. wt 1xHA | ns | 0.5655 |
| ko vs. DG 1xHA | ns | 0.9199 |
| wt 1xHA vs. DG 1xHA | * | 0.0444 |
| G2/M | | |
| wt vs. ko | ns | 0.9929 |
| wt vs. wt 1xHA | ns | 0.9989 |
| wt vs. DG 1xHA | ns | 0.7784 |
| ko vs. wt 1xHA | ns | 0.9673 |
| ko vs. DG 1xHA | ns | 0.5697 |
| wt 1xHA vs. DG 1xHA | ns | 0.7397 |
| Ploidy | | |
| wt vs. ko | ns | 0.9985 |
| wt vs. wt 1xHA | ns | 0.9639 |
| wt vs. DG 1xHA | ns | 0.7158 |
| ko vs. wt 1xHA | ns | 0.9912 |
| ko vs. DG 1xHA | ns | 0.8276 |
| wt 1xHA vs. DG 1xHA | ns | 0.9005 |

#### FIGURE S3F

Statistics calculated for cilia in MCCs analysis
one-way anova with Tukey's multiple comparisons test

| **FOXJ1+** | Summary | Adjusted P Value |
|---|---|---|
| +/+ vs. +/DG | ns | 0.7921 |
| +/+ vs. DG/DG | ns | 0.8609 |
| +/DG vs. DG/DG | ns | 0.9917 |

| **FOXJ1+ MCCs** | Summary | Adjusted P Value |
|---|---|---|
| +/+ vs. +/DG | ns | 0.2957 |
| +/+ vs. DG/DG | ns | 0.684 |
| +/DG vs. DG/DG | ns | 0.7587 |

| **Centrin2/Deup1+ MCCs** | Summary | Adjusted P Value |
|---|---|---|
| +/+ vs. +/DG | ns | 0.8901 |
| +/+ vs. DG/DG | ns | 0.9259 |
| +/DG vs. DG/DG | ns | 0.9959 |

| **Centrin2/CEP164+ MCCs** | Summary | Adjusted P Value |
|---|---|---|
| +/+ vs. +/DG | ns | 0.9789 |
| +/+ vs. DG/DG | ns | 0.9608 |
| +/DG vs. DG/DG | ns | 0.9007 |

### FIGURE S4B

Statistics calculated for centriole foci number analysis in RPE1
one-way anova with Sidak's multiple comparisons test

**CEP135**

+/+ vs. -/-

| | | |
|---|---|---|
| 0 | ns | >0.9999 |
| 1 | ns | >0.9999 |
| 2 | ns | 0.6814 |
| 3 | ns | 0.9555 |
| 4 | ns | 0.9995 |
| >4 | ns | >0.9999 |

**CEP164**

+/+ vs. -/-

| | | |
|---|---|---|
| 0 | **** | <0.0001 |
| 1 | **** | <0.0001 |
| 2 | ns | >0.9999 |
| 3 | ns | >0.9999 |
| 4 | ns | >0.9999 |
| >4 | ns | >0.9999 |

**Acetylated Tubulin**

+/+ vs. -/-

| | | |
|---|---|---|
| not ciliated | **** | <0.0001 |
| ciliated | **** | <0.0001 |

#### FIGURE S5A

Statistics calculated for centriole width and length analysis in MEFs
one-way anova with Tukey's multiple comparisons test

| **Proximal Width** | Summary | Adjusted P Value |
|---|---|---|
| +/+ vs. +/DG | * | 0.0342 |
| +/+ vs. DG/DG | ns | 0.1125 |
| +/DG vs. DG/DG | ns | 0.9362 |

| **Distal Width** | Summary | Adjusted P Value |
|---|---|---|
| +/+ vs. +/DG | *** | 0.0004 |
| +/+ vs. DG/DG | *** | 0.0003 |
| +/DG vs. DG/DG | ns | 0.8412 |

| **Longest MT** | Summary | Adjusted P Value |
|---|---|---|
| +/+ vs. +/DG | * | 0.0289 |
| +/+ vs. DG/DG | **** | <0.0001 |
| +/DG vs. DG/DG | **** | <0.0001 |

| **Shortest MT** | Summary | Adjusted P Value |
|---|---|---|
| +/+ vs. +/DG | ** | 0.0031 |
| +/+ vs. DG/DG | **** | <0.0001 |
| +/DG vs. DG/DG | **** | <0.0001 |

#### FIGURE S5B

Statistics calculated for centriole width and length analysis in RPE1
Unpaired t test with Welch's correction

| **Proximal Width** | Summary | Adjusted P Value |
|---|---|---|
| +/+ vs. DG/DG | * | 0.0284 |

| **Distal Width** | Summary | Adjusted P Value |
|---|---|---|
| +/+ vs. DG/DG | ns | 0.8791 |

| **Longest MT** | Summary | Adjusted P Value |
|---|---|---|
| +/+ vs. DG/DG | **** | <0.0001 |

| **Shortest MT** | Summary | Adjusted P Value |
|---|---|---|
| +/+ vs. DG/DG | **** | <0.0001 |

#### FIGURE S5C

Statistics calculated for centriole foci number analysis in RPE1
two-way anova with Tukey's multiple comparisons test

**CEP135**

| **0** | Summary | Adjusted P Value |
|---|---|---|
| +/+ vs. DG #1 | ns | 0.7627 |
| +/+ vs. DG #2 | ns | 0.9215 |
| DG #1 vs. DG #2 | ns | 0.9463 |
| **1** | | |
| +/+ vs. DG #1 | ns | 0.9639 |
| +/+ vs. DG #2 | ns | 0.465 |
| DG #1 vs. DG #2 | ns | 0.6255 |
| **2** | | |
| +/+ vs. DG #1 | **** | <0.0001 |
| +/+ vs. DG #2 | **** | <0.0001 |
| DG #1 vs. DG #2 | * | 0.0478 |
| **3** | | |
| +/+ vs. DG #1 | **** | <0.0001 |
| +/+ vs. DG #2 | **** | <0.0001 |
| DG #1 vs. DG #2 | ns | 0.6479 |
| **4** | | |
| +/+ vs. DG #1 | **** | <0.0001 |
| +/+ vs. DG #2 | **** | <0.0001 |
| DG #1 vs. DG #2 | ns | 0.9837 |
| **>4** | | |
| +/+ vs. DG #1 | ns | 0.1657 |
| +/+ vs. DG #2 | ns | 0.1937 |
| DG #1 vs. DG #2 | ns | 0.9963 |

### FIGURE S5E

Statistics calculated for centriole foci number analysis in RPE1
two-way anova with Sidak's multiple comparisons test

**SAS-6**

| +/+ vs. -/- | Summary | Adjusted P Value |
|---|---|---|
| 0 | ** | 0.0016 |
| 1 | ns | 0.8495 |
| 2 | **** | <0.0001 |
| 3 | ns | >0.9999 |
| 4 | ns | >0.9999 |
| >4 | ns | >0.9999 |

**CEP135**

| +/+ vs. -/- | | |
|---|---|---|
| 0 | ns | 0.7469 |
| 1 | ns | 0.9022 |
| 2 | **** | <0.0001 |
| 3 | **** | <0.0001 |
| 4 | * | 0.0363 |
| >4 | ns | 0.9977 |

**CPAP**

| +/+ vs. -/- | | |
|---|---|---|
| 0 | ns | 0.9989 |
| 1 | ns | 0.7565 |
| 2 | *** | 0.0007 |
| 3 | ns | 0.2939 |
| 4 | ns | 0.9999 |
| >4 | ns | 0.7983 |

**CEP295**

| +/+ vs. -/- | | |
|---|---|---|
| 0 | ns | >0.9999 |
| 1 | ns | 0.2364 |
| 2 | ns | 0.3677 |
| 3 | ns | 0.672 |
| 4 | ns | 0.3483 |
| >4 | ns | >0.9999 |

**CEP120**

| +/+ vs. -/- | | |
|---|---|---|
| 0 | ns | 0.9788 |
| 1 | ns | 0.9877 |
| 2 | ns | 0.9944 |
| 3 | ns | 0.7117 |
| 4 | ns | 0.8186 |
| >4 | ns | 0.9998 |

**CP110**

| +/+ vs. -/- | | |
|---|---|---|
| 0 | ns | >0.9999 |
| 1 | ns | >0.9999 |
| 2 | ns | 0.1277 |
| 3 | ** | 0.0086 |
| 4 | ns | 0.4441 |
| >4 | ns | 0.9914 |

**POC5**

| +/+ vs. -/- | | |
|---|---|---|
| 0 | * | 0.0484 |
| 1 | **** | <0.0001 |
| 2 | **** | <0.0001 |
| 3 | ns | 0.9813 |
| 4 | ns | 0.9909 |
| >4 | ns | >0.9999 |

**Centrin2**

| +/+ vs. -/- | | |
|---|---|---|
| 0 | *** | 0.0006 |
| 1 | **** | <0.0001 |
| 2 | **** | <0.0001 |
| 3 | ns | 0.8725 |
| 4 | **** | <0.0001 |
| >4 | ns | 0.9973 |

**C2CD3**

| +/+ vs. -/- | | |
|---|---|---|
| 0 | **** | <0.0001 |
| 1 | **** | <0.0001 |
| 2 | **** | <0.0001 |
| 3 | ns | 0.9996 |
| 4 | **** | <0.0001 |
| >4 | ns | >0.9999 |

**TALPID3**

| +/+ vs. -/- | | |
|---|---|---|
| 0 | **** | <0.0001 |
| 1 | ** | 0.0073 |
| 2 | **** | <0.0001 |
| 3 | ns | 0.981 |
| 4 | *** | 0.0005 |
| >4 | ns | >0.9999 |

**CEP164**

| +/+ vs. -/- | | |
|---|---|---|
| 0 | **** | <0.0001 |
| 1 | **** | <0.0001 |
| 2 | ns | 0.997 |
| 3 | ns | >0.9999 |
| 4 | ns | >0.9999 |
| >4 | ns | >0.9999 |

#### FIGURE S6A

Statistics calculated for foci number analysis in DLD1
two-way anova with Sidak's multiple comparisons test

| **CPAP** | Šídák's multiple comparisons test | |
|---|---|---|
| +/+ vs. -/- | | |
| 0 | ns | 0.8037 |
| 1 | ns | 0.9972 |
| 2 | ns | 0.0834 |
| 3 | ns | 0.9831 |
| 4 | ns | 0.9999 |
| >4 | ns | >0.9999 |

| **POC5** | | |
|---|---|---|
| +/+ vs. -/- | Summary | Adjusted P Value |
| 0 | No | ns |
| 1 | No | ns |
| 2 | Yes | *** |
| 3 | No | ns |
| 4 | No | ns |
| >4 | No | ns |

| **CEP295** | | |
|---|---|---|
| +/+ vs. -/- | | |
| 0 | ns | 0.997 |
| 1 | ns | 0.9861 |
| 2 | ns | 0.184 |
| 3 | ns | >0.9999 |
| 4 | ns | 0.9961 |
| >4 | ns | 0.2788 |

| **TALPID3** | | |
|---|---|---|
| +/+ vs. -/- | | |
| 0 | Yes | **** |
| 1 | Yes | ** |
| 2 | Yes | **** |
| 3 | No | ns |
| 4 | Yes | *** |
| >4 | No | ns |

| **CEP120** | | |
|---|---|---|
| +/+ vs. -/- | | |
| 0 | ns | 0.9451 |
| 1 | * | 0.0358 |
| 2 | ns | 0.1282 |
| 3 | ns | 0.3901 |
| 4 | ns | 0.7235 |
| >4 | ns | 0.6925 |

| **Ankrd26** | | |
|---|---|---|
| +/+ vs. -/- | | |
| 0 | Yes | **** |
| 1 | Yes | **** |
| 2 | No | ns |
| 3 | No | ns |
| 4 | No | ns |
| >4 | No | ns |

#### FIGURE S6B

Statistics calculated for CEP135 foci number analysis in DLD1
two-way anova with Tukey's multiple comparisons test

| **CEP135** | | |
|---|---|---|
| **0** | Summary | Adjusted P Value |
| +/+ vs. -/- | ns | 0.4967 |
| +/+ vs. -/- WT dox2d | ns | 0.9997 |
| +/+ vs. -/- WT dox4d | ns | 0.9162 |
| +/+ vs. -/- DG dox2d | ns | 0.9998 |
| +/+ vs. -/- DG dox4d | ns | >0.9999 |
| -/- vs. -/- WT dox2d | ns | 0.8226 |
| -/- vs. -/- WT dox4d | ns | 0.9743 |
| -/- vs. -/- DG dox2d | ns | 0.4793 |
| -/- vs. -/- DG dox4d | ns | 0.5155 |
| -/- WT dox2d vs. -/- WT dox4d | ns | 0.9926 |
| -/- WT dox2d vs. -/- DG dox2d | ns | 0.9961 |
| -/- WT dox2d vs. -/- DG dox4d | ns | 0.9998 |
| -/- WT dox4d vs. -/- DG dox2d | ns | 0.8675 |
| -/- WT dox4d vs. -/- DG dox4d | ns | 0.9254 |
| -/- DG dox2d vs. -/- DG dox4d | ns | 0.9997 |
| **1** | | |
| +/+ vs. -/- | ns | 0.9998 |
| +/+ vs. -/- WT dox2d | ns | 0.9983 |
| +/+ vs. -/- WT dox4d | ns | 0.9703 |
| +/+ vs. -/- DG dox2d | ns | 0.9999 |
| +/+ vs. -/- DG dox4d | ns | >0.9999 |
| -/- vs. -/- WT dox2d | ns | >0.9999 |
| -/- vs. -/- WT dox4d | ns | 0.9955 |
| -/- vs. -/- DG dox2d | ns | >0.9999 |
| -/- vs. -/- DG dox4d | ns | >0.9999 |
| -/- WT dox2d vs. -/- WT dox4d | ns | 0.9998 |
| -/- WT dox2d vs. -/- DG dox2d | ns | >0.9999 |
| -/- WT dox2d vs. -/- DG dox4d | ns | 0.9998 |
| -/- WT dox4d vs. -/- DG dox2d | ns | 0.9981 |
| -/- WT dox4d vs. -/- DG dox4d | ns | 0.9901 |
| -/- DG dox2d vs. -/- DG dox4d | ns | >0.9999 |
| **2** | | |
| +/+ vs. -/- | **** | <0.0001 |
| +/+ vs. -/- WT dox2d | ns | 0.0667 |
| +/+ vs. -/- WT dox4d | ** | 0.0057 |
| +/+ vs. -/- DG dox2d | ** | 0.0046 |
| +/+ vs. -/- DG dox4d | ** | 0.0016 |
| -/- vs. -/- WT dox2d | ns | 0.1914 |
| -/- vs. -/- WT dox4d | ns | 0.2071 |
| -/- vs. -/- DG dox2d | ns | 0.7093 |
| -/- vs. -/- DG dox4d | ns | 0.3956 |
| -/- WT dox2d vs. -/- WT dox4d | ns | 0.9993 |
| -/- WT dox2d vs. -/- DG dox2d | ns | 0.9664 |
| -/- WT dox2d vs. -/- DG dox4d | ns | 0.9854 |
| -/- WT dox4d vs. -/- DG dox2d | ns | 0.9938 |
| -/- WT dox4d vs. -/- DG dox4d | ns | 0.9991 |
| -/- DG dox2d vs. -/- DG dox4d | ns | >0.9999 |
| **3** | | |
| +/+ vs. -/- | ns | 0.4923 |
| +/+ vs. -/- WT dox2d | ns | 0.6342 |
| +/+ vs. -/- WT dox4d | ns | 0.8094 |
| +/+ vs. -/- DG dox2d | ns | 0.8135 |
| +/+ vs. -/- DG dox4d | ns | 0.4749 |
| -/- vs. -/- WT dox2d | ns | >0.9999 |
| -/- vs. -/- WT dox4d | ns | 0.9957 |
| -/- vs. -/- DG dox2d | ns | 0.9997 |
| -/- vs. -/- DG dox4d | ns | >0.9999 |
| -/- WT dox2d vs. -/- WT dox4d | ns | 0.9973 |
| -/- WT dox2d vs. -/- DG dox2d | ns | 0.9998 |
| -/- WT dox2d vs. -/- DG dox4d | ns | >0.9999 |
| -/- WT dox4d vs. -/- DG dox2d | ns | >0.9999 |
| -/- WT dox4d vs. -/- DG dox4d | ns | 0.9945 |
| -/- DG dox2d vs. -/- DG dox4d | ns | 0.9996 |
| **4** | | |
| +/+ vs. -/- | ns | 0.9909 |
| +/+ vs. -/- WT dox2d | ns | 0.9729 |
| +/+ vs. -/- WT dox4d | ns | 0.6904 |
| +/+ vs. -/- DG dox2d | ns | 0.5733 |
| +/+ vs. -/- DG dox4d | ns | 0.891 |
| -/- vs. -/- WT dox2d | ns | 0.9999 |
| -/- vs. -/- WT dox4d | ns | 0.9552 |
| -/- vs. -/- DG dox2d | ns | 0.8716 |
| -/- vs. -/- DG dox4d | ns | 0.997 |
| -/- WT dox2d vs. -/- WT dox4d | ns | 0.9961 |
| -/- WT dox2d vs. -/- DG dox2d | ns | 0.9697 |
| -/- WT dox2d vs. -/- DG dox4d | ns | >0.9999 |
| -/- WT dox4d vs. -/- DG dox2d | ns | 0.999 |
| -/- WT dox4d vs. -/- DG dox4d | ns | 0.9989 |
| -/- DG dox2d vs. -/- DG dox4d | ns | 0.9802 |
| **>4** | | |
| +/+ vs. -/- | ns | 0.3215 |
| +/+ vs. -/- WT dox2d | ns | 0.9909 |
| +/+ vs. -/- WT dox4d | ns | 0.9837 |
| +/+ vs. -/- DG dox2d | ns | 0.7824 |
| +/+ vs. -/- DG dox4d | ns | 0.8053 |
| -/- vs. -/- WT dox2d | ns | 0.8476 |
| -/- vs. -/- WT dox4d | ns | 0.7463 |
| -/- vs. -/- DG dox2d | ns | 0.9971 |
| -/- vs. -/- DG dox4d | ns | 0.9692 |
| -/- WT dox2d vs. -/- WT dox4d | ns | >0.9999 |
| -/- WT dox2d vs. -/- DG dox2d | ns | 0.9885 |
| -/- WT dox2d vs. -/- DG dox4d | ns | 0.9963 |
| -/- WT dox4d vs. -/- DG dox2d | ns | 0.9799 |
| -/- WT dox4d vs. -/- DG dox4d | ns | 0.9922 |
| -/- DG dox2d vs. -/- DG dox4d | ns | >0.9999 |

#### FIGURE S6C

Statistics calculated for centriole microtubule length analysis in DLD1
one-way anova with Tukey's multiple comparisons test

| **Longest MT** | Summary | Adjusted P Value |
|---|---|---|
| +/+ vs. -/- | *** | 0.0008 |
| +/+ vs. -/- WT dox2d | ns | 0.9835 |
| +/+ vs. -/- WT dox4d | ns | 0.9431 |
| +/+ vs. -/- DG dox2d | * | 0.0179 |
| +/+ vs. -/- DG dox4d | ** | 0.0017 |
| -/- vs. -/- WT dox2d | ** | 0.0057 |
| -/- vs. -/- WT dox4d | **** | <0.0001 |
| -/- vs. -/- DG dox2d | ns | 0.889 |
| -/- vs. -/- DG dox4d | ns | 0.9041 |
| -/- WT dox2d vs. -/- WT dox4d | ns | 0.4564 |
| -/- WT dox2d vs. -/- DG dox2d | ns | 0.0948 |
| -/- WT dox2d vs. -/- DG dox4d | * | 0.0159 |
| -/- WT dox4d vs. -/- DG dox2d | **** | <0.0001 |
| -/- WT dox4d vs. -/- DG dox4d | **** | <0.0001 |
| -/- DG dox2d vs. -/- DG dox4d | ns | >0.9999 |

| **Shortest MT** | Summary | Adjusted P Value |
|---|---|---|
| +/+ vs. -/- | **** | <0.0001 |
| +/+ vs. -/- WT dox2d | ns | 0.1282 |
| +/+ vs. -/- WT dox4d | ns | 0.9876 |
| +/+ vs. -/- DG dox2d | **** | <0.0001 |
| +/+ vs. -/- DG dox4d | **** | <0.0001 |
| -/- vs. -/- WT dox2d | * | 0.0347 |
| -/- vs. -/- WT dox4d | **** | <0.0001 |
| -/- vs. -/- DG dox2d | ns | 0.999 |
| -/- vs. -/- DG dox4d | ns | 0.6772 |
| -/- WT dox2d vs. -/- WT dox4d | ns | 0.158 |
| -/- WT dox2d vs. -/- DG dox2d | ns | 0.0501 |
| -/- WT dox2d vs. -/- DG dox4d | **** | <0.0001 |
| -/- WT dox4d vs. -/- DG dox2d | **** | <0.0001 |
| -/- WT dox4d vs. -/- DG dox4d | **** | <0.0001 |
| -/- DG dox2d vs. -/- DG dox4d | ns | 0.2643 |

#### FIGURE S6E

Statistics calculated for centriole defects analysis upon Taxol treatment
two-way anova with Tukey's multiple comparisons test

| **normal** | Summary | Adjusted P Value |
|---|---|---|
| +/+ DMSO vs. +/+ Taxol | ns | 0.9933 |
| +/+ DMSO vs. -/- DMSO | **** | <0.0001 |
| +/+ DMSO vs. -/- Taxol | * | 0.0445 |
| +/+ Taxol vs. -/- DMSO | **** | <0.0001 |
| +/+ Taxol vs. -/- Taxol | ns | 0.0735 |
| -/- DMSO vs. -/- Taxol | *** | 0.0007 |
| **short** | | |
| +/+ DMSO vs. +/+ Taxol | ns | 0.8082 |
| +/+ DMSO vs. -/- DMSO | * | 0.0299 |
| +/+ DMSO vs. -/- Taxol | ns | 0.4622 |
| +/+ Taxol vs. -/- DMSO | ns | 0.1569 |
| +/+ Taxol vs. -/- Taxol | ns | 0.9296 |
| -/- DMSO vs. -/- Taxol | ns | 0.3915 |
| **broken** | | |
| +/+ DMSO vs. +/+ Taxol | ns | 0.7921 |
| +/+ DMSO vs. -/- DMSO | *** | 0.0007 |
| +/+ DMSO vs. -/- Taxol | ns | 0.3767 |
| +/+ Taxol vs. -/- DMSO | *** | 0.0001 |
| +/+ Taxol vs. -/- Taxol | ns | 0.0844 |
| -/- DMSO vs. -/- Taxol | * | 0.0194 |
| **long** | | |
| +/+ DMSO vs. +/+ Taxol | ns | 0.9906 |
| +/+ DMSO vs. -/- DMSO | ns | 0.9954 |
| +/+ DMSO vs. -/- Taxol | ns | 0.9954 |
| +/+ Taxol vs. -/- DMSO | ns | 0.951 |
| +/+ Taxol vs. -/- Taxol | ns | 0.951 |
| -/- DMSO vs. -/- Taxol | ns | >0.9999 |

Statistics calculated for longest MT and Shortest MT analysis upon Taxol treatment
one-way anova with Tukey's multiple comparisons test

| Ratio | Summary | Adjusted P Value |
|---|---|---|
| +/+ DMSO vs. +/+ Taxol | ns | 0.9788 |
| +/+ DMSO vs. -/- DMSO | ** | 0.0032 |
| +/+ DMSO vs. -/- Taxol | ns | 0.5236 |
| +/+ Taxol vs. -/- DMSO | ** | 0.0012 |
| +/+ Taxol vs. -/- Taxol | ns | 0.3077 |
| -/- DMSO vs. -/- Taxol | ns | 0.0964 |

| Longest MT | Summary | Adjusted P Value |
|---|---|---|
| +/+ DMSO vs. +/+ Taxol | ns | 0.9945 |
| +/+ DMSO vs. -/- DMSO | **** | <0.0001 |
| +/+ DMSO vs. -/- Taxol | ns | 0.1892 |
| +/+ Taxol vs. -/- DMSO | **** | <0.0001 |
| +/+ Taxol vs. -/- Taxol | ns | 0.3446 |
| -/- DMSO vs. -/- Taxol | **** | <0.0001 |

| Shortest MT | Summary | Adjusted P Value |
|---|---|---|
| +/+ DMSO vs. +/+ Taxol | ns | 0.9855 |
| +/+ DMSO vs. -/- DMSO | **** | <0.0001 |
| +/+ DMSO vs. -/- Taxol | ns | 0.0552 |
| +/+ Taxol vs. -/- DMSO | **** | <0.0001 |
| +/+ Taxol vs. -/- Taxol | * | 0.0266 |
| -/- DMSO vs. -/- Taxol | **** | <0.0001 |

#### FIGURE S6G

Statistics calculated for centriole markers intensity analysis upon Taxol treatment
one-way anova with Tukey's multiple comparisons test

| **CEP135** | Summary | Adjusted P Value |
|---|---|---|
| +/+ DMSO vs. +/+ Taxol | * | 0.0217 |
| +/+ DMSO vs. -/- DMSO | ** | 0.0026 |
| +/+ DMSO vs. -/- Taxol | ** | 0.0015 |
| +/+ Taxol vs. -/- DMSO | ns | 0.3932 |
| +/+ Taxol vs. -/- Taxol | ns | 0.2087 |
| -/- DMSO vs. -/- Taxol | ns | 0.9566 |

| **Centrin2** | Summary | Adjusted P Value |
|---|---|---|
| +/+ DMSO vs. +/+ Taxol | ns | 0.1776 |
| +/+ DMSO vs. -/- DMSO | * | 0.0107 |
| +/+ DMSO vs. -/- Taxol | * | 0.0262 |
| +/+ Taxol vs. -/- DMSO | ns | 0.2524 |
| +/+ Taxol vs. -/- Taxol | ns | 0.5494 |
| -/- DMSO vs. -/- Taxol | ns | 0.9073 |

| **talpid3** | Summary | Adjusted P Value |
|---|---|---|
| +/+ DMSO vs. +/+ Taxol | ns | 0.3761 |
| +/+ DMSO vs. -/- DMSO | * | 0.0233 |
| +/+ DMSO vs. -/- Taxol | ns | 0.0754 |
| +/+ Taxol vs. -/- DMSO | ns | 0.2516 |
| +/+ Taxol vs. -/- Taxol | ns | 0.6461 |
| -/- DMSO vs. -/- Taxol | ns | 0.8336 |

| **CEP164** | Summary | Adjusted P Value |
|---|---|---|
| +/+ DMSO vs. +/+ Taxol | ns | 0.3483 |
| +/+ DMSO vs. -/- DMSO | *** | 0.0002 |
| +/+ DMSO vs. -/- Taxol | *** | 0.0005 |
| +/+ Taxol vs. -/- DMSO | *** | 0.0008 |
| +/+ Taxol vs. -/- Taxol | ** | 0.0036 |
| -/- DMSO vs. -/- Taxol | ns | 0.5782 |

### FIGURE S7D

Statistics calculated for centriole markers intensity upon CEP120 siRNA
one-way anova with Tukey's multiple comparisons test

| **CEP120** | Summary | Adjusted P Value |
|---|---|---|
| -/- WT no siRNA vs. -/- WT CEP120 siRNA | **** | <0.0001 |
| -/- WT no siRNA vs. -/- DG no siRNA | ns | 0.9999 |
| -/- WT no siRNA vs. -/- DG CEP120 siRNA | **** | <0.0001 |
| -/- WT CEP120 siRNA vs. -/- DG no siRNA | **** | <0.0001 |
| -/- WT CEP120 siRNA vs. -/- DG CEP120 siRNA | ns | 0.8679 |
| -/- DG no siRNA vs. -/- DG CEP120 siRNA | **** | <0.0001 |

| **cep135** | Summary | Adjusted P Value |
|---|---|---|
| -/- WT no siRNA vs. -/- WT CEP120 siRNA | * | 0.0457 |
| -/- WT no siRNA vs. -/- DG no siRNA | ns | 0.8035 |
| -/- WT no siRNA vs. -/- DG CEP120 siRNA | *** | 0.0007 |
| -/- WT CEP120 siRNA vs. -/- DG no siRNA | ns | 0.1625 |
| -/- WT CEP120 siRNA vs. -/- DG CEP120 siRNA | * | 0.0351 |
| -/- DG no siRNA vs. -/- DG CEP120 siRNA | ** | 0.0018 |

#### FIGURE S7E

Statistics calculated for centriole foci number upon CEP120 siRNA
two-way anova with Tukey's multiple comparisons test

| HYLS1 | Summary | Adjusted P Value |
|---|---|---|
| **0** | | |
| -/- WT no siRNA vs. -/- WT CEP120 siRNA | *** | 0.0002 |
| -/- WT no siRNA vs. -/- DG no siRNA | **** | <0.0001 |
| -/- WT no siRNA vs. -/- DG CEP120 siRNA | **** | <0.0001 |
| -/- WT CEP120 siRNA vs. -/- DG no siRNA | ns | 0.0943 |
| -/- WT CEP120 siRNA vs. -/- DG CEP120 siRNA | **** | <0.0001 |
| -/- DG no siRNA vs. -/- DG CEP120 siRNA | ** | 0.001 |
| **1** | | |
| -/- WT no siRNA vs. -/- WT CEP120 siRNA | ns | 0.4327 |
| -/- WT no siRNA vs. -/- DG no siRNA | ns | 0.9096 |
| -/- WT no siRNA vs. -/- DG CEP120 siRNA | ns | 0.5846 |
| -/- WT CEP120 siRNA vs. -/- DG no siRNA | ns | 0.152 |
| -/- WT CEP120 siRNA vs. -/- DG CEP120 siRNA | ns | 0.9939 |
| -/- DG no siRNA vs. -/- DG CEP120 siRNA | ns | 0.2375 |
| **2** | | |
| -/- WT no siRNA vs. -/- WT CEP120 siRNA | ** | 0.0099 |
| -/- WT no siRNA vs. -/- DG no siRNA | **** | <0.0001 |
| -/- WT no siRNA vs. -/- DG CEP120 siRNA | **** | <0.0001 |
| -/- WT CEP120 siRNA vs. -/- DG no siRNA | ns | 0.1035 |
| -/- WT CEP120 siRNA vs. -/- DG CEP120 siRNA | **** | <0.0001 |
| -/- DG no siRNA vs. -/- DG CEP120 siRNA | * | 0.0179 |
| **3** | | |
| -/- WT no siRNA vs. -/- WT CEP120 siRNA | ns | 0.3465 |
| -/- WT no siRNA vs. -/- DG no siRNA | ns | 0.9092 |
| -/- WT no siRNA vs. -/- DG CEP120 siRNA | ns | 0.1987 |
| -/- WT CEP120 siRNA vs. -/- DG no siRNA | ns | 0.732 |
| -/- WT CEP120 siRNA vs. -/- DG CEP120 siRNA | ns | 0.9848 |
| -/- DG no siRNA vs. -/- DG CEP120 siRNA | ns | 0.5212 |
| **4** | | |
| -/- WT no siRNA vs. -/- WT CEP120 siRNA | ns | 0.7406 |
| -/- WT no siRNA vs. -/- DG no siRNA | ns | 0.9363 |
| -/- WT no siRNA vs. -/- DG CEP120 siRNA | ns | 0.4216 |
| -/- WT CEP120 siRNA vs. -/- DG no siRNA | ns | 0.9717 |
| -/- WT CEP120 siRNA vs. -/- DG CEP120 siRNA | ns | 0.9489 |
| -/- DG no siRNA vs. -/- DG CEP120 siRNA | ns | 0.7657 |
| **>4** | | |
| -/- WT no siRNA vs. -/- WT CEP120 siRNA | ns | 0.9561 |
| -/- WT no siRNA vs. -/- DG no siRNA | ns | 0.9997 |
| -/- WT no siRNA vs. -/- DG CEP120 siRNA | ns | 0.9329 |
| -/- WT CEP120 siRNA vs. -/- DG no siRNA | ns | 0.9318 |
| -/- WT CEP120 siRNA vs. -/- DG CEP120 siRNA | ns | 0.9998 |
| -/- DG no siRNA vs. -/- DG CEP120 siRNA | ns | 0.9027 |

| CEP120 | Summary | Adjusted P Value |
|---|---|---|
| **0** | | |
| -/- WT no siRNA vs. -/- WT CEP120 siRNA | **** | <0.0001 |
| -/- WT no siRNA vs. -/- DG no siRNA | ns | 0.9924 |
| -/- WT no siRNA vs. -/- DG CEP120 siRNA | **** | <0.0001 |
| -/- WT CEP120 siRNA vs. -/- DG no siRNA | **** | <0.0001 |
| -/- WT CEP120 siRNA vs. -/- DG CEP120 siRNA | *** | 0.0004 |
| -/- DG no siRNA vs. -/- DG CEP120 siRNA | **** | <0.0001 |
| **1** | | |
| -/- WT no siRNA vs. -/- WT CEP120 siRNA | ** | 0.008 |
| -/- WT no siRNA vs. -/- DG no siRNA | ns | 0.8467 |
| -/- WT no siRNA vs. -/- DG CEP120 siRNA | ** | 0.0076 |
| -/- WT CEP120 siRNA vs. -/- DG no siRNA | ** | 0.0011 |
| -/- WT CEP120 siRNA vs. -/- DG CEP120 siRNA | ns | >0.9999 |
| -/- DG no siRNA vs. -/- DG CEP120 siRNA | ** | 0.001 |
| **2** | | |
| -/- WT no siRNA vs. -/- WT CEP120 siRNA | **** | <0.0001 |
| -/- WT no siRNA vs. -/- DG no siRNA | * | 0.0287 |
| -/- WT no siRNA vs. -/- DG CEP120 siRNA | **** | <0.0001 |
| -/- WT CEP120 siRNA vs. -/- DG no siRNA | **** | <0.0001 |
| -/- WT CEP120 siRNA vs. -/- DG CEP120 siRNA | ** | 0.002 |
| -/- DG no siRNA vs. -/- DG CEP120 siRNA | **** | <0.0001 |
| **3** | | |
| -/- WT no siRNA vs. -/- WT CEP120 siRNA | ns | 0.1317 |
| -/- WT no siRNA vs. -/- DG no siRNA | ns | 0.067 |
| -/- WT no siRNA vs. -/- DG CEP120 siRNA | * | 0.0485 |
| -/- WT CEP120 siRNA vs. -/- DG no siRNA | *** | 0.0003 |
| -/- WT CEP120 siRNA vs. -/- DG CEP120 siRNA | ns | 0.9592 |
| -/- DG no siRNA vs. -/- DG CEP120 siRNA | **** | <0.0001 |
| **4** | | |
| -/- WT no siRNA vs. -/- WT CEP120 siRNA | ns | 0.6321 |
| -/- WT no siRNA vs. -/- DG no siRNA | ns | 0.9975 |
| -/- WT no siRNA vs. -/- DG CEP120 siRNA | ns | 0.5758 |
| -/- WT CEP120 siRNA vs. -/- DG no siRNA | ns | 0.5175 |
| -/- WT CEP120 siRNA vs. -/- DG CEP120 siRNA | ns | 0.9997 |
| -/- DG no siRNA vs. -/- DG CEP120 siRNA | ns | 0.4632 |
| **>4** | | |
| -/- WT no siRNA vs. -/- WT CEP120 siRNA | ns | 0.9866 |
| -/- WT no siRNA vs. -/- DG no siRNA | ns | 0.6296 |
| -/- WT no siRNA vs. -/- DG CEP120 siRNA | ns | 0.9666 |
| -/- WT CEP120 siRNA vs. -/- DG no siRNA | ns | 0.43 |
| -/- WT CEP120 siRNA vs. -/- DG CEP120 siRNA | ns | 0.9993 |
| -/- DG no siRNA vs. -/- DG CEP120 siRNA | ns | 0.3622 |

| CEP135 | Summary | Adjusted P Value |
|---|---|---|
| **0** | | |
| -/- WT no siRNA vs. -/- WT CEP120 siRNA | ns | 0.9886 |
| -/- WT no siRNA vs. -/- DG no siRNA | ns | >0.9999 |
| -/- WT no siRNA vs. -/- DG CEP120 siRNA | ns | 0.9729 |
| -/- WT CEP120 siRNA vs. -/- DG no siRNA | ns | 0.9886 |
| -/- WT CEP120 siRNA vs. -/- DG CEP120 siRNA | ns | 0.9995 |
| -/- DG no siRNA vs. -/- DG CEP120 siRNA | ns | 0.9729 |
| **1** | | |
| -/- WT no siRNA vs. -/- WT CEP120 siRNA | ns | 0.9858 |
| -/- WT no siRNA vs. -/- DG no siRNA | ns | 0.9797 |
| -/- WT no siRNA vs. -/- DG CEP120 siRNA | ns | 0.8521 |
| -/- WT CEP120 siRNA vs. -/- DG no siRNA | ns | 0.8837 |
| -/- WT CEP120 siRNA vs. -/- DG CEP120 siRNA | ns | 0.9668 |
| -/- DG no siRNA vs. -/- DG CEP120 siRNA | ns | 0.6378 |
| **2** | | |
| -/- WT no siRNA vs. -/- WT CEP120 siRNA | ** | 0.01 |
| -/- WT no siRNA vs. -/- DG no siRNA | ns | 0.1772 |
| -/- WT no siRNA vs. -/- DG CEP120 siRNA | ** | 0.0011 |
| -/- WT CEP120 siRNA vs. -/- DG no siRNA | ns | 0.537 |
| -/- WT CEP120 siRNA vs. -/- DG CEP120 siRNA | ns | 0.813 |
| -/- DG no siRNA vs. -/- DG CEP120 siRNA | ns | 0.1408 |
| **3** | | |
| -/- WT no siRNA vs. -/- WT CEP120 siRNA | ns | 0.2083 |
| -/- WT no siRNA vs. -/- DG no siRNA | ns | 0.9067 |
| -/- WT no siRNA vs. -/- DG CEP120 siRNA | ns | 0.5473 |
| -/- WT CEP120 siRNA vs. -/- DG no siRNA | ns | 0.5418 |
| -/- WT CEP120 siRNA vs. -/- DG CEP120 siRNA | ns | 0.9033 |
| -/- DG no siRNA vs. -/- DG CEP120 siRNA | ns | 0.9079 |
| **4** | | |
| -/- WT no siRNA vs. -/- WT CEP120 siRNA | ns | 0.9731 |
| -/- WT no siRNA vs. -/- DG no siRNA | ns | 0.5316 |
| -/- WT no siRNA vs. -/- DG CEP120 siRNA | ns | 0.5901 |
| -/- WT CEP120 siRNA vs. -/- DG no siRNA | ns | 0.7846 |
| -/- WT CEP120 siRNA vs. -/- DG CEP120 siRNA | ns | 0.8339 |
| -/- DG no siRNA vs. -/- DG CEP120 siRNA | ns | 0.9997 |
| **>4** | | |
| -/- WT no siRNA vs. -/- WT CEP120 siRNA | ns | 0.9832 |
| -/- WT no siRNA vs. -/- DG no siRNA | ns | 0.98 |
| -/- WT no siRNA vs. -/- DG CEP120 siRNA | ns | 0.9549 |
| -/- WT CEP120 siRNA vs. -/- DG no siRNA | ns | >0.9999 |
| -/- WT CEP120 siRNA vs. -/- DG CEP120 siRNA | ns | 0.9987 |
| -/- DG no siRNA vs. -/- DG CEP120 siRNA | ns | 0.9992 |
